# Chromosomal inversions and the demography of speciation in *Drosophila montana* and *Drosophila flavomontana*

**DOI:** 10.1101/2022.11.15.516589

**Authors:** N. Poikela, D. R. Laetsch, V. Hoikkala, K. Lohse, M Kankare

## Abstract

Chromosomal inversions may play a central role in speciation given their ability to locally reduce recombination and therefore genetic exchange between diverging populations. We analysed long- and short-read whole-genome data from sympatric and allopatric populations of two *Drosophila virilis* group species, *D. montana* and *D. flavomontana*, to understand if inversions have contributed to their divergence. We identified three large alternatively fixed inversions on the X chromosome and one on each of the autosomes 4 and 5. A comparison of demographic models estimated for inverted and non-inverted (colinear) chromosomal regions suggest that these inversions arose before the time of the species split. We detected a low rate of interspecific gene flow (introgression) from *D. montana* to *D. flavomontana*, which was further reduced inside inversions and was lower in allopatric than in sympatric populations. Together, these results suggest that the inversions were already present in the common ancestral population, and that gene exchange between the sister taxa was reduced within inversions both before and after the onset of species divergence. Such ancestrally polymorphic inversions may foster speciation by allowing the accumulation of genetic divergence in loci involved in adaptation and reproductive isolation inside inversions early in the speciation process, while gene exchange at colinear regions continues until the evolving reproductive barriers complete speciation. The overlapping X inversions are particularly good candidates for driving the speciation process of *D. montana* and *D. flavomontana*, since they harbour strong genetic incompatibilities that were detected in a recent study of experimental introgression.

**Significance:** Chromosomal inversions, genomic rearrangements with reversed gene order, have been extensively studied, but it remains unclear whether and how inversions play a role in species divergence. Analysis of long- and short-read whole-genome data for two *Drosophila* sister species, *D. montana* and *D. flavomontana*, revealed five alternatively fixed inversions. Modelling the demographic history of these inversions shows that they were segregating already in the common ancestor of the species and that they have reduced gene exchange between these sister taxa both before and after the onset of species divergence. These results are compatible with a scenario in which ancestrally polymorphic inversions aid species divergence by protecting divergently selected loci from erosion via gene flow during the earliest stages of speciation.

## Introduction

Chromosomal inversions, genomic regions with reversed gene order, may facilitate adaptation and speciation in the face of gene flow because they suppress recombination between alternate rearrangements, which creates and preserves associations between sets of alleles conferring local adaptation, mate choice and genetic incompatibilities (Sturtevant 1921; Butlin 2005; Hoffmann and Rieseberg 2008). While inversions have been found in many species of insects, fish, birds, mammals and plants, their frequency varies widely between and even within taxa (Stone et al. 1960; Wellenreuther & Bernatchez 2018) and it remains an open question whether and how inversions contribute to the evolution of species divergence. Genomic data from young species pairs offer the chance to reconstruct both the demographic history of species divergence in the face of gene flow and the history of alternatively fixed inversions and interspecific gene flow (introgression) (Faria et al. 2018; Faria & Navarro 2010).

Inversions may facilitate adaptation and speciation in many ways (reviewed in Hoffmann and Rieseberg 2008; Jackson 2011; Faria et al. 2018). A new inversion may be favoured by selection if it protects epistatic interactions (Hoffmann & Rieseberg 2008) and/or locally adapted alleles (Kirkpatrick & Barton 2006) from recombination with immigrant alleles that reside in an alternate rearrangement. Also, an inversion may be under selection if its breakpoints disrupt reading frames of genes, or change the expression of genes (Wright & Schaeffer 2022; Matzkin et al. 2005; Villoutreix et al. 2021). While the probability of fixation of an inversion between diverging populations depends on the strength of selection and the levels of gene flow (Hoffmann & Rieseberg 2008), its potential to contribute to local adaptation and/or speciation in the long term depends also on whether populations evolve in isolation or in the face of gene flow. Upon secondary contact, alternatively fixed inversions may protect existing incompatibilities from gene flow between diverging populations, while non-inverted (colinear) regions are more susceptible to the homogenizing effects of gene flow (Noor et al. 2001). In contrast, if populations diverge in the presence of gene flow, we expect incompatibilities to accumulate in inverted regions (Navarro & Barton 2003). In both scenarios, inversions harbouring incompatibilities delay species’ fusion and provide time for additional barriers to evolve. For example, prezygotic reproductive barriers are expected to be more easily reinforced in response to genetic incompatibilities and maladaptive hybridization (reinforcement) (Servedio & Noor 2003), if the causal loci are located within inversions (Trickett & Butlin 1994; Butlin 2005; Dagilis & Kirkpatrick 2016). Two kinds of empirical observations give indirect support for these theories. First, genes maintaining local adaptation, premating barriers and genetic incompatibilities between species have been found to be concentrated in alternatively fixed inversions (Fishman et al. 2013; Lowry & Willis 2010; Noor et al. 2001). Second, fixed inversions generally have elevated genetic divergence compared to colinear regions (Noor et al. 2007; Kulathinal et al. 2009; Lohse et al. 2015). However, it has proven extremely difficult to distinguish speciation histories in which inversions have acted as triggers of speciation from scenarios in which alternately fixed inversions arise incidentally either because they are polymorphic in the ancestral population for reasons that may have nothing to do with local adaptation (Fuller et al. 2019; Faria & Navarro 2010; Guerrero & Hahn 2017) or because they arise after speciation is complete.

So far only a few studies have dissected the evolutionary history of inversions to explore their role in adaptation (Lundberg et al. 2023) and speciation (e.g. Lohse et al. 2015; Fuller et al. 2018). Demographic models can be used to systematically compare the species’ divergence time estimated from colinear regions (*T_col_*) and the origin of inversions (*T_inv_*) and as well as the amount of long-term effective introgression between inverted (*M_inv_*) and colinear (*M_col_*) regions (Noor and Bennett 2009). Similarly, recent or ongoing introgression can be diagnosed by comparing estimates of *M* between sympatric and allopatric population pairs (Noor and Bennett 2009). There are at least three scenarios for the evolutionary history of alternately fixed inversions. First, inversions arise and fix after speciation is largely complete, most likely for reasons unrelated to the speciation process. In this case, we expect reduced introgression (*M*_inv_ < *M*_col_) within inversions, but the same split time estimates for inversions and colinear regions (*T*_inv_ = *T*_col_). Second, inversions fix during the speciation process because they contribute to local adaptation and/or formation of reproductive isolation at an early stage of high gene flow (Kirkpatrick & Barton 2006). Such inversions should have reduced introgression compared to colinear regions (*M*_inv_ < *M*_col_) and their estimated divergence time either predates that of colinear regions (*T*_inv_ > *T*_col_) (if they have been segregating in the common ancestral population) or is the same (*T*_inv_ = *T*_col_) (if they arose at the onset of divergence). Crucially however, irrespective of their age, we expect that these inversions have fixed because they act as barriers to gene flow, i.e. they protect alleles that are involved in local adaptation, mate choice and/or genetic incompatibilities. Finally, in a third scenario, inversions are segregating in the ancestral population due to forces that have nothing to do with local adaptation or speciation. Importantly, we would expect any inversion that segregates in the ancestral population to be alternately fixed between the two species by chance alone with probability of 1/2 (Guerrero & Hahn 2017). Such coincidental inversions that fix differentially with no effect on species divergence could still help impede species fusion upon secondary contact if they contain BDMIs (Noor et al. 2001). However, the predictions for the coincidental inversion scenario in terms of demographic parameters are the same as in the second scenario above.

The two *Drosophila virilis* group species, *Drosophila montana* and *Drosophila flavomontana*, offer a great opportunity to investigate the potential role of inversions in species divergence. Based on polytene chromosome studies, these species have several alternatively fixed inversions (Throckmorton 1982; Stone et al. 1960), which, however, have so far not been characterised at the genomic level. *D. montana* and *D. flavomontana* have diverged ∼3.8 Mya in the Rocky Mountains, and the two species presently inhabit variable climates in the Rocky Mountains and along the western coast of North America (Hoikkala and Poikela 2022; Yusuf et al. 2022). In the mountains, *D. montana* has spread to higher altitudes than *D. flavomontana*, while on the western coast, where *D. flavomontana* has expanded relatively recently, both species live at low altitudes (Fig. 1, Table S1; Patterson 1952; Hoikkala and Poikela 2022). Thus, in both regions, populations of the two species can be regarded as sympatric or parapatric. However, *D. montana* also has allopatric populations at high latitudes, e.g. in Alaska, where *D. flavomontana* does not exist (Fig. 1, Table S1). Reproductive isolation between *D. montana* females and *D. flavomontana* males is nearly complete, characterized by an extremely strong prezygotic isolation and inviability and sterility of F_1_ females and males (Poikela et al. 2019). In contrast, prezygotic isolation between *D. flavomontana* females and *D. montana* males is relatively weaker, and shows signs of reinforcement in sympatric populations of *D. flavomontana* (Poikela et al. 2019). Furthermore, in these crosses F_1_ hybrid males are sterile but F_1_ hybrid females can be crossed with males of both parental species to obtain backcross progenies in both directions (Poikela et al. 2019, 2022). Importantly, evidence for strong BDMI(s) between these species located within inversions on the X chromosome has been found (Poikela et al. 2022). This prevents introgression from *D. montana* to *D. flavomontana* across the entire X chromosome during early backcross generations (Poikela et al. 2022). Despite the strong reproductive isolation, interspecific hybrids have been found in nature (Patterson 1952; Throckmorton 1982).

Here, we explored whether and how inversions have contributed to the species divergence of *D. montana* and *D. flavomontana*. We used long- and short-read sequencing data from allopatric and sympatric populations of the species to generate highly contiguous assemblies for both species, which in turn enabled us to accurately identify the presence of alternatively fixed inversions. We used demographic modelling to estimate the age of these inversions and their potential effect on the long-term rate of introgression and asked the following specific questions:

1. How many alternatively fixed inversions do *D. montana* and *D. flavomontana* carry?
2. When did these inversions most likely arise and how does their age compare to the species divergence time?
3. Do these inversions show reduced introgression compared to colinear regions as would be expected if they arose during or before the onset of species divergence?

**Figure 1.**
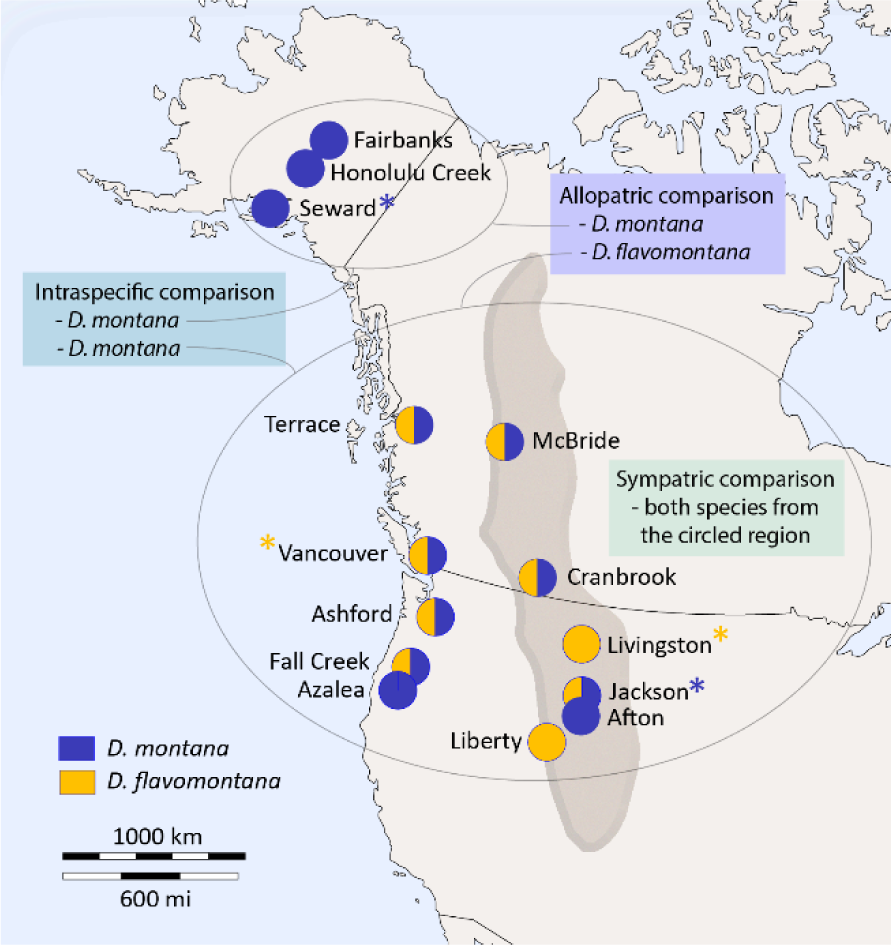
Sampling sites of sympatric (or parapatric) and allopatric *D. montana* and *D. flavomontana* populations in North America. Pie charts indicate the sampling sites for one or both species. Long-read PacBio data were obtained from two isofemale strains per species (sample sites indicated with asterisks). Short-read Illumina data were obtained from single wild-caught females for all sites shown. Gray area illustrates the Rocky Mountains of North America.

## Results and Discussion

We generated long-read PacBio (Pacific Biosciences) sequencing data for females from two *D. montana* and two *D. flavomontana* isofemale strains and short-read Illumina re-sequencing data for 12 *D. montana* and 9 *D. flavomontana* wild-caught females (one female per population per species) originating from allopatric and sympatric populations (Fig. 1, Table S1, S2). These data enabled us to generate contiguous, high-quality genome assemblies for both species to accurately identify alternatively fixed inversions and to examine the species’ evolutionary history and the role of inversions and introgression therein. In the following, we refer to comparison between *D. montana* and *D. flavomontana* samples from the Rocky Mountains and the western coast as “sympatric”, and the comparisons between *D. montana* from Alaska and *D. flavomontana* from the mountains and the coast as “allopatric” (Fig. 1, Table S1). To evaluate the timing of potential recent introgression in sympatry, we estimated the divergence time for *D. montana* living in contact (sympatry) and in isolation (allopatry) with *D. flavomontana*, and we refer to this comparison as “intraspecific” (Fig. 1, Table S1).

### Construction and annotation of species genomes

Two genome assemblies for each species were generated using the PacBio data of two *D. montana* and *D. flavomontana* isofemale strains and the Illumina data for the respective founder females collected from the wild (Table S1 & S2). The assembled genomes had a total length of 181-194Mb (Table S3), which resemble those of previously assembled *D. montana*, *D. flavomontana* and several other *Drosophila* species (128-198Mb) (Miller et al. 2018; Parker et al. 2018; Yusuf et al. 2022). A small proportion of each assembly (0-18 contigs, spanning = 0.0-9.9Mb) was excluded as contaminant sequences, mainly bacteria, based on the coverage, GC% and taxonomic annotation of contigs (Fig. S1-S4). From the 3285 BUSCO groups, we identified 97.3-98.5% as complete BUSCOs, of which 96.9-98.0% were single-copy and 0.4-0.5% duplicated BUSCOs (Table S3). The BUSCO values were similar to the ones in other *Drosophila* assemblies (Miller et al. 2018). Repetitive sequences comprised 25.5-29.9% of our assemblies (Table S3), which is close to the repeat content reported for other *Drosophila* species (e.g. 26.5% in *D. virilis*, 28.6% in *D. melanogaster*, 22.9% in *D. mojavensis*, and 19.9% in *D. elegans*; NCBI Annotation Report). Our annotations included 15,696-16,056 genes per assembly, which is plausible given the number of genes reported for other *Drosophila* assemblies (e.g. Yang et al. 2018). Overall, the combination of long- and short-read data resulted in more contiguous assemblies for both species (N50 values of 1.3-11.0Mb; Table S3) compared to the previously published *D. montana* and *D. flavomontana* genomes which were based on short-read data (e.g. N50 of 41kb in *D. montana;* Parker et al. 2018; Yusuf et al. 2022).

We built a chromosome-level reference genome for *D. montana* by scaffolding with the genome of another *virilis* group species, *Drosophila lummei,* and for *D. flavomontana* by first scaffolding one assembly with the other (within species) and then with the *D. lummei* genome (see Methods for details). For both chromosome-level genomes, the total genome size, BUSCO values and the number of repeats and genes slightly decreased compared to the original, non-scaffolded assemblies (Table S3). Given greater span and completeness (as measured by BUSCO) of the *D. montana* compared to the *D. flavomontana* genome, subsequent analyses were performed using *D. montana* as a reference by default. However, to quantify the effect of reference bias, we repeated the demographic inference using *D. flavomontana* as a reference.

To understand how chromosomes of *D. montana* and *D. flavomontana* relate to the more studied *D. virilis*, we compared the genomes of *D. montana* and *D. flavomontana* (species of the *montana* phylad of the *virilis* group) and *D. virilis* and *D. lummei* (species of the *virilis* phylad of the *virilis* group) (Yusuf et al. 2022). While chromosome synteny is highly variable between distantly related *Drosophila* species, such as *D. melanogaster* and *D. virilis* (Schaeffer et al. 2008), it is relatively similar between the *virilis* group species (Fig. 2; Stone et al. 1960). The most noticeable difference is that in *D. montana* and *D. flavomontana*, chromosome 2 has left (2L) and right (2R) arms that are separated by a (sub)metacentric centromere, while in *D. virilis* and *D. lummei* the centromere is located near one end of the chromosome 2 (Fig. 2; Stone et al. 1960).

**Figure 2.**
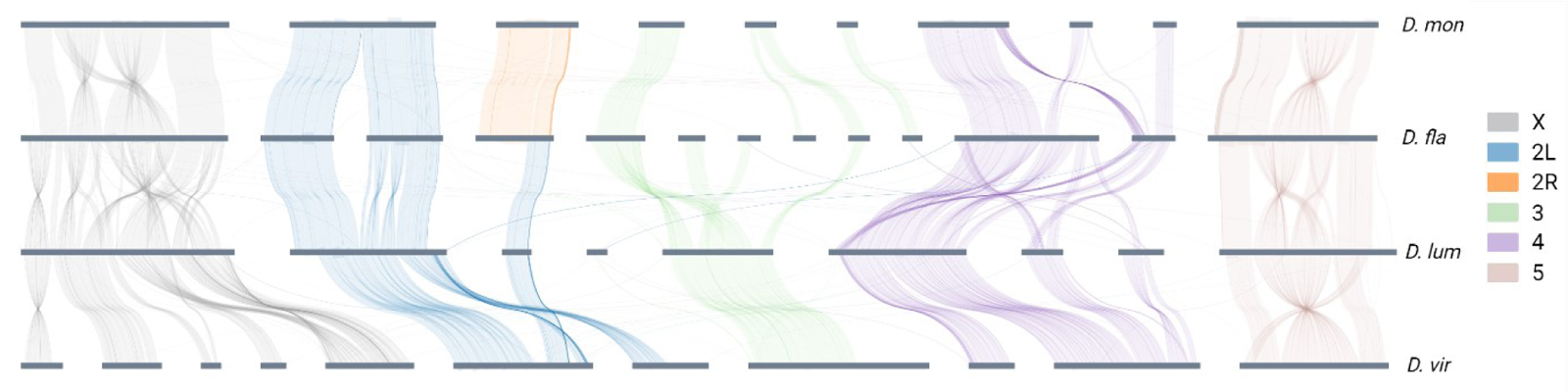
Chromosome synteny between *D. montana* (*D. mon*: monSE13F37), *D. flavomontana* (*D. fla*: flaMT13F11), *D. lummei* (*D. lum*) and *D. virilis* (*D. vir*). Different chromosomes and chromosome arms are marked with different colours. The plot shows contigs larger than 2Mb.

### Genetic differentiation and climatic variability of *D. montana* and *D. flavomontana* populations

To investigate the genetic structure of *D. montana* and *D. flavomontana* populations, we performed a principal component analysis (PCA) on the Illumina re-sequence data for the 12 and 9 wild-caught females of *D. montana* and *D. flavomontana*, respectively (Table S1 & S2). The PCA included 9,102,309 filtered SNPs from coding, intronic and intergenic regions. The first two principal components (PCs) had Eigenvalues >1, and PC1 explained majority (50%) of the total genetic variance and clearly separated *D. montana* and *D. flavomontana* samples from each other (Fig. 3A, Table S4). PC2 explained 4% of the total variance and captured variation mainly within *D. montana*, while variation in *D. flavomontana* was lower (Fig. 3A, Table S4). PC2 separated allopatric Alaskan *D. montana* populations (Honolulu Creek, Seward, Fairbanks) from sympatric mountainous and coastal *D. montana* populations, but also showed some variation within the allopatric and sympatric populations (Fig. 3A, Table S4).

Next, we explored climatic variability of fly sampling sites to determine the extent to which climatic conditions may have affected the genetic differentiation of the samples. We performed a PCA on 19 bioclimatic variables of each fly sampling site (Table S5, S6) to reduce correlations between the variables and summarized climatic patterns prevailing in the sites. This PCA revealed three principal components (PCs) with Eigenvalues >1, of which the first two PCs explained ∼80% of the climatic variation (Fig. 3B, Table S7). The first PC clearly separated inland and coastal populations and suggested that populations from the mountainous inland experience cold winters and high seasonal temperature variation, while coastal populations experience milder temperatures and high precipitation throughout the year (Fig. 3B, Table S8). The second PC separated populations based on summer temperatures and variation in diurnal temperatures and distinguished Alaskan populations (Honolulu Creek, Seward, Fairbanks) from the other populations (Fig. 3B, Table S8).

Together these results show that *D. montana* and *D. flavomontana* populations are genetically diverged regardless of their climatic origin and species coexistence. Genetic differentiation was greater among *D. montana* populations than among *D. flavomontana* populations, which is likely due to *D. montana*’s larger geographic range and the fact that *D. flavomontana* has spread across North America more recently than *D. montana* (Hoikkala & Poikela 2022). Finally, the genetic differentiation between allopatric (from Alaska) and sympatric (from the Rocky Mountains and the western coast) *D. montana* populations likely reflects a demographic history of intraspecific divergence, local adaptation to climatic conditions, or both.

**Figure 3.**
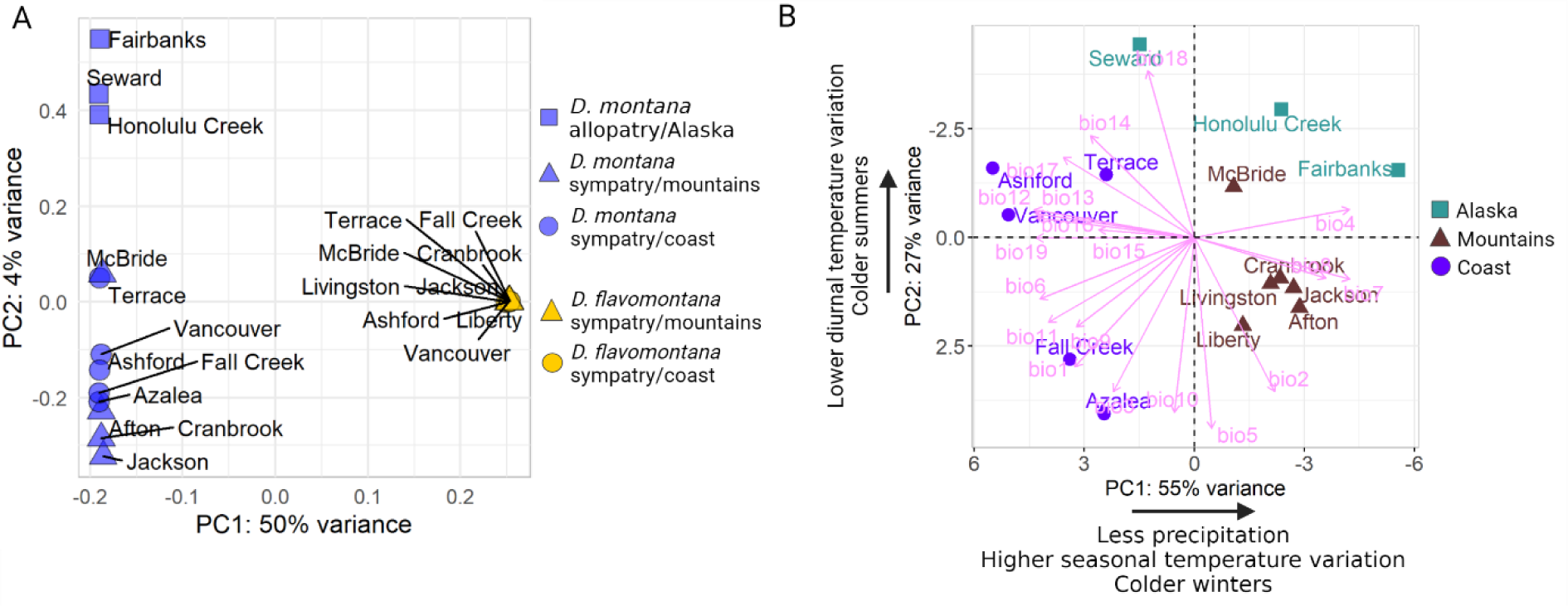
A principal component analysis (PCA) (A) on whole-genome SNP data of *D. montana* and *D. flavomontana* Illumina re-sequence samples originating from different sites of North America and (B) on 19 bioclimatic variables (detailed explanations in Table S5) of each fly sampling site.

### *D. montana* and *D. flavomontana* chromosomes differ by several large inversions

We combined long- and short-read genomic data to characterize inversion breakpoints in *D. montana* and *D. flavomontana*. We identified five large (>0.5Mb) inversions that were alternatively fixed between North American *D. montana* and *D. flavomontana* samples (Table S9, Fig. 2, S5-S10). The X chromosome contained three partly overlapping inversions, one of which was fixed in *D. montana* (7.2Mb) and two in *D. flavomontana* (11.0Mb and 3.1Mb) (Table S9, S5-S10). Chromosomes 4 and 5 each contained one inversion fixed in *D. flavomontana* (15.9Mb and 9.2Mb, respectively) (Table S9, Fig. S5). All these inversions were homozygous in Illumina re-sequenced individuals of both species (Table S9). In contrast, chromosomes 2 and 3 did not contain any fixed inversion differences. However, a subset of reads indicated that the left arm of the 2^nd^ chromosome (2L) contained an inversion (3.9Mb) that was heterozygous in all *D. montana* samples (Table S9, Fig. S5). Since this inversion signal is derived solely from raw reads and not from genome comparisons, we cannot exclude the possibility that this is a false positive. Because this putative inversion is not fixed between the species, it was excluded from further analysis. The sizes of inversions were obtained from the genome assemblies of each species (Table S9). Overall repeat density was higher at four of the ten breakpoints compared to the mean values for the X chromosome and autosomes (Fig. S11). Generally, high abundance of repeats may contribute to the origin of inversions (Kapun & Flatt 2019). Intriguingly, a known TE (Mariner-2_DVi) was found at the distal breakpoint of the shorter fixed X inversion in *D. flavomontana* but not in *D. montana* genomes, which could potentially be associated with the establishment of that inversion (Supplementary file 1). PacBio read support (ranging between 16 and 106 reads), as well as genes and repetitive regions located at inversion breakpoints are shown in Supplementary file 1.

Based on polytene chromosome studies (Stone et al. 1960), the three alternatively fixed inversions between *D. montana* and *D. flavomontana* on the X chromosome likely correspond to inversions E, F and G. These inversions were not distinguished in more detail in Stone et al. (1960), and, in contrast to our results, Stone et al. (1960) suggest that all three X inversions are fixed in *D. flavomontana*. The inversions on the 4^th^ and 5^th^ chromosome have been named J and E in karyotype studies, respectively (Stone et al. 1960).

The average size of the inversions fixed in *D. montana* was 7.2Mb and in *D. flavomontana* 9.8Mb (Table S9), which resembles the average reported size of inversions in animals and plants (8.4Mb) (Wellenreuther & Bernatchez 2018). Our finding of a larger number of inversions on the X is consistent with theory showing that the fixation probability of X chromosomal inversions is higher than that of autosomal inversions, because selection can more effectively favour beneficial and purify deleterious recessive X-linked alleles than autosomal ones (Charlesworth et al. 2018, 1987; Connallon et al. 2018; Vicoso & Charlesworth 2006). Moreover, the higher content of repetitive sequences we find on the X chromosome compared to autosomes (Fig. S11), which has also been observed in other *Drosophila* species (Cridland et al. 2013), may predispose the X chromosome to sequence breakage and thus facilitate the formation of inversions (Kapun & Flatt 2019).

The polytene chromosome studies by Stone et al. (1960) and Throckmorton (1982) suggest that *D. montana* and *D. flavomontana* carry additional inversions that were not detected in this study. In particular, *D. flavomontana* may harbour one fixed inversion of unknown size on chromosome 3 (inversion E; Stone et al. 1960), which we might have missed due to the higher fragmentation of this chromosome compared to the other chromosome contigs (Table S10). Given our limited sample size and explicit focus on fixed inversion differences between species, polymorphic inversions previously found in these species (Stone et al. 1960), which may be associated with local adaptation or other evolutionary processes (Fang et al. 2012; Kapun et al. 2016; Wallberg et al. 2017), were also not identified here.

### Genetic divergence between *D. montana* and *D. flavomontana* is greater inside than outside inversions

We analysed the mean genetic divergence (d_xy_) to test whether inversions have reduced recombination and introgression between *D. montana* and *D. flavomontana*, and if so, whether this is ancient or recent. In the latter case, d_XY_ should be lower in sympatry compared to allopatry (Harrison & Larson 2014; Noor & Bennett 2009). We estimated d_xy_ separately for coding, intronic and intergenic sequences and inverted and colinear regions of the genome. Given the potentially different evolutionary history of the X chromosome and the autosomes (Charlesworth et al. 2018; Vicoso & Charlesworth 2006), analyses for the X were performed separately. We also carried out separate analyses for allopatric and sympatric comparisons of the species. We focus mainly on absolute measure of genetic divergence (d_xy_), since relative differentiation, i.e. F_ST_, measures both variation in genetic diversity and divergence, and so is harder to interpret (Cruickshank & Hahn 2014; Charlesworth 1998; Noor & Bennett 2009).

Mean divergence (d_xy_) between *D. montana* and *D. flavomontana* was remarkably similar for intergenic and intronic sequences but much lower for coding sequences (Fig. 4, Table S11), as expected given the stronger selective constraint on coding sites (Halligan & Keightley 2006). Moreover, d_xy_ was slightly, but consistently lower for sympatric compared to allopatric comparisons of the species across all chromosome regions (Fig. 4, Table S11).

At non-coding sequences (i.e. intergenic and intronic), mean d_xy_ was consistently higher in inverted compared to colinear regions in allopatric and sympatric comparisons (Fig. 4, Table S11). At coding sequences, mean d_xy_ was increased for inversions on the 4^th^ and the X chromosome compared to colinear regions both in allopatric and sympatric comparisons (Fig. 4, Table S11). Plotting d_xy_ in sliding windows showed an increase in genetic divergence, especially around the inversion breakpoints and for overlapping X inversions for sympatric and allopatric comparisons of the species (Fig. 5; chromosomes shown individually in Fig. S12). A similar increase in d_xy_ within *D. montana* (intraspecific comparison) was seen around some of the breakpoints on chromosome 4 and 5, but not on chromosome X (Fig. 5). Based on a correlation analysis between inter and intraspecific d_xy_, the chromosome 4 inversion appears to be an outlier in having a greater correlation than colinear regions (Fig. S13). This increased d_xy_ both in inter and intraspecific crosses is potentially explained by a number of inversions that are polymorphic within *D. montana* on chromosome 4 (Stone et al. 1960; Throckmorton 1982).

F_ST_ was also generally higher for inverted compared to colinear regions, especially in allopatry, although these differences were non-significant (Table S11). The fact that the differences between inverted and colinear regions are less clear for F_ST_ than d_xy_ reflects the susceptibility of F_ST_ to variation in genetic diversity (Fig. S14-S16).

Overall, our finding of higher genetic divergence inside compared to outside of inversions is consistent with the idea that inversions suppress gene exchange and facilitate the accumulation/preservation of genetic differences (Fig. 4, Table S11; Navarro and Barton 2003; Kirkpatrick and Barton 2006). We also found that genetic divergence was highest around inversion breakpoints and in the series of overlapping inversions on the X (Fig. 5), where recombination is the most suppressed (Hoffmann and Rieseberg 2008). Similar signatures of elevated genetic divergence between closely related species inside and around inversion breakpoints have been detected e.g. in other *Drosophila* species pairs (Noor et al. 2007; Kulathinal et al. 2009; Lohse et al. 2015), *Helianthus* sunflowers (Barb et al. 2014), *Sorex* shrews (Basset et al. 2006), and *Anopheles* mosquito (Michel et al. 2006). Finally, our finding of lower genetic divergence in sympatry compared to allopatry (Fig. 4, Table S11) is consistent with low levels of recent introgression in sympatry (Harrison & Larson 2014; Noor & Bennett 2009).

**Figure 4.**
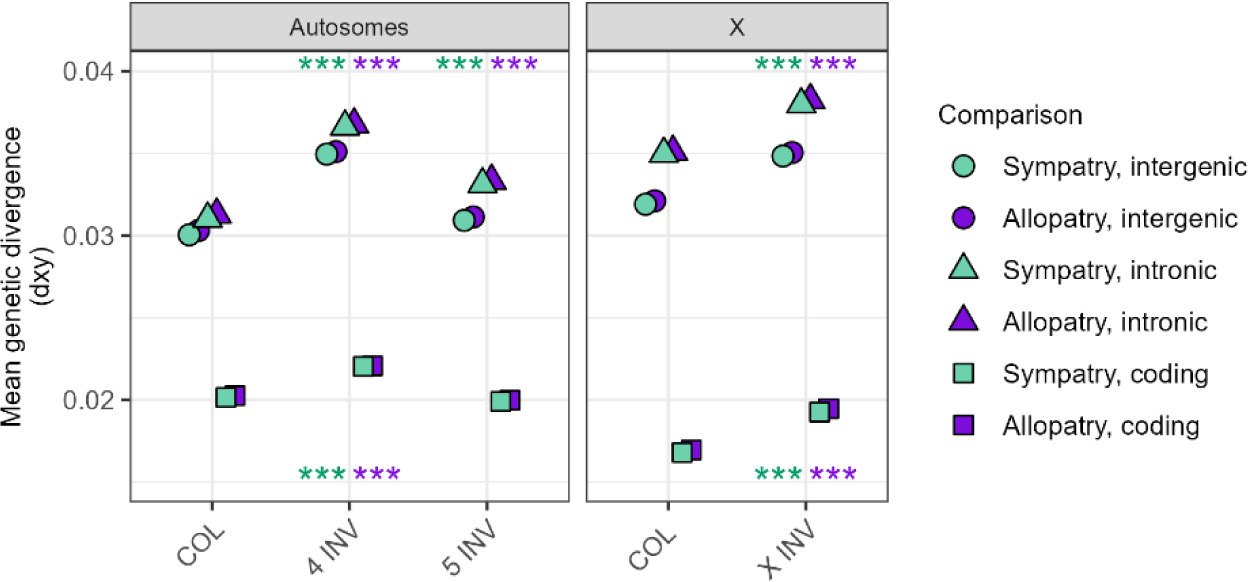
Mean genetic divergence (measured as dxy) at intergenic, intronic and coding sequences of colinear (COL, background) and inverted (INV) chromosome partitions on the autosomes and the X. Divergence is shown for allopatric (purple) and sympatric (green) comparisons of *D. montana* and *D. flavomontana*. Significance levels were inferred from simulations, where COL regions were compared to INV regions separately for autosomes and the X chromosome, for intergenic, intronic and coding sequences, and for allopatric and sympatric comparisons (*** P < 0.001; P-values for intergenic and intronic sequences shown above and for coding sequences below dots; Table S11).

**Figure 5.**
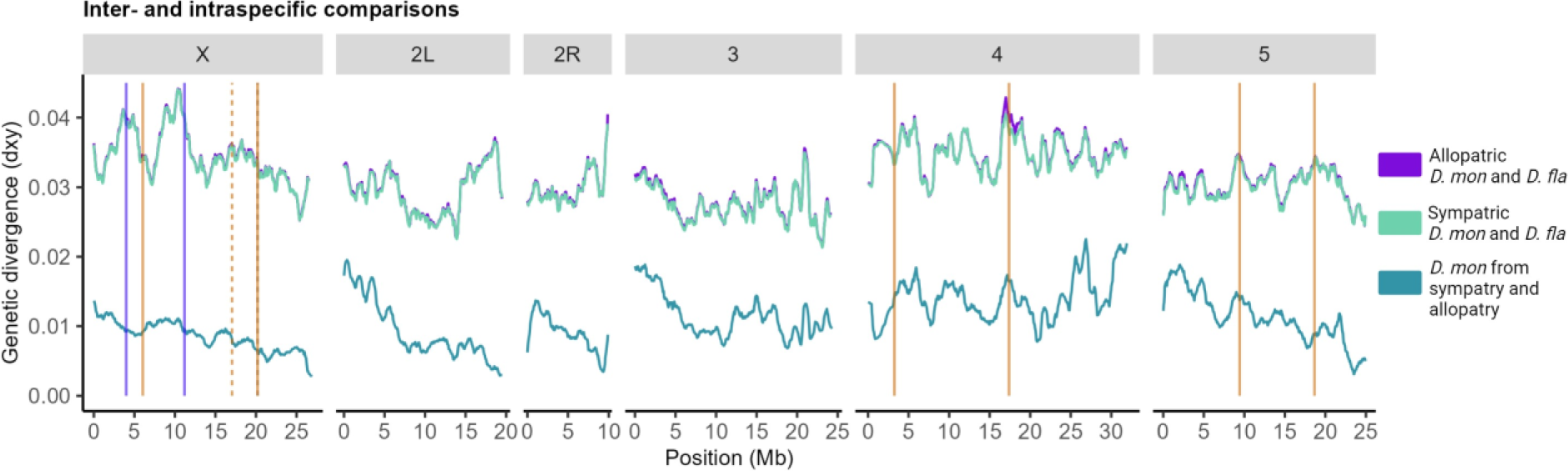
Genetic divergence (measured as dxy) across the genome (including intergenic regions) in sliding windows (window size 5,000 blocks, step size 500 blocks, block length 64b) for allopatric and sympatric comparisons of *D. montana* and *D. flavomontana* (interspecific), and for *D. montana* originating from allopatry and sympatry (intraspecific). Vertical lines represent breakpoints of different inversions (Fig. S5, Table S9): blue solid lines, and orange solid and dashed lines indicate alternatively fixed inversions of *D. montana* and *D. flavomontana*, respectively.

### No evidence for genes being under divergent selection in inversions

Alternatively fixed inversions may become hotspots for positively selected genetic differences, which can enhance adaptation and/or give rise to pre- and postzygotic barriers (Navarro & Barton 2003; Kirkpatrick & Barton 2006). To investigate whether genes under divergent selection are enriched within inversions, we performed a dN/dS analysis for *D. montana* and *D. flavomontana* using branch-site model in codeML (Supplementary file 2).

We found 157 genes with evidence for divergent selection in *D. montana* and *D. flavomontana* (out of a total of 7423 single copy orthologues (SCOs)). Altogether 45 positively selected genes were located inside inversions (1997 SCOs within inversions altogether), but the inversions were not significantly enriched for genes under divergent selection (G=0.159, P=0.690). However, it is unlikely that we detected all genes under divergent selection since statistical power of the approach may be relatively low for closely related species. While we find no signal of increased divergent selection in inversions in terms of numbers of genes involved, the divergent genes inside inversions we identified include plausible targets for selection on potential barrier traits, such as chemoreception (odorant receptor 19a) (Hallem & Carlson 2006) and male fertility (testis-specific serine/threonine-protein kinase 3) (Nozawa et al. 2023). Moreover, even though none of the genes located nearby the inversion breakpoints were under divergent selection (Supplementary file 1 & 2), some of them may still be targets of selection as they have translocated alongside the inversions, and such translocations may give rise to new expression patterns (Villoutreix et al. 2021).

### Hierarchical model comparison suggests species diverge with very low levels of post-divergence gene flow from *D. montana* to *D. flavomontana*

We used gIMble (Laetsch et al. 2023), an analytic likelihood method, to fit a series of demographic models of species divergence with and without long-term post-divergence gene flow, i.e. isolation with migration (IM) and strict divergence (DIV) models (Fig. S17), to the data summarised in terms of the bSFS (see Methods). The evolutionary history of the X chromosome (Charlesworth et al. 2018; Vicoso & Charlesworth 2006) and inversions (Lohse et al. 2015; Fuller et al. 2018) may differ from other chromosome regions, and these genomic partitions were therefore analysed separately from colinear, autosomal regions. To minimize the direct effects of selection, our initial analysis was limited to intergenic sequences of the colinear autosomal regions (repetitive regions were excluded). We performed separate analyses for sympatric and allopatric comparisons of *D. montana* and *D. flavomontana*. To evaluate the timing of potential recent introgression in sympatry compared to allopatry, we also performed a separate analysis for intraspecific comparison of *D. montana* (*D. montana* living in contact vs in isolation with *D. flavomontana*). We carried out this initial model comparison for the DIV and two IM models four times, using both *D. montana* and *D. flavomontana* as a reference genome to evaluate the potential effects of reference bias, and performed separate analyses for two different block lengths (64b and 128b). Parameter estimates and support values (lnCL) under all demographic models are shown in Table 1 for 64b blocks and using *D. montana* as a reference. Analogous analyses for all four combinations of block length and reference genomes are given in Table S12.

For both allopatric and sympatric comparisons and for three of the four combinations of block lengths and reference genomes used, the best-fitting demographic scenario was an IM model assuming introgression from *D. montana* into *D. flavomontana* (Table 1 & S12). Our parametric bootstrap analyses showed that the improvement in fit of this IM model compared to the DIV model was significant suggesting a low but genuine signal of introgression (Fig. S18 & S19). The only exception was the analysis using shorter 64b blocks and *D. flavomontana* as a reference genome. In this case, the DIV model could not be rejected (Fig. S19). However, estimates for all parameters (T and N_e_s) were extremely similar regardless of the model (DIV and IM), block size (64b and 128b) and reference genome (*D. montana* and *D. flavomontana*) used (Table S12). Given the overall support for post-divergence gene flow and inherent bias of multilocus inference to underestimate migration, we assume an IM model with migration from *D. montana* into *D. flavomontana* as the best-fitting / most plausible scenario throughout for all subsequent analyses (Table 1). Yusuf et al. (2022) also recently found signatures of introgression between *D. montana* and *D. flavomontana* using a different approach, which gives further support for our introgression signal.

In contrast, for the intraspecific comparison of *D. montana*, the DIV model could not be rejected in any analysis. When using 64b blocks, DIV and IM models had equal support, irrespective of which species was used as a reference (Table 1 & S12). Analyses based on longer 128b blocks estimated slightly higher support for an IM model assuming post-divergence gene flow from allopatric (Alaskan) *D. montana* to sympatric (coastal/mountain) *D. montana* (Table 1 & S12). However, the parametric bootstrap analyses showed that the improvement in fit compared to the simpler DIV model was non-significant (Fig. S18 & S19). Consequently, the subsequent intraspecific analyses were conducted using the DIV model (Table 1).

**Table 1.** Support (measured as ΔlnCL) and parameter estimates for divergence time (*T* in years/generations), migration rate (*m*) and effective population sizes (Ne) for studied populations and their common ancestral population under strict divergence (DIV, *m*=0) and isolation with migration (IM) models with both gene flow directions. The model comparison is based on 64b blocks and the *D. montana* reference genome, and was performed for intergenic autosomal colinear regions to minimize the effects of selection. Grey shading indicates the best-fit model for each comparison.

| Comparison | Model | *D. mon* $N_e$ | *D. fla* $N_e$ | Ancestral $N_e$ | $T$ | $m$ | lnCL | $\Delta\ln\text{CL}$ |
| --- | --- | --- | --- | --- | --- | --- | --- | --- |
| Allopatric | DIV | 693,000 | 395,000 | 1,464,000 | 2,379,000 | - | -45651205 | 12869 |
|  | IM *D. mon* --> *D. fla* | 705,000 | 382,000 | 1,403,000 | 2,539,000 | 1.09E-08 | -45638336 | 0 |
|  | IM *D. fla* --> *D. mon* | 692,000 | 396,000 | 1,457,000 | 2,398,000 | 1.21E-09 | -45650952 | 12616 |
| Sympatric | DIV | 720,000 | 392,000 | 1,459,000 | 2,343,000 | - | -136875659 | 47790 |
|  | IM *D. mon* --> *D. fla* | 735,000 | 377,000 | 1,388,000 | 2,526,000 | 1.29E-08 | -136827869 | 0 |
|  | IM *D. fla* --> *D. mon* | 719,000 | 393,000 | 1,447,000 | 2,376,000 | 2.13E-09 | -136873722 | 45853 |
| | Model | *D. mon allop* $N_e$ | *D. mon symp* $N_e$ | Ancestral $N_e$ | $T$ | $m$ | lnCL | $\Delta\ln\text{CL}$ |
| Intraspecific | DIV | 1,087,000 | 1,560,000 | 858,000 | 210,000 | - | -39887382 | 0 |
|  | IM *D. mon symp.* --> *D. mon allop.* | 1,087,000 | 1,560,000 | 858,000 | 210,000 | 1.50E-15 | -39887382 | 0 |
|  | IM *D. mon allop.* --> *D. mon symp.* | 1,079,000 | 1,441,000 | 858,000 | 217,000 | 3.32E-07 | -39887408 | 27 |

### Species-specific inversions were fixed earlier or around the species’ split and introgression was lower inside compared to outside of inversions and in allopatry compared to sympatry

We used the best-fit isolation with migration (IM) model (Table 1) to examine the potential role of inversions and introgression in the speciation history of *D. montana* and *D. flavomontana*. As before, all analyses were limited to intergenic regions to minimize the effects of selection, and separate analyses were carried out for the X chromosome and autosomes, for inverted and colinear regions, and for sympatric and allopatric populations of the species. To estimate the timing of potential recent introgression, we analysed the split time for *D. montana* living in contact (sympatry) or in isolation (allopatry) with *D. flavomontana* using the simpler strict divergence (DIV) model (intraspecific comparison; Table 1).

Taking the estimates for the colinear autosomal background as face value, *D. montana* and *D. flavomontana* have diverged ca 2.5 Mya (Table 2, Fig. 6A & 6C). The divergence time estimates of the inversions differ from each other, and the inversions on the 4^th^, 5^th^ and the X chromosome predate the divergence time estimated for the colinear background (ca 2.8-3.3 Mya) (Table 2, Fig. 6A & 6C). For all chromosome partitions, genetic diversity (π) and the effective population size (N_e_) of *D. montana* were approximately two times as large as those of *D. flavomontana* (Table 2, Fig. S14). *D. montana* populations living in contact (sympatry) and in isolation (allopatry) with *D. flavomontana* have diverged approximately 210 000 years ago (Table 2, Fig. 6C), an order of magnitude more recent than the split between *D. montana* and *D. flavomontana*.

Estimated long-term gene flow from *D. montana* to *D. flavomontana* was lower inside than outside of inversions both on the autosomes and the X (Table 2, Fig. 6B), which is in accordance with the finding that genetic divergence of non-coding (intergenic and intronic) sequences was consistently higher inside than outside of inversions (Fig. 4, Table S11). Moreover, migration rate estimates were higher in sympatry compared to allopatry (Table 2, Fig. 6B), which again agrees with the slightly, but consistently lower genetic divergence in sympatric compared to allopatric comparisons of the species (Fig. 4, Table S11).

**Table 2.** Parameters for effective populations sizes, divergence time (*T* in years/generations), and migration rate (M) estimated from 64b blocks under the IM*mon* è *fla* model for allopatric and sympatric comparisons and under the DIV model for intraspecific comparison.

| Comparison | Genomic region | | Ne ancestral | Ne *D.mon* | Ne *D.flu* | $T$ | $M$ |
| --- | --- | --- | --- | --- | --- | --- | --- |
| Allopatry | Autosomes | COL | 1,403,000 | 705,000 | 382,000 | 2,539,000 | 0.0083 |
|  |  | 4 INV | 1,644,000 | 798,000 | 474,000 | 2,941,000 | 0.0049 |
|  |  | 5 INV | 1,364,000 | 718,000 | 396,000 | 2,777,000 | 0.0060 |
|  | X | COL | 1,700,000 | 443,000 | 225,000 | 2,605,000 | 0.0074 |
|  |  | X INV | 1,904,000 | 441,000 | 368,000 | 2,829,000 | 0.0038 |
| Sympatry | Autosomes | COL | 1,388,000 | 735,000 | 377,000 | 2,526,000 | 0.0097 |
|  |  | 4 INV | 1,603,000 | 865,000 | 470,000 | 2,988,000 | 0.0067 |
|  |  | 5 INV | 1,321,000 | 752,000 | 392,000 | 2,823,000 | 0.0076 |
|  | X | COL | 1,592,000 | 493,000 | 219,000 | 2,769,000 | 0.0088 |
|  |  | X INV | 1,591,000 | 607,000 | 365,000 | 3,321,000 | 0.0078 |
| Comparison | Genomic region | | Ne ancestral | Ne allopat *D. mon* | Ne symp *D. mon* | $T$ | |
| Intraspecific | Autosomes | COL | 858,000 | 1,087,000 | 1,560,000 | 210,000 |  |

**Figure 6.**
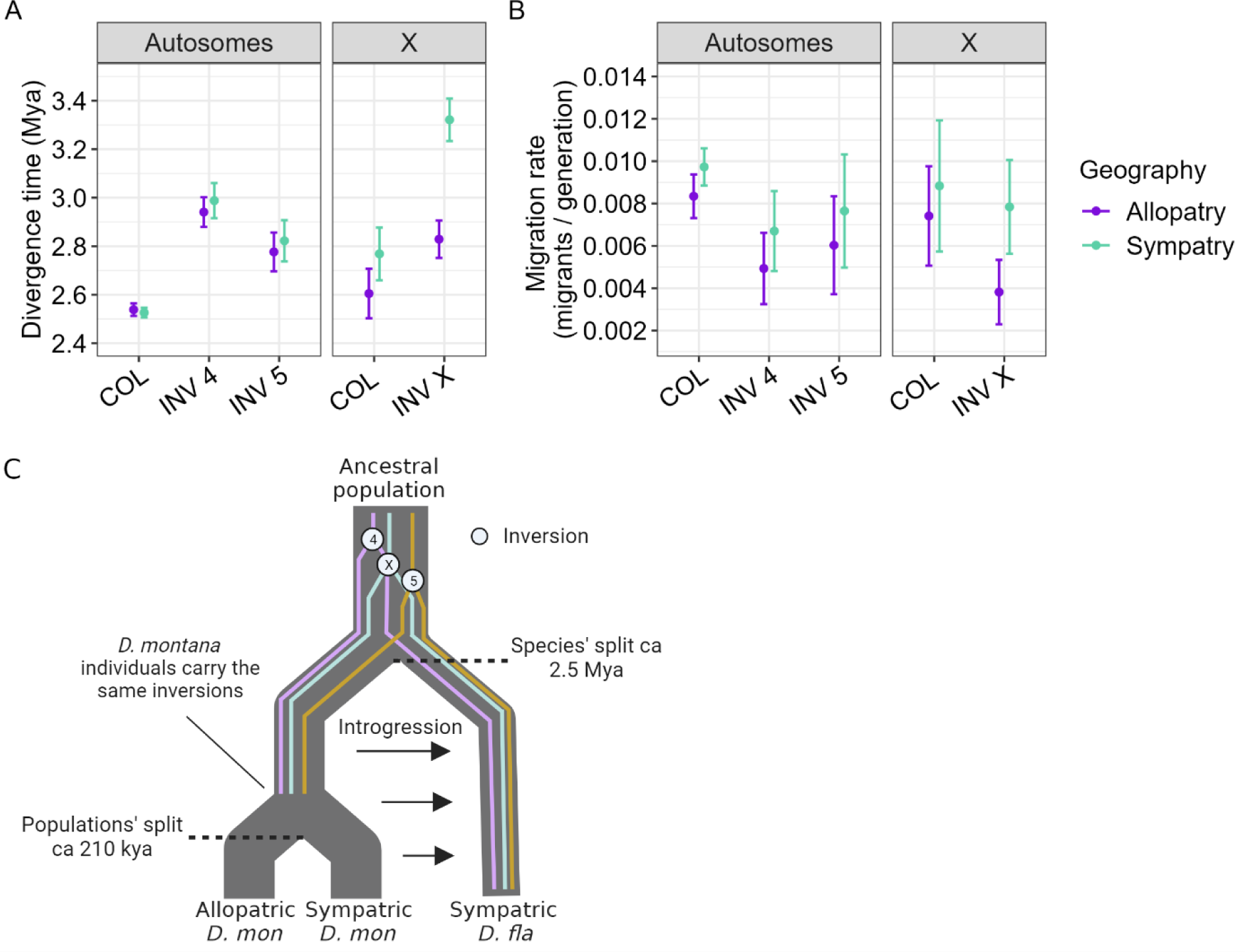
Estimates of (A) split times and (B) migration rates between *D. montana* and *D. flavomontana* for different chromosome partitions and for allopatric (purple) and sympatric (green) comparisons. Confidence intervals were estimated as +/-2 SD. (C) Illustration of the likely evolutionary history of *D. montana* and *D. flavomontana*.

Taken together, our analyses suggest that *D. montana* and *D. flavomontana* diverged ca 2.5 Mya from a large ancestral population, which is broadly compatible with a recent estimate of 3.8 Mya based on small introns and the same molecular clock calibration (Yusuf et al. 2022). Crucially, split time estimates for all five fixed inversions we have identified on chromosomes X, 4 and 5 predate the estimated species split time based on the colinear background, which implies that these inversions must have existed already in the common ancestral population. In other words, we can rule out the possibility that the inversions arose after the onset of species divergence (in which case we would expect the divergence time estimates of inversions to overlap the estimated species divergence time). The reduced introgression for inversions compared to colinear regions we have estimated is a clear and expected consequence of reduced recombination and gene flow between alternative arrangements at each inversion.

What is less clear, is the extent to which local adaptation in the face of gene flow in the ancestral population facilitated the fixation of these inversions (and vice versa) or whether the fixed inversions are a mere byproduct of population processes unrelated to speciation. The fact that three fixed inversions on the X are i) overlapping and ii) associated with a strong incompatibility preventing introgression from *D. montana* to *D. flavomontana* across the entire X chromosome (Poikela et al. 2022) suggest that at least the inversions on the X contributed to the build-up of reproductive isolation and acted as barriers to gene flow early on in the speciation process (Noor & Bennett 2009; Fuller et al. 2018). For example, these inversions may have been important in initial ecological divergence of local populations of the ancestor, followed by the fast accumulation of genetic divergence and genetic incompatibilities. In contrast, we currently have no evidence that the inversions on chromosomes 4 and 5 are enriched for BDMIs (Poikela et al. 2022) or loci under divergent selection, so we cannot rule out a scenario in which these inversions have been maintained in the ancestral populations by balancing selection and have subsequently become fixed between *D. montana* and *D. flavomontana* simply by chance (Guerrero & Hahn 2017). In fact, even for the X inversions we cannot verify whether the associated incompatibility allele(s) arose before or around the species’ split, or afterwards (post-speciation event). We stress that our comparison of divergence times estimated under the IM model between inverted and colinear parts of the genome relies on the assumption of neutrality (which is why we have restricted analyses of demographic history to intergenic sequence). If however, some fraction of the intergenic partition is under selective constraint, we might expect higher genetic divergence within inversion: Berdan et al. (2021) recently showed using simulations that heterozygous inversions may accumulate non-adaptive, mildly deleterious mutations via less effective purifying selection within inversions, leading to higher genetic divergence even without any reduction in recombination between alternative arrangements.

Although we find evidence for post-divergence gene flow, it is worth highlighting that our estimate of long-term rate of migration from *D. montana* to *D. flavomontana* is extremely small compared to analogous estimates for other young *Drosophila* sister species (Lohse et al. 2015); e.g. *D. mojavensis* and *D. arizonae* have approximately one migrant per generation, while our estimate for *D. montana* and *D. flavomontana* is roughly 1 migrant in 80 generations, two orders of magnitude lower. Thus, even the total probability of a lineage sampled in *D. flavomontana* to trace back to *D. montana* via migration (1-e∧(-T M)) is only 3.2%. This low rate of long-term effective migration agrees well with our previous evidence for strong pre- and postzygotic barriers between the species (Poikela et al. 2019). In addition, the species’ differences in the usage of host trees (Throckmorton 1982) and the ability to tolerate cold (Poikela et al. 2021) might have contributed to ecological isolation and reduced their encounters in nature. Intriguingly, we found higher levels of introgression in sympatry compared to allopatry, which suggests at least some introgression from *D. montana* to *D. flavomontana* over the past ∼210 000 years, i.e. after the allopatric (Alaskan) *D. montana* populations diverged from *D. montana* coexisting with *D. flavomontana*. Even low levels of introgression and selection against introgressed ancestry in the new genetic background may facilitate reinforcement of prezygotic barriers to prevent maladaptive hybridization between species, and eventually complete the speciation process (Cruickshank & Hahn 2014; Servedio & Noor 2003). This is consistent with our previous finding that *D. flavomontana* has developed stronger prezygotic barriers against *D. montana* in sympatry compared to allopatry, presumably as a result of reinforcement (Poikela et al. 2019).

Our demographic inferences are limited in several ways. Firstly, the IM model is overly simplistic in assuming an instantaneous onset of species divergence and a constant rate of introgression throughout the species’ evolutionary history. However, given the overall extremely low estimate of gene flow and the computational limitations of gIMble, we have not attempted to fit – and therefore cannot exclude − more realistic (but necessarily more complex) demographic scenarios of either historical gene flow which reduced due to the emergence of strong barriers or sudden discrete bursts of admixture following periods of complete isolation. Secondly, our inference ignores recombination within blocks, a simplifying assumption that is known to lead to biased parameter estimates (Wall 2003). In particular, we found that the estimates of *T* obtained from parametric bootstrap replicates (simulated with recombination) are substantially larger (∼3.4 MY) than the true values (Table 2, Fig. 6 & S20), which suggests that we have overestimated species divergence time overall. Finally, our approach of fitting an IM model to inverted regions ignores the fact that inversions arise in a single individual and may be fixed in a selective sweep. An inversion arising and sweeping to fixation immediately after the onset of species divergence would result in a lower estimate of N_e_ for the species in which they fixed. If anything, we see the opposite pattern: i.e. larger estimates of *D. flavomontana* N_e_ for the inversions on chromosomes 4 and 5 compared to the colinear background (Table 1), which is again compatible with an inversion origin in the ancestral population before the estimated species split. It is striking that all inversions date to a short interval just before the species split (∼600k years/generations) which is the same order as the (ancestral) population size. Given that we infer a substantially larger effective size for the ancestral population than fo*r D. montana* and *D. flavomontana*, one could interpret the interval in the ancestral population in which the inversions arose as the period of (rather than before) speciation.

Even though many species pairs differ from each other by multiple inversions, the majority of inversion differences must have arisen after speciation (Faria & Navarro 2010). Performing pairwise comparisons for younger and older species would offer a more holistic view of the role of inversions in speciation events. In our case, characterizing inversions and investigating divergence times and introgression across all species of the *montana* phylad of the *virilis* group (*D. montana*, *D. flavomontana*, *D. borealis* and *D. lacicola*) (Hoikkala & Poikela 2022) could provide valuable additional information. In general, investigating millions of years old events by fitting necessarily drastically simplified scenarios of reality involve uncertainties.

## Conclusions

It has proven extremely difficult to test if and how inversions facilitate speciation, and empirical evidence on the role of inversions in speciation is largely lacking (Faria & Navarro 2010; Fuller et al. 2019). We explored these questions in two sister species of the *D. virilis* group, *D. montana* and *D. flavomontana*. Our main goals were (1) to characterize alternatively fixed chromosomal inversions of *D. montana* and *D. flavomontana*, (2) to investigate the age of the inversions, and (3) to identify whether the inversions have restricted gene exchange between *D. montana* and *D. flavomontana* during or before the onset of species divergence, which could have facilitated the accumulation or preservation of incompatibilities in the presence of gene flow.

Taking advantage of long- and short-read genome sequencing technologies, we generated the high-quality contiguous reference assemblies for *D. montana* and *D. flavomontana*. These genomes enabled us to accurately characterize inversions that are alternatively fixed between these sister species across their distribution area in North America. We were able to assign the majority of these to inversions that were previously described for the species based on polytene chromosome studies (Stone et al. 1960). Our analyses show that the inversions on chromosomes X, 4 and 5 arose before the onset of species divergence. Thus, the elevated genetic divergence within inversions results most likely from restricted recombination between alternative rearrangements, which were either under balancing selection or locally beneficial in different populations of the ancestral form. However, the X inversions have been found to contain strong BDMI(s) that effectively restricts introgression from *D. montana* to *D. flavomontana* across the X chromosome in the first few backcross generations (Poikela et al. 2022), and provide evidence for the enrichment of BDMIs within inversions. Accordingly, our results are compatible with the idea that ancestrally polymorphic inversions, particularly the X chromosomal inversions in our case, can drive speciation potentially by facilitating initial ecological divergence and fast accumulation of genetic divergence and genetic incompatibilities (Fuller et al. 2018), while colinear regions keep exchanging genetic material until strong reproductive isolation has formed.

Even though the estimates of introgression between the species were extremely low, *D. flavomontana* has experienced some introgression from *D. montana* over the past ∼210 000 years in sympatric populations of the species. In general, selection can strengthen prezygotic barriers between species in response to low levels of poor functioning introgressed alleles, which likely leads to the strengthening of overall reproductive isolation and the completion of the speciation process (Cruickshank & Hahn 2014; Servedio & Noor 2003). This agrees with our previous evidence on *D. flavomontana* having developed stronger prezygotic barriers against *D. montana* in sympatric compared to allopatric populations of the species, potentially as a result of reinforcement (Poikela et al. 2019).

Overall, our results are compatible with the idea that inversions may be early triggers of speciation process and highlight the value of interpreting the evolutionary effects of inversions through the lens of demographic models. However, in doing so, we have ignored much of the mechanistic and selective details of inversion evolution. Regions with repetitive sequences, such as transposable elements, tRNAs, ribosomal genes or segmental duplications, are prone to breakage and are often the initial source of an inversion (Kapun & Flatt 2019). An in-depth investigation into the repetitive sequences or small structural variations around the inversion breakpoints would increase our understanding on how the inversions originated in the first place. Moreover, inversions are not static through their lifetime, but evolve in response to changes in selection, genetic drift, new mutations and gene flux (occurring via double cross-overs and gene conversion), as well as by interactions with other parts of the genome (Faria et al. 2018). Given the many, sometimes entangled processes affecting the origin and the fixation of inversions, models that can extract information about both demography and the selective forces acting on inversions in the early stages of speciation are the next obvious step in understanding how inversions facilitate the origin of species (Faria et al. 2018).

## Materials and Methods

### Sample collections and maintenance

*D. montana* and *D. flavomontana* females were collected from several sites in the Rocky Mountains and along the western coast of North America, and Alaska 2013−2015 (Fig. 1, Table S1). Sites in the Rocky Mountains and the western coast of North America are either inhabited by both species (sympatric sites: Jackson, Cranbrook, McBride, Terrace, Vancouver, Ashford, Fall Creek), or by one of the species with nearby sites inhabited by both species (parapatric sites: Liberty, Afton, Livingston, Azalea) (Fig. 1, Table S1). *D. montana* also has stable populations in high latitudes in Alaska, where *D. flavomontana* does not exist. We refer to the comparisons between *D. montana* and *D. flavomontana* from the Rocky Mountains and from the western coast as “sympatry”, and those between *D. montana* from Alaska and *D. flavomontana* from the Rocky Mountains or the western coast as “allopatry” (Fig. 1). Intraspecific comparison was performed for *D. montana* living in isolation (allopatry) and in contact (sympatry) with *D. flavomontana* (Fig. 1).

The newly collected females were brought to the fly laboratory, with a constant light, 19±1 °C and ∼60% humidity, at the University of Jyväskylä, Finland. Females that had mated with one or several males in nature were allowed to lay eggs in malt vials for several days, after which they were stored in 70% EtOH at -20 °C. The emerged F_1_ progeny of each female were kept together to produce the next generation and to establish isofemale strains. After that also the F_1_ females were stored in 70% EtOH at -20 °C.

### DNA extractions and sequencing

We performed PacBio long-read sequencing from two *D. montana* and two *D. flavomontana* isofemale strains that had been kept in the fly laboratory since their establishment (Fig. 1, Table S1). DNA of the Seward *D. montana* and both *D. flavomontana* samples were extracted from a pool of 60 three days old females per isofemale strain using cetyltrimethylammonium bromide (CTAB) solution with RNAse treatment, Phenol-Chloroform-Isoamyl alcohol (25:24:1) and Chloroform-Isoamyl alcohol (24:1) washing steps, and ethanol precipitation at the University of Jyväskylä, Finland. DNA of the Jackson *D. montana* sample was extracted with the “DNA Extraction SOP For Animal Tissue” protocol and purified with Ampure beads at BGI (Beijing Genomics Institute). Quality-checked DNA extractions of the Seward *D. montana* sample and both *D. flavomontana* samples were used to generate >15kb PacBio libraries, which were all sequenced on two SMRT cells within a PacBio Sequel system (Pacific Biosciences, United States) at the Norwegian Sequencing Centre in 2018. DNA of the Jackson *D. montana* sample was used to generate >20kb PacBio libraries and was sequenced on one SMRT cell using the PacBio Sequel system at BGI in 2019. Average PacBio raw read coverage was 27-35X per sample, except for the Jackson *D. montana* sample which was sequenced at 77X coverage. Detailed information on the PacBio raw reads of each sample is provided in Table S2.

We generated Illumina re-sequencing data for 12 *D. montana* and 9 *D. flavomontana* single wild-caught females or their F_1_ daughters from several locations in North America (Fig. 1, Table S1). DNA extractions were carried out at the University of Jyväskylä, Finland, using a CTAB method as described above. Quality-checked DNA extractions were used to produce an Illumina library for each sample in three batches. First, Nextera libraries were used to generate 150b paired-end (PE) reads on two lanes using HiSeq4000 Illumina instrument at Edinburgh Genomics in 2017. Second, one Truseq library was used to generate 150b PE reads on one lane of a HiSeq4000 Illumina instrument at the Norwegian Sequencing Centre in 2018. Third, Truseq libraries were used to generate 150b PE reads on one lane of a HiSeq X-Ten Illumina instrument at BGI in 2019. We generated on average 53-94X coverage per sample, except for the *D. montana* sample from Seward, which was sequenced to 435X coverage. Detailed information on Illumina raw reads is provided in Table S2.

### *De novo* genome assemblies, scaffolding and chromosome synteny

We generated initial *de novo* assemblies for each PacBio dataset and the respective Illumina reads using the wtdbg2 pipeline v2.5 (RedBean; Ruan & Li, 2020) and MaSuRCA hybrid assembler v3.3.9 (Zimin et al., 2017). To improve assembly contiguity, we used quickmerge for both assemblies of each sample (Chakraborty et al. 2016). The initial assembly statistics are given in Table S13. We polished the resulting assemblies with the respective Illumina reads using Pilon v1.23 (Walker et al. 2014) and removed uncollapsed heterozygous regions using purge_dups (Guan et al. 2020).

We identified genomic contaminants in the assemblies with BlobTools v1.1 (Laetsch & Blaxter 2017). PacBio and Illumina reads were first mapped back to each assembly with minimap2 (Li, 2018) and BWA mem (Burrows-Wheeler Aligner) v0.7.17 (Li & Durbin, 2009), respectively. Contigs in the assemblies were then partitioned into taxonomic groups based on similarity search against the NCBI nucleotide database (BLASTn 2.9.0+; Camacho et al., 2009) and Uniref90 (Diamond v0.9.17; Buchfink, Xie, & Huson, 2015). Finally, contigs were visualized on a scatter plot and coloured by putative taxonomic groups (Fig. S1-S4). Non-Diptera contigs were removed manually from the assemblies based on sequence GC content, read coverage and taxonomy assignment. We estimated the completeness of the assemblies with the BUSCO pipeline v5.1.2 using the Diptera database “diptera_odb10” (Seppey et al. 2019), which searches for the presence of 3285 conserved single copy Diptera orthologues.

We constructed chromosome-level reference genomes for both species by scaffolding contigs of the original assemblies with a reference guided scaffolding tool RagTag v2.1.0 (Alonge et al., 2019), which orients and orders the input contigs based on a reference using minimap2 (Li 2018). We used default settings except for the increased grouping confidence score (-i) which was increased to 0.6. For *D. montana*, we scaffolded the Seward *D. montana* assembly with the *D. lummei* genome (Poikela et al. in prep.), which was constructed using PacBio and Illumina reads and assigned to chromosomes using the published *Drosophila virilis* chromosome map (Schäfer et al. 2010) and *D. virilis* assembly dvir_r1.03_FB2015_02 obtained from Flybase. For *D. flavomontana*, we first scaffolded the Livingston *D. flavomontana* assembly with the Vancouver *D. flavomontana* assembly and then with *D. lummei*. In *D. montana* and *D. flavomontana*, chromosome 2 has right (2R) and left (2L) arms, separated by a (sub)metacentric centromere, whereas in other *virilis* group species the centromere is located near one end of the chromosome 2 (Stone et al. 1960). Therefore, scaffolding of chromosomes 2L and 2R was not feasible with the *D. lummei* genome.

For the *D. montana* chromosome-level reference genome, the X (29.1Mb), 2L (20.2Mb) and 2R (11.0Mb) chromosomes could not be further scaffolded, while the lengths of chromosomes 3, 4 and 5 were increased substantially by scaffolding. The longest contig of chromosome 3 increased from 5.8Mb to 26.0Mb (constructed from 37 contigs), chromosome 4 from 12.3Mb to 32.5Mb (28 contigs), and chromosome 5 from 19.5Mb to 26.5Mb (11 contigs; Table S10). For the *D. flavomontana* chromosome-level reference genome, the X chromosome (29.0Mb) could not be further scaffolded, while the lengths of all other chromosomes increased due to scaffolding. Chromosome 2L increased from 10.2Mb to 20.4Mb in length (3 contigs), 2R from 10.4Mb to 10.6Mb (3 contigs), the 3^rd^ chromosome from 7.8Mb to 24.5Mb (33 contigs), the 4^th^ chromosome from 20.0Mb to 30.7Mb (14 contigs), and the 5^th^ chromosome from 23.5Mb to 27.2Mb (4 contigs; Table S10).

Finally, we investigated chromosome synteny between species of the *montana* phylad (*D. montana* and *D. flavomontana;* monSE13F37 and flaMT13F11 assemblies) and *virilis* phylad (*D. virilis* and *D. lummei*) (Yusuf et al. 2022) using minimap2synteny.py (Mackintosh et al. 2023). Prior to using minimap2synteny.py, we aligned species’ assemblies using minimap2 v.2.17 (Li 2018) with the option - x asm10 and kept alignments with a mapping quality of 60.

### Genome annotations

All genome assemblies were annotated for repetitive regions and genes. *De novo* libraries of repeat sequences were built for each assembly using RepeatModeler v2.0.1 (Flynn et al. 2019), and repetitive regions were softmasked, together with *Drosophila*-specific repeats (Dfam_3.1 (Hubley et al. 2016) and RepBase-20181026 (Bao et al. 2015)), using RepeatMasker v4.1.0 (Smit et al. 2013-2015). Gene models were predicted on the softmasked assemblies of *D. montana* using the BRAKER2 pipeline. For gene annotation, we used RNA-seq data (Illumina Truseq 150b PE) from whole-body female and male *D. montana* adult flies collected in Finland (Parker et al. 2021). RNA-seq reads were trimmed for adapter contamination and read quality using fastp v0.20.0 (Chen et al. 2018), and mapped to both softmasked *D. montana* assemblies using STAR v2.7.0 (Dobin et al. 2013). Finally, *D. montana* gene annotations were carried out with BRAKER2s ab initio gene prediction pipeline with RNA-seq evidence using Augustus v3.3.3 and GeneMark-ET v4.48 (Hoff et al. 2019, 2016; Li et al. 2009; Barnett et al. 2011; Lomsadze et al. 2014; Stanke et al. 2006, 2008). Protein predictions of the Jackson *D. montana* assembly with the best BUSCO values (see the Results) were used to annotate both *D. flavomontana* and both chromosome-level genomes using the BRAKER2s ab initio gene prediction pipeline with GenomeThreader and AUGUSTUS (Hoff et al. 2016, 2019; Gremme 2013; Buchfink et al. 2015; Stanke et al. 2006, 2008). Annotation completeness was assessed using BUSCO v5.1.2 against the ‘diptera_odb10’ database (Seppey et al. 2019).

### Mapping, variant calling and variant filtering

To investigate genome-wide variation in sympatric and allopatric populations of the species, we mapped all Illumina samples to the *D. montana* chromosome-level assembly. For this, we quality-checked Illumina PE reads of each sample with FastQC v0.11.8 (Andrews 2010) and trimmed them for adapter contamination and low-quality bases using fastp v0.20.0 (Chen et al. 2018). We mapped each trimmed Illumina sample against the genome using BWA mem v0.7.17 with read group information (Li & Durbin 2009), sorted alignments with SAMtools v1.10 (Li et al. 2009) and marked PCR duplicates with sambamba v0.7.0 (Tarasov et al. 2015). The resulting BAM-files were used for variant calling with freebayes v1.3.1-dirty (Garrison & Marth 2012). Raw variants were processed with gIMble preprocess (genome-wide IM blockwise likelihood estimation toolkit; Laetsch et al. 2023). In brief, non-SNP variants were deconstructed into allelic primitives, where remaining non-SNPs were removed in addition to any SNP variant within two bases of a non-SNP. Genotype calls of remaining SNP variants were set to missing if any of the following assumptions was violated: i) sample depth (FMT/DP) between 8 and 2SD from the mean coverage, ii) read directionality placement balance (RPL>=1, RPR>=1), or iii) read strand placement balance (SAF>=1, SAR>=1).

### Principal component analysis of SNP and climatic data

To group Illumina samples according to their species and population type, we performed a principal component analysis (PCA) on the filtered VCF-file, including all samples, chromosomes, and coding, intronic and intergenic SNPs using PLINK v1.9 package (Chang et al. 2015). The VCF-file was converted to PLINK’s BED/BIM format and the PCA was run with PLINK’s --pca function.

We performed another PCA on the climatic variables at fly sampling sites to visualize the climatic variation amongst them. First, we downloaded climatic information from WorldClim database v2.1 (2.5min spatial resolution, dataset 1970-2000; Fick and Hijmans 2017) using latitudinal and longitudinal coordinates of each site (Table S1) and extracted 19 bioclimatic variables using the “raster” package v2.8-19 (Hijmans and Etten 2020; Table S5, S6). We then performed PCA on the bioclimatic variables, describing temperature and humidity conditions in each site. We performed the PCA using the “FactoMineR” package (Lê et al. 2008) in R v4.3.1 and R studio v2023.03.0.

### Characterization of chromosomal inversions

We identified large (>500kb) alternatively fixed inversions between *D. montana* and *D. flavomontana* using long- and short-read data as well as genome assemblies. We mapped PacBio reads of each sample against each of the four assemblies using ngmlr v0.2.7 (Sedlazeck et al. 2018) and obtained structural variant (SV) candidates from the SV identification software, Sniffles v1.0.12 (Sedlazeck et al. 2018). We also mapped Illumina PE reads against each of the four assemblies as explained in the “Mapping, variant calling and variant filtering” paragraph. The resulting BAM-files were given to Delly v0.8.1, which identifies structural variants based on paired-end read orientation and split-read evidence (Rausch et al. 2012). We used SURVIVOR (Jeffares et al. 2017) to identify species-specific, geographically widespread inversions that were shared by Sniffles and Delly outputs and that were found in at least in 9 *D. montana* (out of 12) and 6 *D. flavomontana* (out of 9) samples. Putative breakpoints of each inversion were located within a single contig, except for the 4^th^ chromosome inversion where breakpoints were located in two different contigs (Table S9). This inversion was therefore verified by mapping long- and short-read data against the *D. lummei* genome which has a more contiguous chromosome 4 (Table S9, Fig. S10). To determine whether the inversions belong to *D. montana* or *D. flavomontana*, we mapped PacBio reads of *D. lummei* (acting as an outgroup) against *D. montana* and *D. flavomontana* assemblies and investigated SVs using Sniffles. The putative breakpoints of the inversions were confirmed visually with Integrative Genomics Viewer (IGV) v2.8.0 (Thorvaldsdóttir et al. 2013) using both long- and short-read data (an example IGV view shown in Fig. S21).

Alternatively fixed inversions were also illustrated by aligning assemblies of *D. montana*, *D. flavomontana, D. virilis* and *D. lummei* using minimap2synteny.py (as explained in the paragraph *De novo* genome assemblies, scaffolding and chromosome synteny; Fig. 2) and nucmer alignments of the MUMmer package (Marçais et al. 2018) together with Dot plots (https://dot.sandbox.bio/; Fig. S6-S10).

Inversion breakpoints are typically named proximal and distal based on their distance from the centromere. Since there is no prior knowledge of *D. montana* and *D. flavomontana* centromeres, we identified their approximate location based on *D. virilis* chromosome maps (chromosome 2L) and genes (X: *yellow*, 4: *bl*, 5: *Cid5* and *l(2)not*) located near centromeres or telomeres (Schaeffer et al. 2008; Kursel & Malik 2017). The number of PacBio reads supporting each breakpoint as well as genes and repetitive sequences located within the 5 kb region of the breakpoints (2.5 kb flanking each side of the breakpoints) are given in Supplementary file 1.

### Modelling divergence and post-divergence gene flow

We analysed mean genetic divergence (d_xy_) and differentiation (F_st_) and fitted models of species divergence with and without long-term interspecific gene flow between and within the species using gIMble (Laetsch et al. 2023). This analytic likelihood method uses the joint distribution of mutation types in short sequence blocks, the blockwise site frequency spectrum (bSFS), across subsamples of pairs of individual genomes to fit a series of models of speciation history. We summarised data by the bSFS for two block lengths, 64b and 128b.

Given the potentially different evolutionary history of the X chromosome and the autosomes (Charlesworth et al. 2018; Vicoso & Charlesworth 2006), we ran separate analyses for them throughout. Colinear regions, ending at inversion breakpoints, were combined across autosomes as these regions are expected to share the same evolutionary history, while inversions from different chromosomes may differ in age and were analysed separately. The overlapping inversions of the X chromosome were analysed together following Counterman and Noor (2006) and Lohse et al. (2015). We analysed different chromosome partitions separately for allopatric and sympatric comparisons of the species. We also analysed the split time of *D. montana* populations living in isolation (allopatry) and in contact (sympatry) with *D. flavomontana* to evaluate the timing of potential recent introgression between the two species. The intraspecific divergence time was inferred from the colinear autosomal regions, i.e. the same data partition we used to infer the interspecific background history.

We first calculated d_xy_ and F_ST_ for different genomic regions (i.e. colinear and inverted autosomes, and colinear and inverted X chromosome) and for allopatric and sympatric populations to evaluate the role of inversions in suppressing gene exchange. These analyses were carried out separately for coding, intronic and intergenic regions (repetitive regions were excluded from all data partitions). To test whether d_xy_ and F_ST_ were increased within inversions, we simulated datasets corresponding in size to the data sampled for each inversion under the background demography (inferred from colinear autosomal regions) and compared the observed d_xy_ and F_ST_ to the distributions. We simulated inversion datasets under a minimal, conservative model of recombination, which allows for gene conversion but no cross-over. We assumed a rate of (initiation of) gene conversion of 3.59×10^−8^ per base per generation. This corresponds to recent estimates for *Drosophila pseudoobscura* and *Drosophila persimilis* (1.4×10^−5^ converted sites per base per generation; mean GC tract length of 390b) (Korunes and Noor 2018). We simulated sequences of 100kb in length, two orders of magnitude shorter than total length of intergenic sequence per inversion.

Before analysing different chromosome partitions, we investigated the likely evolutionary history of *D. montana* and *D. flavomontana* by comparing the likelihood of different demographic models. We limited this initial model selection of allopatric, sympatric and intraspecific comparisons to intergenic sequences of colinear autosomal regions (repetitive regions excluded) to minimize the effects of selection. The simplest, strict divergence (DIV) model considers isolation at time *T* without interspecific gene flow, i.e. isolation in allopatry (Fig. S17A). The isolation with migration (IM) model allows unidirectional migration rate at a constant rate *M* (Fig. S17B). The IM model was fitted to both gene flow directions (i.e. from *D. montana* to *D. flavomontana* and from *D. flavomontana* to *D. montana*, and from allopatric to sympatric *D. montana* and from sympatric to allopatric *D. montana*). The DIV and IM models allow asymmetric effective population size (N_e_) between the descendent populations, and a separate N_e_ for the ancestral population. Analyses based on the bSFS assume a constant mutation rate (μ) across blocks and no recombination within them. We assumed a mutation rate (*μ*) of 2.8 × 10^-9^ per site per generation, based on an estimate of the spontaneous mutation rate in *Drosophila melanogaster* (Keightley et al. 2014). The estimates of *T* are converted into absolute time using t = T × 2N_e_ × g, where N_e_ = θ/(4μ) and g is generation time. We assumed one generation per year, i.e. the generation time of Alaskan *D. montana* populations and most likely that of the ancestral population of the species, even though other *D. montana* and *D. flavomontana* populations presently have two generations per year (Tyukmaeva et al. 2020). To consider the potential effects of reference bias, we performed model fitting and selection twice using both *D. montana* and *D. flavomontana* chromosome-level assemblies as reference genomes.

To estimate the uncertainty in parameter estimates, i.e. the difference in support (ΔlnCL) between different demographic scenarios, we performed a parametric bootstrap. We used gIMble simulate to simulate 100 replicate datasets (of the same size as the real data in terms of numbers of blocks). To include the effect of linkage between blocks, we simulated data in 1000 chunks assuming a recombination rate of 8.9 × 10^-9^ calculated from the total map length (i.e. 1.76 × 10^-8^ divided by two given the absence of recombination in males). Specifically, we simulated data under the DIV model and fitted that model to the DIV and the best-fitting IM model to each replicate to obtain a null distribution of ΔlnCL between models (see Fig. S18 & S19).

Finally, to investigate the role of inversions in speciation, we performed demographic analyses under the best-fit model separately for different chromosome partitions (i.e. colinear and inverted autosomes, and colinear and inverted X chromosome) and for allopatric and sympatric comparisons of *D. montana* and *D. flavomontana*. The uncertainty in estimates of *T* and *M* for each data partition were inferred from 100 parametric bootstrap replicates/simulations.

### Genes putatively under divergent selection

To identify genes putatively under positive selection between *D. montana* and *D. flavomontana*, wild-caught Illumina females (Table S1) from sympatric populations were assembled with MaSuRCA v3.3.9 (Zimin et al. 2017). Furthermore, *Drosophila littoralis* female (strain ID KL13F60), collected from Korpilahti, Finland (62°00’ N; 25°34’E) in 2013 and sequenced at BGI in 2019 (details in the “*DNA extractions and sequencing”* paragraph), was assembled and used as an outgroup in the dN/dS analysis. The completeness of the assemblies was assessed using BUSCO v5.1.2 with diptera_odb10 database (Seppey et al. 2019). The genomes were annotated using protein predictions of Jackson *D. montana* PacBio assembly with the best BUSCO values (see Table S3) using BRAKER2s ab initio gene prediction pipeline with GenomeThreader and AUGUSTUS (Stanke et al. 2006, 2008; Gremme 2013; Buchfink et al. 2015; Hoff et al. 2016, 2019).

For the dN/dS analysis, we chose samples, which have originated from climatically variable populations (Fig. 3B) and obtain >97% single copy BUSCOs (Table S14). The high BUSCO values, as a proxy of high genome quality, results in a higher number of genes to be included in the analysis. Accordingly, we used *D. montana* samples from Terrace, Fall Creek, Azalea and Cranbrook and *D. flavomontana* from Terrace, Fall Creek, McBride and Cranbrook. Single-copy orthologues (SCO) between the samples were first identified with Orthofinder (v2.5.4) (Emms & Kelly 2019). The rooted phylogenetic tree produced by Orthofinder showed clear groupings of *D. montana*, *D. flavomontana* and the outgroup (Fig. S22).

The SCO proteins were aligned using Prank v.170427 and the corresponding genes codon-aligned with pal2nal v14.1. To identify genes under positive selection, we evaluated the rate of non-synonymous (dN) to synonymous (dS) substitutions (dN/dS), also known as omega (ω), across the orthologs. We used GWideCodeML (Macías et al. 2020) to run CodeML (Yang 1997) with branch-site models for all orthologs. The tree from Orthofinder was unrooted using Retree (Felsenstein 1989) and used as input for GWideCodeML. Two models were defined: the null model H_0_ (parameters model = 2, NSites = 2, fix_omega = 1 and omega = 1) that assumes no positive selection, and the alternative model H_A_ that shares the other settings of H_0_ but does not fix ω (omega = 0), allowing for optimization of this parameter. Both species were tested as being under selection. The built-in likelihood ratio tests of GWideCodeML were used to examine the orthologs, with a significantly better fit of the H_A_ model indicating the presence of positive selection.

The positively selected genes were mapped to the *D. montana* chromosome-level reference genome by extracting a representative protein sequence for each orthogroup from one randomly selected sample (flaCRAN14F7) and blasting it against the *D. montana* chromosome-level reference proteome using Diamond v2.0.15 (Buchfink et al. 2015). We blasted the genes under selection against *D. virilis* RefSeq proteins using BLASTp v2.9.0+ (Camacho et al. 2009) to obtain functional predictions for the orthologues. RefSeq protein IDs and functional predictions for the SCOs and genes putatively under divergent selection are given in Supplementary file 2. Finally, we performed a G-test to explore whether genes under divergent selection are enriched inside inversions.

## Acknowledgements

We want to thank Anna-Lotta Hiillos and Jesse Mänttäri for their help with laboratory work and Edinburgh Genomics, Norwegian Sequencing Centre and BGI for sequencing the samples. This work was supported by a grant from Academy of Finland project 322980 to MK, a grant from Finnish Cultural Foundation (Central Finland regional Fund) to NP and MK, and a grant from Jenny and Antti Wihuri Foundation to NP. KL and DRL are supported by an ERC starting grant (ModelGenomLand, 757648). KL was also supported by a Natural Environmental Research Council (NERC) UK Independent Research fellowship (NE/L011522/1). Figures 2, 3, 5, 6, S5-S10, S12, S15-S17 and S21 were finalized or produced with BioRender.com.

## Author notes

K.L., M.K. and N.P. designed the study. N.P. carried out the DNA extractions, and conducted the genome analyses with input from K.L. and D.R.L. V.H. performed the d_N_/d_S_ analysis. M.K. supervised the laboratory work and funded the study. N.P. and K.L. drafted the manuscript and all the authors helped in finalizing it.

## Data availability

Raw sequencing reads are available at SRA and genome assemblies at Genbank under BioProject PRJNA939085. Unix and R commands and Jupyter Notebooks used in the study are available in https://github.com/noorlinnea.

