## Supplementary material for "Chromosomal inversions and the demography of speciation in *Drosophila montana* and *Drosophila flavomontana*"

### Supplementary tables

Table S1. Detailed information of the samples. PacBio samples were sequenced using pools of females collected from isofemale lines that were kept in the fly laboratory since their establishment, and Illumina samples were sequenced using single wild-caught females or their $F_1$ daughters stored in -20 ˚C after the collections. Prefix "mon" and "fla" before a fly strain ID refer to *D. montana* and *D. flavomontana*, respectively.

| Collecting site | Illumina sample | Strain ID | Generation | Collecting year | PacBio |
|---|---|---|---|---|---|
| Allopatry (Alaska) | | | | | |
| Fairbanks (Moose Creek), Alaska, USA<br>64˚55'26''N; 147˚59'17''W<br>Altitude 180m | Only *D. montana* | monFA13F2 | F1 female | 2013 | |
| Honolulu (Honolulu Creek), Alaska, USA<br>63˚03'49''N; 149˚32'40''W<br>Altitude 495 m | Only *D. montana* | monHO13F3 | F1 female | 2013 | |
| Seward, Alaska, USA<br>60˚09'45''N; 149˚27'09''W<br>Altitude 35 m | Only *D. montana* | monSE13F37 | F1 female | 2013 | 60 females from the isofemale strain |
| Sympatry (Rocky Mountains) | | | | | |
| McBride, British Columbia, Canada<br>53˚07'N; 120˚18'W<br>Altitude 720 m | Both species | monMB14F1<br>flaMB14F10 | Founder female<br>Founder female | 2014<br>2014 | |
| Cranbrook, British Columbia, Canada<br>49˚36'N; 115˚46'W<br>Altitude 940 m | Both species | monCRAN14F16<br>flaCRAN14F7 | Founder female<br>Founder female | 2014<br>2014 | |
| Livingston, Montana, USA<br>45˚20'59''N; 110˚36'30''W<br>Altitude 1 605 m | Only *D. flavomontana* | flaMT13F11 | F1 female | 2013 | 60 females from the isofemale strain |
| Jackson, Wyoming, USA<br>43˚25'48''N; 110˚50'34''W<br>Altitude 1 857 m | Both species | monJX13F3<br>flaJX13F38 | F1 female<br>F1 female | 2013<br>2013 | 60 females from the isofemale strain |
| Afton, Wyoming, USA<br>42˚43'N; 110˚55'W<br>Altitude 2 000 m | Only *D. montana* | monAF15F28 | F1 female | 2015 | |
| Liberty, Utah, USA<br>41˚20'N; 111˚51'W<br>Altitude 1 600 m | Only *D. flavomontana* | flaLB15F3 | F1 female | 2015 | |
| Sympatry (Western coast) | | | | | |
| Terrace, British Columbia, Canada<br>54˚27'N; 128˚34'W<br>Altitude 217 m | Both species | monTER14F11<br>flaTER14F5 | Founder female<br>Founder female | 2014<br>2014 | |
| Vancouver, British Columbia, Canada<br>49°15'N;123°10'W<br>Altitude 4 m | Both species | monVAN14F1<br>flaVAN14F20 | Founder female<br>Founder female | 2014<br>2014 | <br>60 females from the isofemale strain |
| Ashford, Washington, Usa<br>46˚45'21''N; 121˚57'22''W<br>Altitude 573 m | Both species | monASH13F13<br>flaASH13F2 | F1 female<br>F1 female | 2013<br>2013 | |
| Fall Creek, Oregon, USA<br>43˚58'32''N; 122˚45'06''W<br>Altitude 225 m | Both species | mon4Fall<br>fla3Fall | F1 female<br>F1 female | 2010<br>2010 | |
| Azalea, Oregon, USA<br>42˚48'24''N; 123˚13'37''W<br>Altitude 498 m | Only *D. montana* | monAZA2 | F1 female | 2010 | |

Table S2. PacBio and Illumina raw reads. N50 raw reads = half of the raw reads are larger than or equal to the N50 raw read value.

| PacBio reads | | | | Total number of reads | | | | | | Number of reads | | | |
|---|---|---|---|---|---|---|---|---|---|---|---|---|---|
| **Species** | **Region** | **Population** | **Strain ID** | **SMRT 1** | **SMRT 2** | **Mean coverage** | **Max read length (bp)** | **Average read length (bp)** | **N50 raw read (bp)** | **>= 80 000 bp** | **Bases (Gb)** | **Technique** | **Sequencing facility** |
| *D. montana* | Allopatry (Alaska) | Seward | monSE13F37 | 888,644 | 781,439 | 35.1 | 168303 | 5905.9 | 11366 | 81 | 9.9 | PacBio Sequel | Norwegian sequencing center |
| | Sympatry (Rocky Mountains) | Jackson | monJX13F48 | 2,161,616 | - | 62.9 | 111203 | 6422.3 | 8045 | 19 | 13.9 | PacBio Sequel | BGI |
| *D. flavomontana* | Sympatry (Rocky Mountains) | Livingston | flaMT13F11 | 684,309 | 758,824 | 27.1 | 194838 | 5390.7 | 10773 | 121 | 7.8 | PacBio Sequel | Norwegian sequencing center |
| | Sympatry (Western coast) | Vancouver | flaVAN14F20 | 710,709 | 839,748 | 30.9 | 169754 | 5861.8 | 12235 | 133 | 9.1 | PacBio Sequel | Norwegian sequencing center |
| **Illumina 150bp paired-end reads** | | | | **Total number of reads** | | | | | | | | | |
| **Species** | **Region** | **Population** | **Strain ID** | **Lane 1** | **Lane 2** | **Mean coverage** | **Insert size peak (bp)** | | | | | **Technique** | **Sequencing facility** |
| *D. montana* | Allopatry (Alaska) | Seward | monSE13F37 | 315,326,680 | - | 434.6 | 261 | | | | | HiSeq4000 | Norwegian sequencing center |
| | | Honolulu Creek | monHO13F3 | 48,321,573 | - | 72.7 | 255 | | | | | HiSeq X-Ten | BGI |
| | | Fairbanks | monFA13F2 | 52,970,348 | - | 79.9 | 251 | | | | | HiSeq X-Ten | BGI |
| | Sympatry (Rocky Mountains) | McBride | monMB14F1 | 64,726,992 | - | 52.9 | 229 | | | | | HiSeq X-Ten | BGI |
| | | Cranbrook | monCAN14F16 | 53,005,731 | - | 75.2 | 257 | | | | | HiSeq X-Ten | BGI |
| | | Jackson | monJX13F48 | 33,029,719 | 31,973,224 | 81.9 | 150 | | | | | HiSeq4000 | Edinburgh Genomics |
| | | Afton | monAF15F28 | 34,605,493 | 36,274,368 | 84.8 | 150 | | | | | HiSeq4000 | Edinburgh Genomics |
| | Sympatry (Western coast) | Terrace | monTER14F11 | 48,422,463 | - | 66.6 | 239 | | | | | HiSeq X-Ten | BGI |
| | | Vancouver | monVAN14F1 | 30,641,929 | 30,099,059 | 69.2 | 150 | | | | | HiSeq4000 | Edinburgh Genomics |
| | | Ashford | monASH13F13 | 42,182,222 | 41,678,505 | 93.9 | 111 | | | | | HiSeq4000 | Edinburgh Genomics |
| | | Fall Creek | mon4Fall | 55,597,271 | - | 81.7 | 261 | | | | | HiSeq X-Ten | BGI |
| | | Azalea | monAZA2 | 47,287,149 | - | 70.2 | 250 | | | | | HiSeq X-Ten | BGI |
| *D. flavomontana* | Sympatry (Rocky Mountains) | McBride | flaMB14F10 | 49,610,911 | - | 70.5 | 262 | | | | | HiSeq X-Ten | BGI |
| | | Cranbrook | flaCRAN14F7 | 54,276,116 | - | 70.6 | 258 | | | | | HiSeq X-Ten | BGI |
| | | Livingston | flaMT13F11 | 28,832,333 | 30,942,904 | 74.6 | 150 | | | | | HiSeq4000 | Edinburgh Genomics |
| | | Jackson | flaJX13F31 | 41,741,562 | 42,330,188 | 109.1 | 150 | | | | | HiSeq4000 | Edinburgh Genomics |
| | Sympatry (Western coast) | Liberty | flaLB15F3 | 39,780,753 | 40,656,195 | 101.5 | 150 | | | | | HiSeq4000 | Edinburgh Genomics |
| | | Terrace | flaTER14F5 | 50,162,958 | - | 71.9 | 150 | | | | | HiSeq X-Ten | BGI |
| | | Vancouver | flaVAN14F20 | 36,923,245 | 38,165,991 | 91.7 | 150 | | | | | HiSeq4000 | Edinburgh Genomics |
| | | Ashford | flaASH13F2 | 23,396,198 | 22,877,448 | 58.6 | 150 | | | | | HiSeq4000 | Edinburgh Genomics |
| | | Fall Creek | fla3Fall | 50,824,988 | - | 73.0 | 150 | | | | | HiSeq X-Ten | BGI |

Table S3. Contiguity, completeness, and annotation of original and scaffolded chromosome-level genome assemblies. N50 = half of the genome is in contigs larger than or equal to the N50 contig size. N50 count = half of the genome consists of the number indicated in N50 count.

| | *D. montana* Seward, Alaska monSE13F37 | *D. montana* Jackson, USA monJX13F48 | *D. flavomontana* Livingston, USA flaMT13F11 | *D. flavomontana* Vancouver, Canada flaVAN14F20 | *D. montana* chromosome-level genome | *D. flavomontana* chromosome-level genome |
|---|---|---|---|---|---|---|
| **Genome contiguity** | | | | | | |
| **Genome size (Mb)** | 184.3 | 181.0 | 192.6 | 193.5 | 145.5 | 148.1 |
| **Total contigs** | 324 | 796 | 376 | 296 | 6 | 6 |
| **Longest contig (Mb)** | 29.1 | 15.2 | 29.0 | 26.8 | 32.5 | 31.2 |
| **N50 (Mb)** | 11.0 | 1.3 | 9.8 | 7.8 | 26.5 | 28.8 |
| **N50 count** | 5 | 20 | 6 | 6 | 3 | 3 |
| **Genome completeness** | | | | | | |
| **Complete BUSCOs (%; n: 3285)** | 98.1 | 98.5 | 97.5 | 97.3 | 91.9 | 93.5 |
| **Single copy BUSCOs (%)** | 97.6 | 98.0 | 97.1 | 96.9 | 91.7 | 93.2 |
| **Duplicated BUSCOs (%)** | 0.5 | 0.5 | 0.4 | 0.4 | 0.2 | 0.3 |
| **Genome annotation** | | | | | | |
| **GC level (%)** | 40.2 | 40.2 | 40.2 | 40.1 | 40.3 | 40.3 |
| **Masked repeats (%)** | 26.6 | 25.5 | 29.9 | 30.4 | 16.8 | 16.3 |
| **Genes (N)** | 15,696 | 15,819 | 15,940 | 16,056 | 15,446 | 15,299 |
| **Mean gene length (bp)** | 1,640 | 1,637 | 1,568 | 1,564 | 1,571 | 1,566 |
| **Complete BUSCOs (%; n: 3285)** | 94.2 | 94.7 | 92.5 | 92.9 | 88.5 | 89.7 |
| **Single copy BUSCOs (%)** | 84.4 | 84.5 | 84.1 | 84.5 | 79.9 | 81.3 |
| **Duplicated BUSCOs (%)** | 9.8 | 10.2 | 8.4 | 8.4 | 8.8 | 8.4 |

Table S4. Principal components, Eigenvalues, variance, and cumulative variance of 12 *D. montana* and 9 *D. flavomontana* Illumina samples.

| PC | Eigenvalue | Variance (%) | Cumulative variance (%) |
|---|---|---|---|
| PC1 | 14.0 | 50.4 | 50.4 |
| PC2 | 1.1 | 4.0 | 54.5 |
| PC3 | 1.0 | 3.7 | 58.1 |
| PC4 | 1.0 | 3.5 | 61.6 |
| PC5 | 0.9 | 3.4 | 65.1 |
| PC6 | 0.9 | 3.2 | 68.3 |
| PC7 | 0.9 | 3.2 | 71.5 |
| PC8 | 0.9 | 3.1 | 74.6 |
| PC9 | 0.9 | 3.1 | 77.7 |
| PC10 | 0.9 | 3.1 | 80.8 |
| PC11 | 0.9 | 3.1 | 83.9 |
| PC12 | 0.8 | 3.0 | 86.9 |
| PC13 | 0.7 | 2.4 | 89.3 |
| PC14 | 0.5 | 1.8 | 91.1 |
| PC15 | 0.5 | 1.6 | 92.7 |
| PC16 | 0.4 | 1.5 | 94.3 |
| PC17 | 0.4 | 1.5 | 95.7 |
| PC18 | 0.4 | 1.5 | 97.2 |
| PC19 | 0.4 | 1.4 | 98.6 |
| PC20 | 0.4 | 1.4 | 100.0 |

Table S5. A List of 19 bioclimatic variables used in the PCA (WorldClim database v2.0, 2.5 min spatial resolution; dataset 1970-2000; www.worldclim.org) (Fick and Hijmans 2017).

| Variable | Description |
|---|---|
| bio1 | Annual mean temperature |
| bio2 | Mean diurnal range (mean of monthly (max - min temperature)) |
| bio3 | Isothermality (bio2/bio7*100) |
| bio4 | Temperature seasonality (standard deviation * 100) |
| bio5 | Max temperature of the warmest month |
| bio6 | Min temperature of the coldest month |
| bio7 | Annual temperature range (bio5-bio6) |
| bio8 | Mean temperature of the wettest quarter |
| bio9 | Mean temperature of the driest quarter |
| bio10 | Mean temperature of the warmest quarter |
| bio11 | Mean temperature of the coldest quarter |
| bio12 | Annual precipitation |
| bio13 | Precipitation of the wettest month |
| bio14 | Precipitation of the driest month |
| bio15 | Precipitation seasonality (coefficient of variation) |
| bio16 | Precipitation of the wettest quarter |
| bio17 | Precipitation of the driest quarter |
| bio18 | Precipitation of the warmest quarter |
| bio19 | Precipitation of the coldest quarter |

Table S6. 19 bioclimatic variables (explanations are in Table S5) for each site extracted from WorldClim database v2.1 using latitudinal and longitudinal coordinates (2.5 min spatial resolution; dataset 1970-2000; www.worldclim.org) (Fick and Hijmans 2017).

| Population | Longitude | Altitude | Latitude | bio1 | bio2 | bio3 | bio4 | bio5 | bio6 | bio7 | bio8 | bio9 | bio10 | bio11 | bio12 | bio13 | bio14 | bio15 | bio16 | bio17 | bio18 | bio19 |
|---|---|---|---|---|---|---|---|---|---|---|---|---|---|---|---|---|---|---|---|---|---|---|
| Fairbanks | -147.99 | 180.00 | 64.92 | -2.84 | 11.05 | 23.49 | 1371.73 | 21.74 | -25.33 | 47.06 | 14.00 | -10.18 | 14.00 | -19.01 | 304 | 49 | 8 | 52 | 135 | 33 | 135 | 50 |
| Honolulu Creek | -149.54 | 495.00 | 63.06 | -1.22 | 9.57 | 26.43 | 1027.63 | 19.13 | -17.09 | 36.22 | 10.26 | -1.91 | 12.16 | -12.49 | 636 | 103 | 25 | 50 | 281 | 88 | 251 | 107 |
| Seward | -149.54 | 35.00 | 60.16 | 0.68 | 6.54 | 29.29 | 605.24 | 13.48 | -8.84 | 22.32 | 0.65 | 2.64 | 8.74 | -5.60 | 1104 | 162 | 52 | 42 | 423 | 161 | 210 | 302 |
| Terrace | -128.57 | 217.00 | 54.45 | 6.41 | 7.72 | 27.57 | 730.96 | 21.38 | -6.63 | 28.01 | 1.75 | 13.43 | 15.32 | -2.71 | 1379 | 210 | 50 | 52 | 578 | 157 | 170 | 487 |
| Vancouver | -122.60 | 4.00 | 49.24 | 10.04 | 9.15 | 36.73 | 577.06 | 24.20 | -0.70 | 24.90 | 3.52 | 17.02 | 17.09 | 3.06 | 1972 | 308 | 64 | 49 | 818 | 221 | 226 | 691 |
| Ashford | -121.96 | 573.00 | 46.76 | 6.25 | 9.40 | 39.81 | 512.91 | 20.72 | -2.88 | 23.60 | 1.02 | 12.65 | 13.00 | 0.69 | 2333 | 361 | 48 | 58 | 1057 | 199 | 213 | 964 |
| Fall Creek | -122.75 | 225.00 | 43.98 | 10.93 | 11.96 | 45.11 | 524.28 | 27.29 | 0.78 | 26.51 | 5.27 | 17.47 | 17.70 | 5.00 | 1497 | 233 | 20 | 61 | 676 | 111 | 112 | 609 |
| Azalea | -123.23 | 498.00 | 42.81 | 11.05 | 13.74 | 48.03 | 555.79 | 28.90 | 0.30 | 28.60 | 5.03 | 18.11 | 18.20 | 4.80 | 1145 | 196 | 13 | 70 | 559 | 62 | 71 | 507 |
| McBride | -120.16 | 720.00 | 53.30 | 4.07 | 11.31 | 32.24 | 850.10 | 22.13 | -12.94 | 35.07 | 4.04 | 0.10 | 14.23 | -6.66 | 706 | 75 | 39 | 21 | 210 | 127 | 201 | 163 |
| Cranbrook | -115.74 | 940.00 | 49.52 | 5.55 | 12.24 | 32.16 | 899.03 | 25.95 | -12.10 | 38.06 | 14.11 | 0.93 | 16.50 | -5.85 | 461 | 61 | 26 | 27 | 151 | 85 | 133 | 113 |
| Livingston | -110.61 | 1605.00 | 45.36 | 5.13 | 14.06 | 38.05 | 836.35 | 25.90 | -11.05 | 36.95 | 12.93 | -4.85 | 15.66 | -4.85 | 500 | 72 | 24 | 35 | 185 | 83 | 154 | 83 |
| Jackson | -110.84 | 1875.00 | 43.43 | 3.51 | 15.73 | 38.05 | 924.38 | 26.55 | -14.80 | 41.35 | 8.07 | -1.79 | 14.87 | -8.04 | 456 | 52 | 28 | 20 | 132 | 94 | 110 | 120 |
| Afton | -110.92 | 2000.00 | 42.72 | 3.14 | 16.64 | 36.87 | 1030.70 | 27.49 | -17.63 | 45.12 | 8.47 | 15.39 | 15.39 | -10.24 | 466 | 53 | 30 | 16 | 140 | 104 | 104 | 112 |
| Liberty | -111.89 | 1600.00 | 41.33 | 5.95 | 13.73 | 36.51 | 866.64 | 27.02 | -10.59 | 37.61 | 4.65 | 16.83 | 16.83 | -4.49 | 515 | 55 | 27 | 21 | 162 | 89 | 89 | 129 |

Table S7. Principal components, Eigenvalues, variance, and cumulative variance of bioclimatic variables.

| PC | Eigenvalue | Variance (%) | Cumulative variance (%) |
|---|---|---|---|
| PC1 | 10.44 | 54.93 | 54.93 |
| PC2 | 5.11 | 26.88 | 81.80 |
| PC3 | 1.43 | 7.55 | 89.35 |
| PC4 | 0.71 | 3.76 | 93.11 |
| PC5 | 0.54 | 2.86 | 95.98 |
| PC6 | 0.46 | 2.44 | 98.42 |
| PC7 | 0.14 | 0.76 | 99.17 |
| PC8 | 0.10 | 0.51 | 99.68 |
| PC9 | 0.05 | 0.24 | 99.92 |
| PC10 | 0.01 | 0.04 | 99.97 |
| PC11 | 0.00 | 0.02 | 99.98 |
| PC12 | 0.00 | 0.01 | 100.00 |
| PC13 | 0.00 | 0.00 | 100.00 |

Table S8. Contributions (loadings) of the bioclimatic variables (see also Table S5-S6) on each principal component (PC).

| Variable | PC1 | PC2 | PC3 | PC4 | PC5 |
|---|---|---|---|---|---|
| bio1 | 4.89 | 8.34 | 0.59 | 0.29 | 5.21 |
| bio2 | 2.19 | 11.82 | 3.21 | 0.29 | 0.50 |
| bio3 | 2.25 | 11.84 | 0.04 | 4.12 | 0.93 |
| bio4 | 8.23 | 0.39 | 0.67 | 0.15 | 6.01 |
| bio5 | 0.10 | 18.16 | 0.38 | 7.42 | 0.10 |
| bio6 | 8.15 | 1.87 | 0.01 | 2.70 | 5.19 |
| bio7 | 8.24 | 0.87 | 0.19 | 8.88 | 3.99 |
| bio8 | 5.88 | 0.82 | 6.64 | 2.34 | 19.88 |
| bio9 | 4.75 | 4.00 | 3.41 | 0.01 | 27.41 |
| bio10 | 0.14 | 15.23 | 0.05 | 5.93 | 1.23 |
| bio11 | 7.30 | 3.59 | 0.14 | 2.11 | 6.41 |
| bio12 | 8.78 | 0.38 | 0.62 | 4.68 | 1.12 |
| bio13 | 8.60 | 0.29 | 2.60 | 4.66 | 0.89 |
| bio14 | 3.68 | 5.16 | 20.32 | 4.30 | 0.51 |
| bio15 | 3.12 | 0.03 | 44.66 | 0.04 | 0.98 |
| bio16 | 8.50 | 0.19 | 3.23 | 4.07 | 1.75 |
| bio17 | 5.88 | 3.19 | 11.31 | 7.15 | 0.06 |
| bio18 | 0.74 | 13.84 | 0.03 | 7.23 | 14.33 |
| bio19 | 8.57 | 0.00 | 1.89 | 3.62 | 3.51 |

Table S9. Genomic coordinates of alternatively fixed chromosomal inversions of *D. montana* and *D. flavomontana*, based on the original assemblies of the species as well as the chromosome-level genomes of both species (Table S3). Each of the five alternatively fixed inversions as well as one putative heterozygous inversion are shown in their own line. The breakpoints are named proximal and distal based on their distance to the centromere (the approximate locations of the centromeres were identified using *D. virilis* chromosome maps and genes, see Methods for details).

| | | **Inversion breakpoints in *D. montana* and *D. flavomontana* genomes** | | | | | **Inversion breakpoints in *D. montana* chromosome-level genome** | | | | | **Inversion breakpoints in *D. flavomontana* chromosome-level genome** | | | | | |
|---|---|---|---|---|---|---|---|---|---|---|---|---|---|---|---|---|---|
| | | | | **Breakpoint** | | | | | **Breakpoint** | | | | | **Breakpoint** | | | |
| **Species the inversion belongs to** | **Chromosome** | **Contig** | **Contig length (bp)** | **Distal** | **Proximal** | **Size (bp)** | **Scaffold** | **Scaffold length** | **Distal** | **Proximal** | **Size (bp)** | **Scaffold** | **Scaffold length** | **Distal** | **Proximal** | **Size (bp)** | **Representation in Illumina samples** |
| *D. montana* | X | monSE13F37.00001 | 29140820 | 3993916 | 11173058 | 7179142 | X_chromosome | 29140820 | 3993916 | 11173058 | 7179143 | X_chromosome | 28975156 | 6309450 | 20284519 | 13975070 | homozygous in all *D. montana* samples |
| | | monJX13F48.00118, monJX13F48.00006 | 253764, 4625534 | 104108 | 1345305 | - | | | | | | | | | | | |
| | | flaMT13F11.00001 | 28975156 | 6309450 | 20284519 | 13975069 | | | | | | | | | | | |
| | | flaVAN14F20.00001 | 26778568 | 22735439 | 8730382 | 14005057 | | | | | | | | | | | |
| *D. flavomontana* | X | monSE13F37.00001 | 28975156 | 6041987 | 20201805 | 14159819 | X_chromosome | 29140820 | 6041987 | 20201805 | 14159819 | X_chromosome | 28975156 | 11363874 | 22321964 | 10958091 | homozygous in all *D. montana* samples |
| | | monJX13F48.00021, monJX13F48.00002 | 1229388, 13454835 | 142107 | 6726561 | - | | | | | | | | | | | |
| | | flaMT13F11.00001 | 28975156 | 11363874 | 22321964 | 10958091 | | | | | | | | | | | |
| | | flaVAN14F20.00001 | 26778568 | 17674592 | 6694000 | 10980592 | | | | | | | | | | | |
| *D. flavomontana* | X | monSE13F37.00001 | 29140820 | 17044685 | 20200540 | 3155884 | X_chromosome | 29140820 | 17044685 | 20200540 | 3155884 | X_chromosome | 28975156 | 11368068 | 14486661 | 3118594 | homozygous in all *D. flavomontana* samples |
| | | monJX13F48.00021, monJX13F48.00002 | 1229388, 13454835 | 144296 | 9907081 | - | | | | | | | | | | | |
| | | flaMT13F11.00001 | 28975156 | 11368068 | 14486661 | 3119573 | | | | | | | | | | | |
| | | flaVAN14F20.00001 | 26778568 | 17672247 | 14539822 | 3132425 | | | | | | | | | | | |
| *D. montana* | 2L | monSE13F37.00002 | 20245637 | 15098446 | 19046306 | 3947861 | 2L_chromosome | 20245637 | 15098446 | 19046306 | 3947861 | 2L_chromosome | 20392248 | 1181134 | 5145967 | 3964834 | heterozygous in all *D. montana* samples |
| | | monJX13F48.00003 | 9138818 | 8048854 | 4117707 | 3931148 | | | | | | | | | | | |
| | | flaMT13F11.00005 | 10178141 | 1181134 | 5145967 | 3964833 | | | | | | | | | | | |
| | | flaVAN14F20.00004, flaVAN14F20.00047 | 17678878, 659972 | 2535965 | 477262 | - | | | | | | | | | | | |
| *D. flavomontana* | 4 | monSE13F37.00004, monSE13F37.00009 | 12274228, 2530810 | 10263367 | 1906955 | - | 4_chromosome | 32544174 | 17382172 | 3246708 | 14135465 | 4_chromosome | 30698533 | 23745872 | 7855585 | 15890288 | homozygous in all *D. flavomontana* samples, |
| Named J in Stone et al. 1960 | | monJX13F48.00036, monJX13F48.00073 | 596273, 350243 | 226846 | 284790 | - | | | | | | | | | | | |
| | | flaMT13F11.00003, flaMT13F11.00008 | 19965754, 5486619 | 12355487 | 1949285 | - | | | | | | | | | | | |
| | | flaVAN14F20.00003, flaVAN14F20.00008 | 19125027, 4169626 | 12375436 | 1985368 | - | | | | | | | | | | | |
| | | lummei.00004* | 18946740 | 1568294 | 15689984 | 14121690 | | | | | | | | | | | |
| *D. flavomontana* | 5 | monSE13F37.00003 | 20245037 | 7861561 | 17088002 | 9226442 | 5_chromosome | 26508887 | 9422591 | 18649749 | 9227159 | 5_chromosome | 27217941 | 21134310 | 11949884 | 9184427 | homozygous in all *D. flavomontana* samples |
| Named E in Stone et al. 1960 | | flaMT13F11.00002 | 23523717 | 17439995 | 8255383 | 9183122 | | | | | | | | | | | |
| | | flaVAN14F20.00002 | 22345759 | 16280386 | 7084336 | 9194450 | | | | | | | | | | | |
| | | monJX13F48.00001 | 15164525 | 11733609 | 2455274 | 9278336 | | | | | | | | | | | |

* *D. flavomontana* inversion breakpoints on the 4th chromosome are found in different contigs in *D. montana* and *D. flavomontana* assemblies, and thus the inversion was verified using *D. lummei* genome.

Table S10. Contigs of each assembly (>1Mb contigs shown here) were assigned to chromosomes using *Drosophila lummei* genome (Poikela et al. in prep.).

| **Seward *D. montana*** | | | **Jackson *D. montana*** | | | **Livingston *D. flavomontana*** | | | **Vancouver *D. flavomontana*** | | |
|---|---|---|---|---|---|---|---|---|---|---|---|
| **Contig length (Mb)** | | **Chromosome** | **Contig length (Mb)** | | **Chromosome** | **Contig length (Mb)** | | **Chromosome** | **Contig length (Mb)** | | **Chromosome** |
| monSE13F37.00001 | 29140820 | X | monJX13F48.00001 | 15164525 | 5 | flaMT13F11.00001 | 28975156 | X | flaVAN14F20.00001 | 26778568 | X |
| monSE13F37.00002 | 20245637 | 2L | monJX13F48.00002 | 13454835 | X | flaMT13F11.00002 | 23523717 | 5 | flaVAN14F20.00002 | 22345759 | 5 |
| monSE13F37.00003 | 19534505 | 5 | monJX13F48.00003 | 9138818 | 2L | flaMT13F11.00003 | 19965754 | 4 | flaVAN14F20.00003 | 19125027 | 4 |
| monSE13F37.00004 | 12274228 | 4 | monJX13F48.00004 | 8789853 | 2L | flaMT13F11.00004 | 10393190 | 2R | flaVAN14F20.00004 | 17678878 | 2L |
| monSE13F37.00005 | 10997687 | 2R | monJX13F48.00005 | 8412689 | 3 | flaMT13F11.00005 | 10178141 | 2L | flaVAN14F20.00005 | 10603604 | 2R |
| monSE13F37.00006 | 5767499 | 3 | monJX13F48.00006 | 4625534 | X | flaMT13F11.00006 | 9776843 | 2L | flaVAN14F20.00006 | 7750958 | 3 |
| monSE13F37.00007 | 3775224 | 3 | monJX13F48.00007 | 3696986 | 5 | flaMT13F11.00007 | 7754770 | 3 | flaVAN14F20.00007 | 4798025 | 3 |
| monSE13F37.00008 | 2568140 | 4 | monJX13F48.00008 | 3308613 | 2R | flaMT13F11.00008 | 5486619 | 4 | flaVAN14F20.00008 | 4169626 | 4 |
| monSE13F37.00009 | 2530810 | 4 | monJX13F48.00009 | 3289462 | 4 | flaMT13F11.00009 | 3073694 | 3 | flaVAN14F20.00009 | 3314633 | 3 |
| monSE13F37.00010 | 2178246 | 3 | monJX13F48.00010 | 3232620 | - | flaMT13F11.00010 | 2473142 | 3 | flaVAN14F20.00010 | 2593425 | 3 |
| monSE13F37.00011 | 1979467 | 5 | monJX13F48.00011 | 2813930 | 2R | flaMT13F11.00011 | 2460368 | 3 | flaVAN14F20.00011 | 2245353 | 2L |
| monSE13F37.00012 | 1726930 | 4 | monJX13F48.00012 | 2122920 | 2R | flaMT13F11.00012 | 2261038 | 3 | flaVAN14F20.00012 | 2071253 | 4 |
| monSE13F37.00013 | 1610333 | - | monJX13F48.00013 | 2080494 | 2R | flaMT13F11.00013 | 2022541 | 3 | flaVAN14F20.00013 | 1841750 | 3 |
| monSE13F37.00014 | 1585447 | 4 | monJX13F48.00014 | 1980436 | 5 | flaMT13F11.00014 | 1824522 | - | flaVAN14F20.00014 | 1714737 | 4 |
| monSE13F37.00015 | 1444893 | 3 | monJX13F48.00015 | 1656297 | 3 | flaMT13F11.00015 | 1668200 | 3 | flaVAN14F20.00015 | 1374044 | - |
| monSE13F37.00016 | 1385966 | 3 | monJX13F48.00016 | 1503212 | 4 | flaMT13F11.00016 | 1643239 | 2L | flaVAN14F20.00016 | 1368565 | - |
| monSE13F37.00017 | 1362064 | 4 | monJX13F48.00017 | 1455675 | 3 | flaMT13F11.00017 | 1376368 | - | flaVAN14F20.00017 | 1336510 | X |
| monSE13F37.00018 | 1364004 | - | monJX13F48.00018 | 1408546 | - | flaMT13F11.00018 | 1364617 | - | flaVAN14F20.00018 | 1295148 | - |
| monSE13F37.00019 | 1327839 | - | monJX13F48.00019 | 1365292 | 4 | flaMT13F11.00019 | 1346930 | 2L | flaVAN14F20.00019 | 1292286 | - |
| monSE13F37.00020 | 1290151 | - | monJX13F48.00020 | 1284102 | - | flaMT13F11.00020 | 1305735 | - | flaVAN14F20.00020 | 1278608 | 2L |
| monSE13F37.00021 | 1271459 | 4 | | | | flaMT13F11.00021 | 1205103 | - | flaVAN14F20.00021 | 1168480 | - |
| monSE13F37.00022 | 1212424 | - | | | | flaMT13F11.00022 | 1163332 | - | flaVAN14F20.00022 | 1144968 | 3 |
| monSE13F37.00023 | 1188514 | 3 | | | | flaMT13F11.00023 | 1128347 | 4 | flaVAN14F20.00023 | 1099907 | 3 |
| monSE13F37.00024 | 1154756 | 4 | | | | flaMT13F11.00024 | 1089159 | - | flaVAN14F20.00024 | 1085779 | - |
| monSE13F37.00025 | 1124584 | - | | | | | | | flaVAN14F20.00025 | 1074364 | - |
| monSE13F37.00026 | 1122473 | - | | | | | | | flaVAN14F20.00026 | 1042346 | - |
| monSE13F37.00027 | 1100476 | X | | | | | | | | | |
| monSE13F37.00028 | 1088638 | 3 | | | | | | | | | |
| monSE13F37.00029 | 1086808 | 4 | | | | | | | | | |
| monSE13F37.00030 | 1052805 | 3 | | | | | | | | | |
| monSE13F37.00031 | 1043766 | - | | | | | | | | | |

Table S11. Summary of the effects of chromosome partition (colinear vs inverted) on $d_{xy}$ and $F_{st}$ in allopatric and sympatric populations of *D. montana* and *D. flavomontana* and in intraspecific comparisons within *D. montana*. $d_{xy}$ and $F_{st}$ was analysed separately for the autosomes and the X chromosome and for intergenic, intronic and coding sequences. P-values were inferred from simulations and significant P-values are in bold.

| Genomic region | Comparison | Comparison | $d_{xy}$ | P-value | $F_{st}$ | P-value |
|---|---|---|---|---|---|---|
| **Intergenic** | **Allopatry** | [Colinear autosomes] | 0.03030 | | 0.5968 | |
| | | 4 inverted | 0.03511 | **< 0.001** | 0.6072 | 0.656 |
| | | 5 inverted | 0.03113 | **< 0.001** | 0.6200 | 0.278 |
| | | [Colinear X] | 0.03211 | | 0.7021 | |
| | | X inverted | 0.03505 | **< 0.001** | 0.7273 | 0.942 |
| | **Sympatry** | [Colinear autosomes] | 0.03004 | | 0.5857 | |
| | | 4 inverted | 0.03495 | **< 0.001** | 0.5930 | 0.752 |
| | | 5 inverted | 0.03092 | **< 0.001** | 0.6110 | 0.290 |
| | | [Colinear X] | 0.03194 | | 0.6876 | |
| | | X inverted | 0.03484 | **< 0.001** | 0.6847 | 0.050 |
| **Introns** | **Allopatry** | [Colinear autosomes] | 0.03127 | | 0.6042 | |
| | | 4 inverted | 0.03676 | **< 0.001** | 0.6114 | 0.596 |
| | | 5 inverted | 0.03333 | **< 0.001** | 0.6160 | 0.494 |
| | | [Colinear X] | 0.03510 | | 0.7191 | |
| | | X inverted | 0.03824 | **< 0.001** | 0.7233 | 0.928 |
| | **Sympatry** | [Colinear autosomes] | 0.03102 | | 0.5959 | |
| | | 4 inverted | 0.03663 | **< 0.001** | 0.5919 | 0.190 |
| | | 5 inverted | 0.03312 | **0.018** | 0.6083 | 0.532 |
| | | [Colinear X] | 0.03498 | | 0.7052 | |
| | | X inverted | 0.03799 | **< 0.001** | 0.6798 | **0.034** |
| **Coding** | **Allopatry** | [Colinear autosomes] | 0.02025 | | 0.6472 | |
| | | 4 inverted | 0.02206 | **< 0.001** | 0.6636 | 0.534 |
| | | 5 inverted | 0.01997 | 0.634 | 0.6405 | 0.208 |
| | | [Colinear X] | 0.01694 | | 0.7570 | |
| | | X inverted | 0.01945 | **< 0.001** | 0.7400 | 0.104 |
| | **Sympatry** | [Colinear autosomes] | 0.02014 | | 0.6394 | |
| | | 4 inverted | 0.02203 | **< 0.001** | 0.6472 | 0.654 |
| | | 5 inverted | 0.01989 | 0.516 | 0.6330 | 0.202 |
| | | [Colinear X] | 0.01678 | | 0.7494 | |
| | | X inverted | 0.01925 | **< 0.001** | 0.7078 | 0.064 |

Table S12. Support (measured as ΔlnCL) and parameter estimates for divergence time (T), migration rate (M) and effective population sizes ($N_e$) for studied populations and their common ancestral population under strict divergence (DIV, M=0) and isolation with migration (IM) models with both gene flow directions. This initial model comparison was performed for intergenic autosomal colinear regions to minimize the effects of selection, using two reference genomes (*D. montana* and *D. flavomontana* ) and two block sizes (64b and 128b). Grey shading indicates the best-fit model for each comparison (see Fig. S18-S19).

| **Reference genome used** | **Block size** | **Comparison** | **Model** | ***D. mon* Ne** | ***D. fla* Ne** | **Ancestral Ne** | **T** | **m** | **lnCL** | **ΔlnCL** |
|---|---|---|---|---|---|---|---|---|---|---|
| *D. montana* | 64 bp | Allopatric comparison | DIV | 692600 | 395250 | 1464316 | 2378570 | - | -45651205 | 12869 |
| | | (*D. mon* vs *D. fla*) | IM *D. mon --> D. fla* | 705262 | 382053 | 1402532 | 2538521 | 1.09E-08 | -45638336 | 0 |
| | | | IM *D. fla --> D. mon* | 691899 | 395700 | 1457419 | 2398028 | 1.21E-09 | -45650952 | 12616 |
| | | Sympatric comparison | DIV | 719795 | 392311 | 1458664 | 2342807 | - | -136875659 | 47790 |
| | | (*D. mon* vs *D. fla*) | IM *D. mon --> D. fla* | 735267 | 376923 | 1387707 | 2526063 | 1.29E-08 | -136827869 | 0 |
| | | | IM *D. fla --> D. mon* | 718542 | 393103 | 1447342 | 2375536 | 2.13E-09 | -136873722 | 45853 |
| | | | **Model** | ***D. mon allop* Ne** | ***D. mon symp* Ne** | **Ancestral Ne** | **T** | **M** | **lnCL** | **ΔlnCL** |
| | | Intraspecific comparison | DIV | 1086549 | 1560399 | 858472 | 209661 | - | -39887382 | 0 |
| | | (*D. mon* vs *D. mon*) | IM *D. mon symp. --> D. mon allop.* | 1086577 | 1560435 | 858470 | 209663 | 1.50E-15 | -39887382 | 0 |
| | | | IM *D. mon allop. --> D. mon symp.* | 1079043 | 1441403 | 857987 | 216639 | 3.32E-07 | -39887408 | 27 |
| | **Block size** | | **Model** | ***D. mon* Ne** | ***D. fla* Ne** | **Ancestral Ne** | **T** | **M** | **lnCL** | **ΔlnCL** |
| | 128 bp | Allopatric comparison | DIV | 728545 | 398774 | 1132250 | 2710977 | - | -26255962 | 15401 |
| | | (*D. mon* vs *D. fla*) | IM *D. mon --> D. fla* | 732787 | 388220 | 1034471 | 2881985 | 7.62E-09 | -26240561 | 0 |
| | | | IM *D. fla --> D. mon* | 725401 | 398056 | 1078665 | 2805472 | 3.40E-09 | -26249397 | 8836 |
| | | Sympatric comparison | DIV | 757078 | 395703 | 1134780 | 2671276 | - | -78733822 | 56290 |
| | | (*D. mon* vs *D. fla*) | IM *D. mon --> D. fla* | 762355 | 383306 | 1022312 | 2868134 | 9.05E-09 | -78677532 | 0 |
| | | | IM *D. fla --> D. mon* | 752964 | 394863 | 1070200 | 2787162 | 4.37E-09 | -78707531 | 29999 |
| | | | **Model** | ***D. mon allop* Ne** | ***D. mon symp* Ne** | **Ancestral Ne** | **T** | **M** | **lnCL** | **ΔlnCL** |
| | | Intraspecific comparison | DIV | 1638756 | 2453210 | 776166 | 281292 | - | -26289327 | 14070 |
| | | (*D. mon* vs *D. mon*) | IM *D. mon symp. --> D. mon allop.* | 417656 | 1269040 | 704689 | 751648 | 5.86E-06 | -26280840 | 5583 |
| | | | IM *D. mon allop. --> D. mon symp.* | 1041239 | 642010 | 696082 | 917483 | 4.42E-06 | -26275257 | 0 |
| **Reference genome used** | **Block size** | **Comparison** | **Model** | ***D. mon* Ne** | ***D. fla* Ne** | **Ancestral Ne** | **T** | **M** | **lnCL** | **ΔlnCL** |
| *D. flavomontana* | 64 bp | Allopatric comparison | DIV | 704135 | 356493 | 1462359 | 2399150 | - | -46064633 | 9057 |
| | | (*D. mon* vs *D. fla*) | IM *D. mon --> D. fla* | 713873 | 346500 | 1415033 | 2518428 | 7.81E-09 | -46055576 | 0 |
| | | | IM *D. fla --> D. mon* | 703666 | 356694 | 1455996 | 2416828 | 1.01E-09 | -46064391 | 8816 |
| | | Sympatric comparison | DIV | 753925 | 352945 | 1439661 | 2344076 | - | -138110861 | 35847 |
| | | (*D. mon* vs *D. fla*) | IM *D. mon --> D. fla* | 767091 | 340667 | 1381988 | 2487543 | 9.83E-09 | -138075014 | 0 |
| | | | IM *D. fla --> D. mon* | 752758 | 353451 | 1425934 | 2383428 | 2.43E-09 | -138107834 | 32820 |
| | | | **Model** | ***D. mon allop* Ne** | ***D. mon symp* Ne** | **Ancestral Ne** | **T** | **M** | **lnCL** | **ΔlnCL** |
| | | Intraspecific comparison | DIV | 1092818 | 2805737 | 826433 | 173390 | - | -37761907 | 54 |
| | | (*D. mon* vs *D. mon*) | IM *D. mon symp. --> D. mon allop.* | 1092873 | 2806425 | 826437 | 173381 | 1.27E-11 | -37761907 | 54 |
| | | | IM *D. mon allop. --> D. mon symp.* | 1079681 | 2499743 | 825799 | 180657 | 5.24E-07 | -37761853 | 0 |
| | **Block size** | | **Model** | ***D. mon* Ne** | ***D. fla* Ne** | **Ancestral Ne** | **T** | **M** | **lnCL** | **ΔlnCL** |
| | 128 bp | Allopatric comparison | DIV | 745937 | 359180 | 1130565 | 2725366 | - | -26607436 | 11739 |
| | | (*D. mon* vs *D. fla*) | IM *D. mon --> D. fla* | 749783 | 350820 | 1050428 | 2862796 | 5.92E-09 | -26595697 | 0 |
| | | | IM *D. fla --> D. mon* | 744008 | 358222 | 1086957 | 2800850 | 2.49E-09 | -26602466 | 6769 |
| | | Sympatric comparison | DIV | 799335 | 355370 | 1118044 | 2674056 | - | -79835822 | 44004 |
| | | (*D. mon* vs *D. fla*) | IM *D. mon --> D. fla* | 804432 | 345201 | 1022931 | 2836521 | 7.25E-09 | -79791818 | 0 |
| | | | IM *D. fla --> D. mon* | 796338 | 354096 | 1060491 | 2776345 | 3.62E-09 | -79813016 | 21198 |
| | | | **Model** | ***D. mon allop* Ne** | ***D. mon symp* Ne** | **Ancestral Ne** | **T** | **M** | **lnCL** | **ΔlnCL** |
| | | Intraspecific comparison | DIV | 1910759 | 5565491 | 742244 | 253322 | - | -25123546 | 17638 |
| | | (*D. mon* vs *D. mon*) | IM *D. mon symp. --> D. mon allop.* | 253437 | 1469856 | 661231 | 717155 | 1.57E-05 | -25111917 | 6009 |
| | | | IM *D. mon allop. --> D. mon symp.* | 1051701 | 887758 | 654390 | 898887 | 5.80E-06 | -25105908 | 0 |

Table S13. Genomes of the initial PacBio assemblies. N50 = half of the genome is in contigs larger than or equal the N50 contig size. N50 count = half of the genome consists of the number indicated in N50 count.

| Assembler | | *D. montana* Seward, Alaska monSE13F37 | *D. montana* Jackson, USA monJX13F48 | *D. flavomontana* Vancouver, Canada flaVAN14F20 | *D. flavomontana* Livingston, USA flaMT13F11 |
|---|---|---|---|---|---|
| **wtdbg2 (RedBean)** | **Genome size (Mb)** | 199.9 | 181.3 | 212.5 | 207.0 |
| | **Total contigs** | 1002 | 1168 | 1780 | 1124 |
| | **Longest contig (Mb)** | 20.3 | 14.5 | 19.8 | 28.8 |
| | **N50 (Mb)** | 2.5 | 0.9 | 2.7 | 2.6 |
| | **N50 count** | 15 | 41 | 12 | 12 |
| | **GC%** | 0.417 | 0.404 | 0.410 | 0.406 |
| **MaSuRCA** | **Genome size (Mb)** | 199.1 | 185.6 | 202.5 | 200.8 |
| | **Total contigs** | 873 | 1251 | 733 | 845 |
| | **Longest contig (Mb)** | 16.5 | 5.5 | 16.1 | 15.6 |
| | **N50 (Mb)** | 1.7 | 0.5 | 2.2 | 2.1 |
| | **N50 count** | 25 | 86 | 15 | 26 |
| | **GC%** | 0.408 | 0.403 | 0.400 | 0.401 |

Table S14. Genome assemblies of single wild-caught females (Illumina 150b paired-end data) used in dN/dS analysis. N50 = half of the genome is in contigs larger than or equal the N50 contig size. N50 count = half of the genome consists of the number indicated in N50 count.

| Species | Population | | Fly strain | Genome size (Mb) | Total contigs | Longest contig (bp) | N50 (bp) | N50 count | GC% | Total complete BUSCOs (%, n: 3285) | Single copy BUSCOs (%) | Duplicated BUSCOs (%) | Sample used in dN/dS analysis |
|---|---|---|---|---|---|---|---|---|---|---|---|---|---|
| *D. littoralis* | Outgroup (Finland) | Korpilahti | KL13F60 | 205.1 | 23756 | 642147 | 67694 | 751 | 40.5 | 98.4 | 97.7 | 0.7 | yes |
| *D. montana* | Sympatry (western coast) | Terrace | monTER14F11 | 218.3 | 32634 | 680450 | 41496 | 1386 | 41.0 | 98.0 | 97.2 | 0.8 | yes |
| | | Vancouver | monVAN14F1 | 205.8 | 50249 | 127841 | 11406 | 4956 | 40.5 | 92.2 | 89.6 | 2.6 | yes |
| | | Ashford | monASH13F13 | 225.3 | 119802 | 52723 | 3986 | 15697 | 40.3 | 79.9 | 78.8 | 1.1 | no |
| | | Fall Creek | mon4Fall | 206.6 | 28369 | 423492 | 45341 | 1191 | 40.4 | 97.8 | 97.1 | 0.7 | yes |
| | | Azalea | monAZA2 | 210.1 | 32250 | 428221 | 36497 | 1498 | 40.5 | 97.9 | 97.4 | 0.5 | yes |
| | Sympatry (Rocky Mountains) | McBride | monMB14F1 | 216.9 | 59047 | 372247 | 17321 | 3115 | 41.0 | 95.6 | 93.8 | 1.8 | yes |
| | | Cranbrook | monCRAN14F16 | 228.2 | 47760 | 728611 | 37548 | 1476 | 41.4 | 97.9 | 97.0 | 0.9 | yes |
| | | Jackson | monJX13F3 | 224.4 | 57826 | 181731 | 11395 | 5290 | 40.5 | 92.4 | 89.2 | 3.2 | yes |
| | | Afton | monAF15F28 | 227.5 | 73398 | 316902 | 9702 | 6161 | 40.6 | 91.7 | 90.0 | 1.7 | yes |
| *D. flavomontana* | Sympatry (western coast) | Terrace | flaTER14F5 | 209.4 | 28746 | 745447 | 58956 | 906 | 40.2 | 98.2 | 97.6 | 0.6 | yes |
| | | Vancouver | flaVAN14F20 | 203.4 | 49591 | 177737 | 13290 | 4092 | 40.4 | 93.2 | 91.8 | 1.4 | yes |
| | | Ashford | flaASH13F2 | 202.2 | 40399 | 165684 | 13556 | 4091 | 40.7 | 95.4 | 94.2 | 1.2 | yes |
| | | Fall Creek | fla3Fall | 203.1 | 29004 | 550168 | 50722 | 1011 | 40.3 | 97.7 | 97.0 | 0.7 | yes |
| | Sympatry (Rocky Mountains) | McBride | flaMB14F10 | 227.1 | 41561 | 588908 | 52916 | 1018 | 41.8 | 98.0 | 97.3 | 0.7 | yes |
| | | Cranbrook | flaCRAN14F7 | 212.8 | 32100 | 613853 | 50615 | 1062 | 40.5 | 98.0 | 97.5 | 0.5 | yes |
| | | Livingston | flaMT13F11 | 204.6 | 65403 | 100004 | 7933 | 7202 | 40.6 | 83.2 | 80.5 | 2.7 | no |
| | | Jackson | flaJX13F38 | 212.8 | 49728 | 213613 | 13230 | 4408 | 40.6 | 90.9 | 88.3 | 2.6 | yes |
| | | Liberty | flaLB15F3 | 205.2 | 52507 | 159220 | 10707 | 5293 | 40.3 | 83.8 | 80.6 | 3.2 | no |

### Supplementary figures

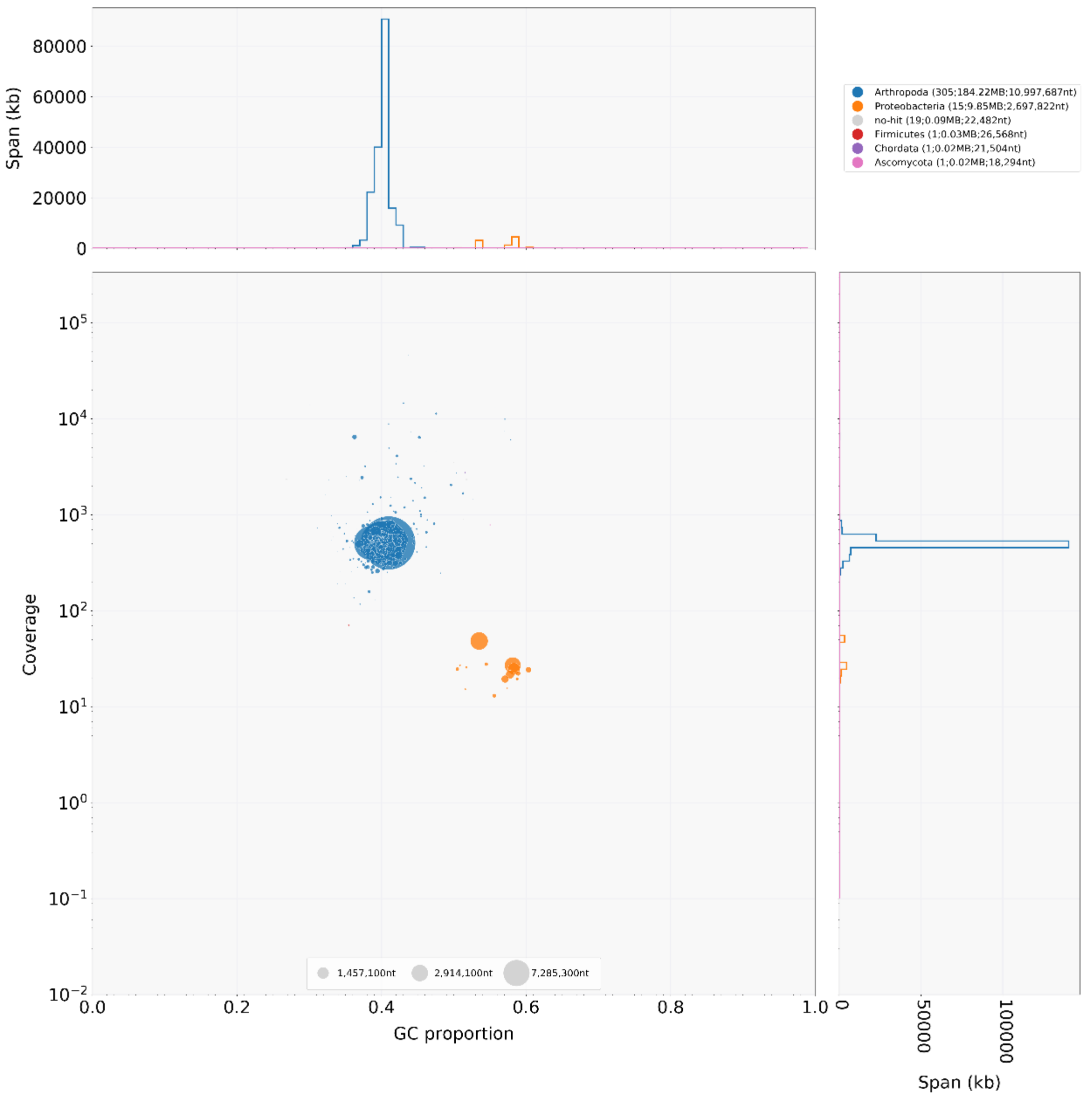

Figure S1. Genomic contaminants of monSE13F37 (*D. montana*) assembly illustrated with blobplot (Laetsch and Blaxter 2017).

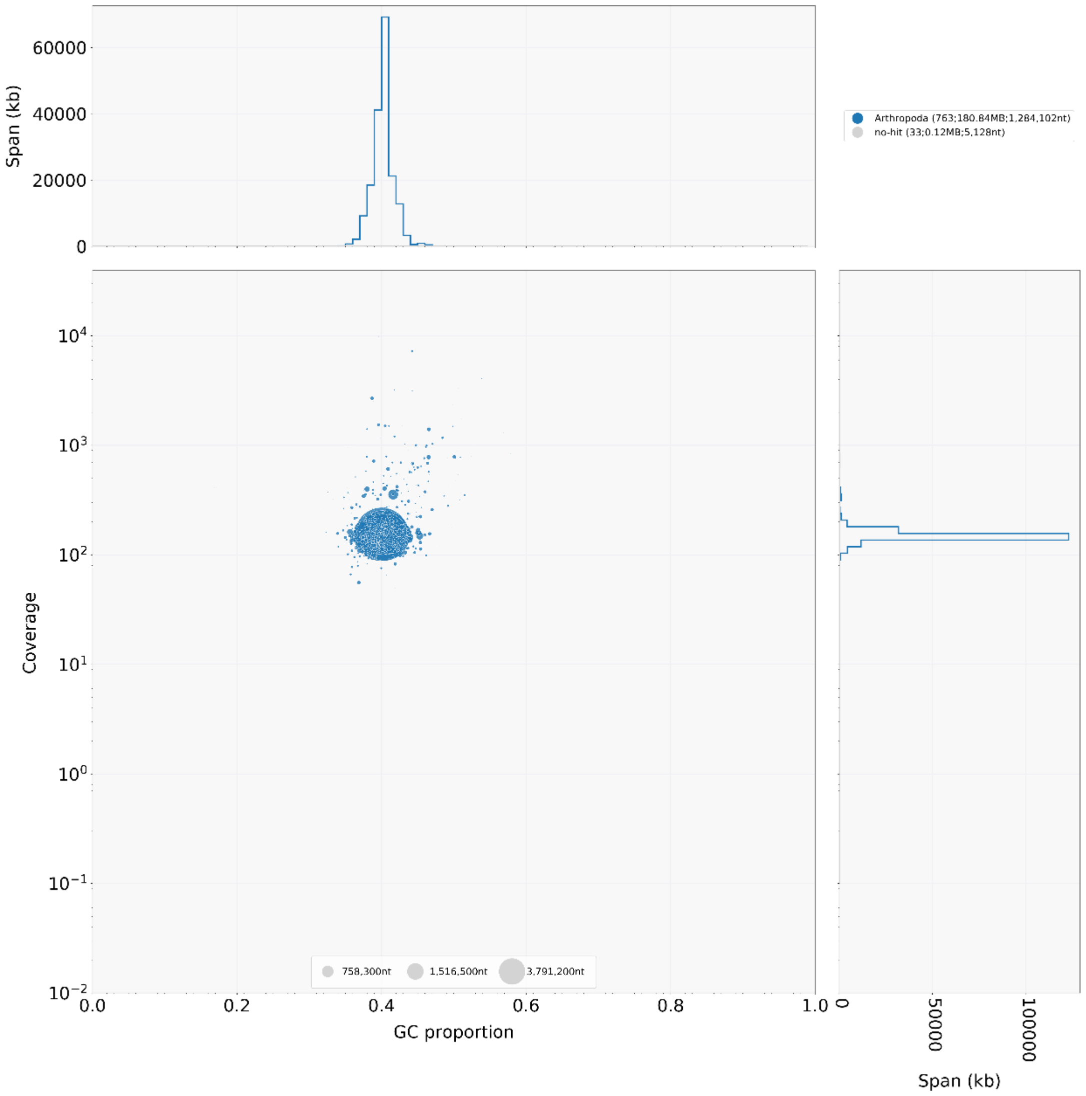

Figure S2. Genomic contaminants of monJX13F48 (*D. montana*) assembly illustrated with blobplot (Laetsch and Blaxter 2017).

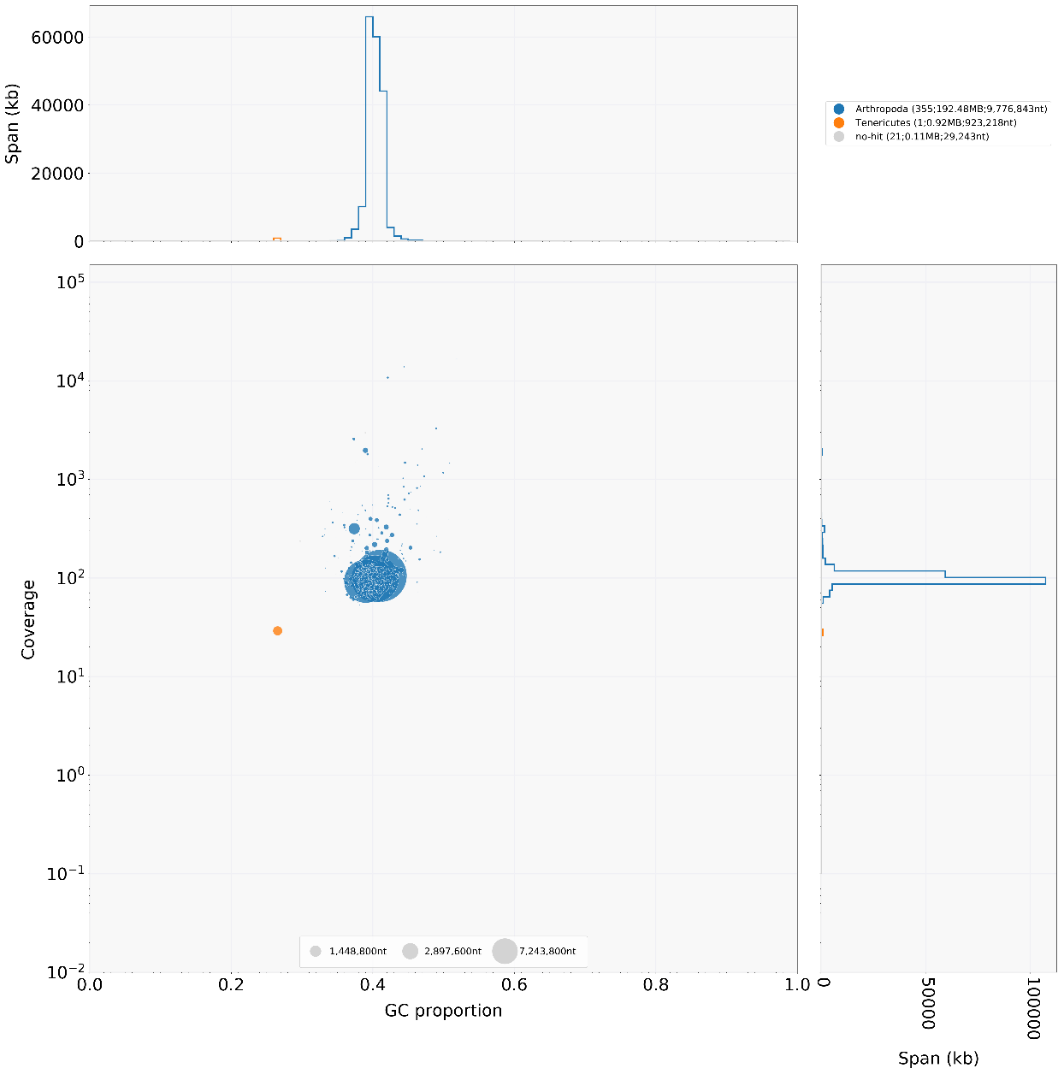

Figure S3. Genomic contaminants of flaMT13F11 (*D. flavomontana*) assembly illustrated with blobplot (Laetsch and Blaxter 2017).

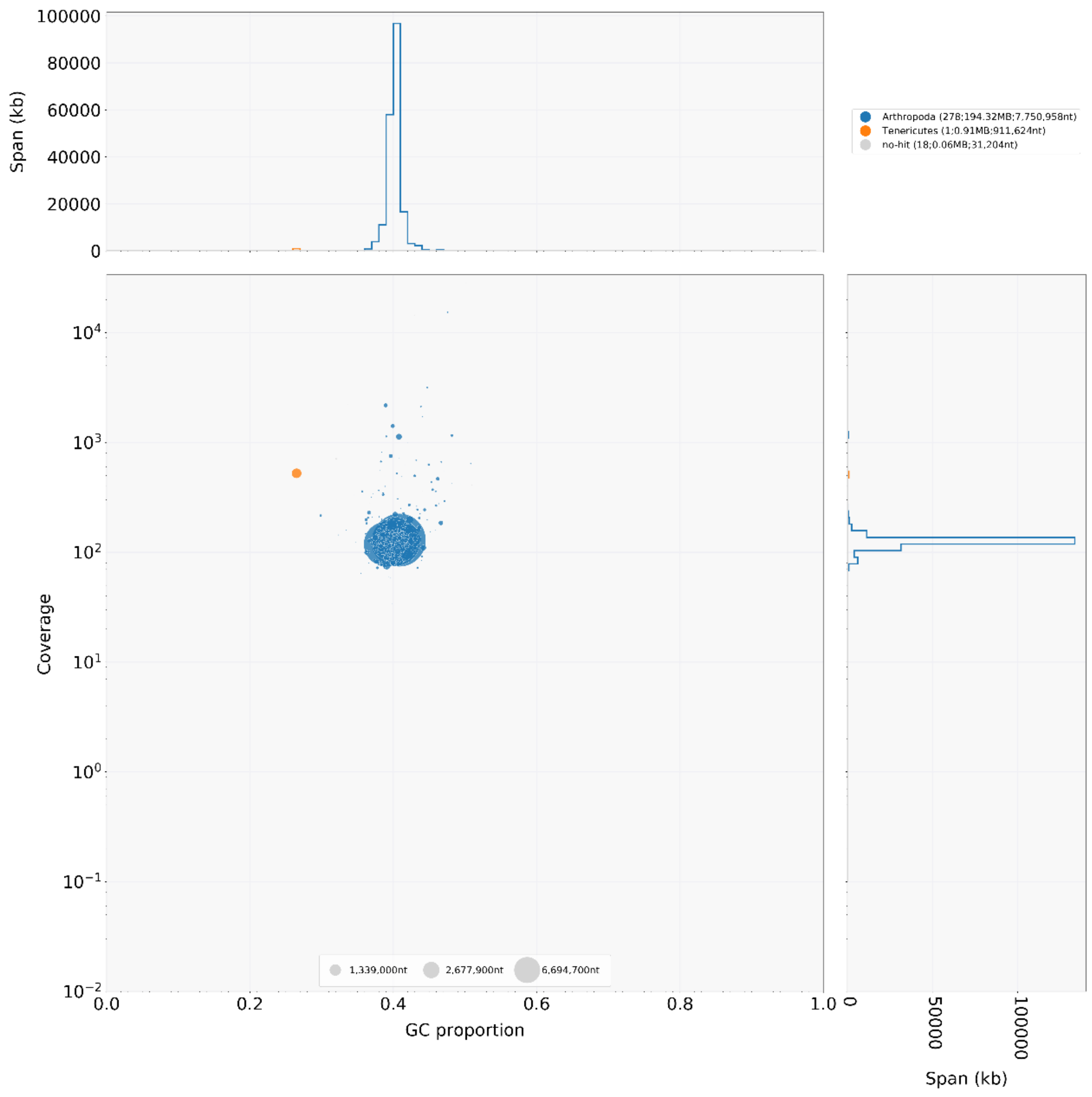

Figure S4. Genomic contaminants of flaVAN14F20 (*D. flavomontana*) assembly illustrated with blobplot (Laetsch and Blaxter 2017).

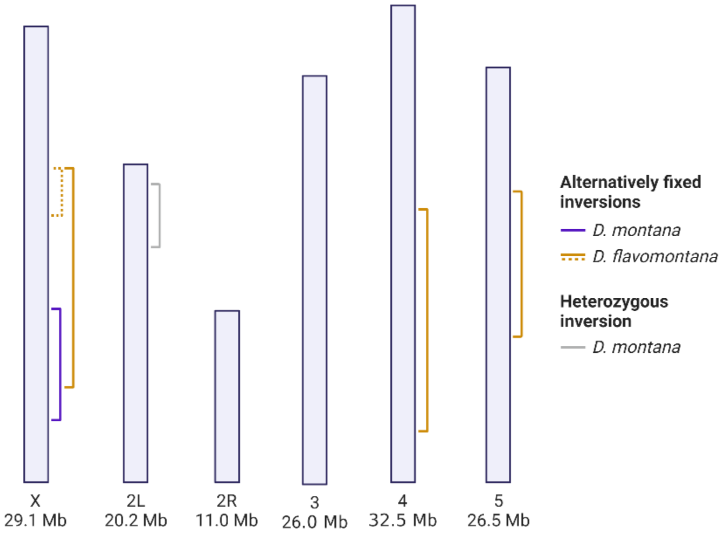

Figure S5. Chromosomes and five large (>0.5Mb) alternatively fixed inversions of *D. montana* (blue solid line) and *D. flavomontana* (orange solid and dashed lines), as well as one putative heterozygous inversion of *D. montana* (light grey solid line) (Table S9). The numbers below each chromosome refer to its length in the *D. montana* assembly. Chromosome 2 has left (2L) and right (2R) arms that are separated by a submetacentric centromere.

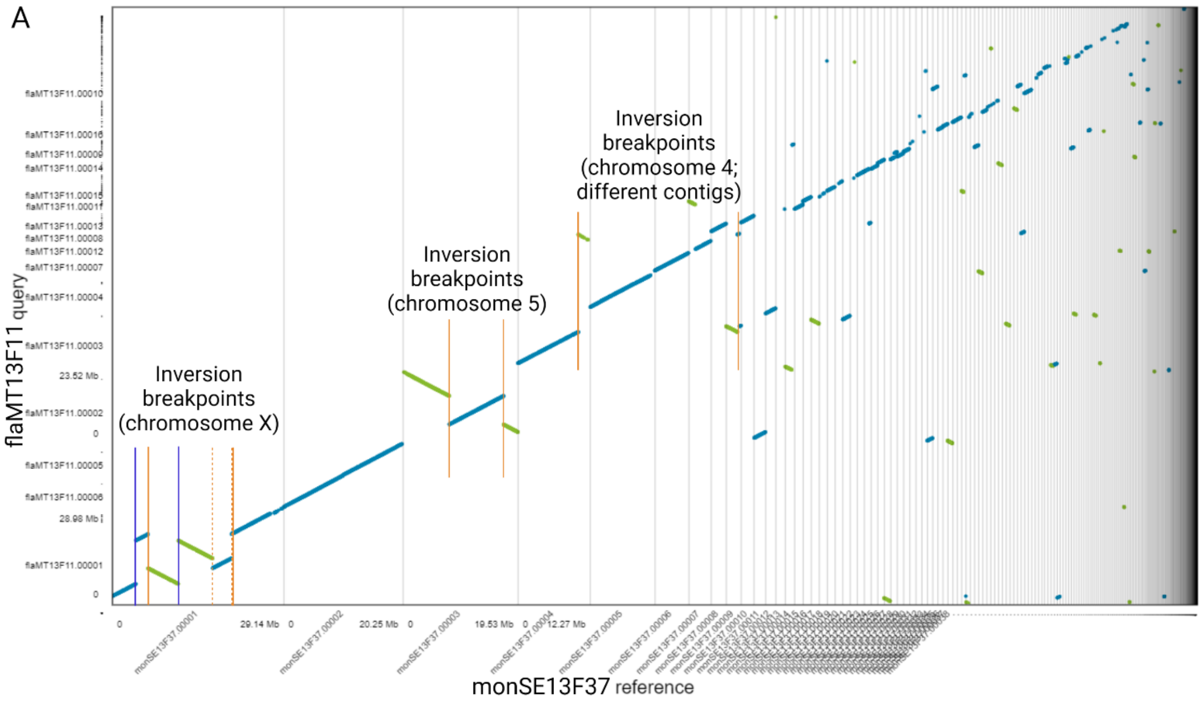

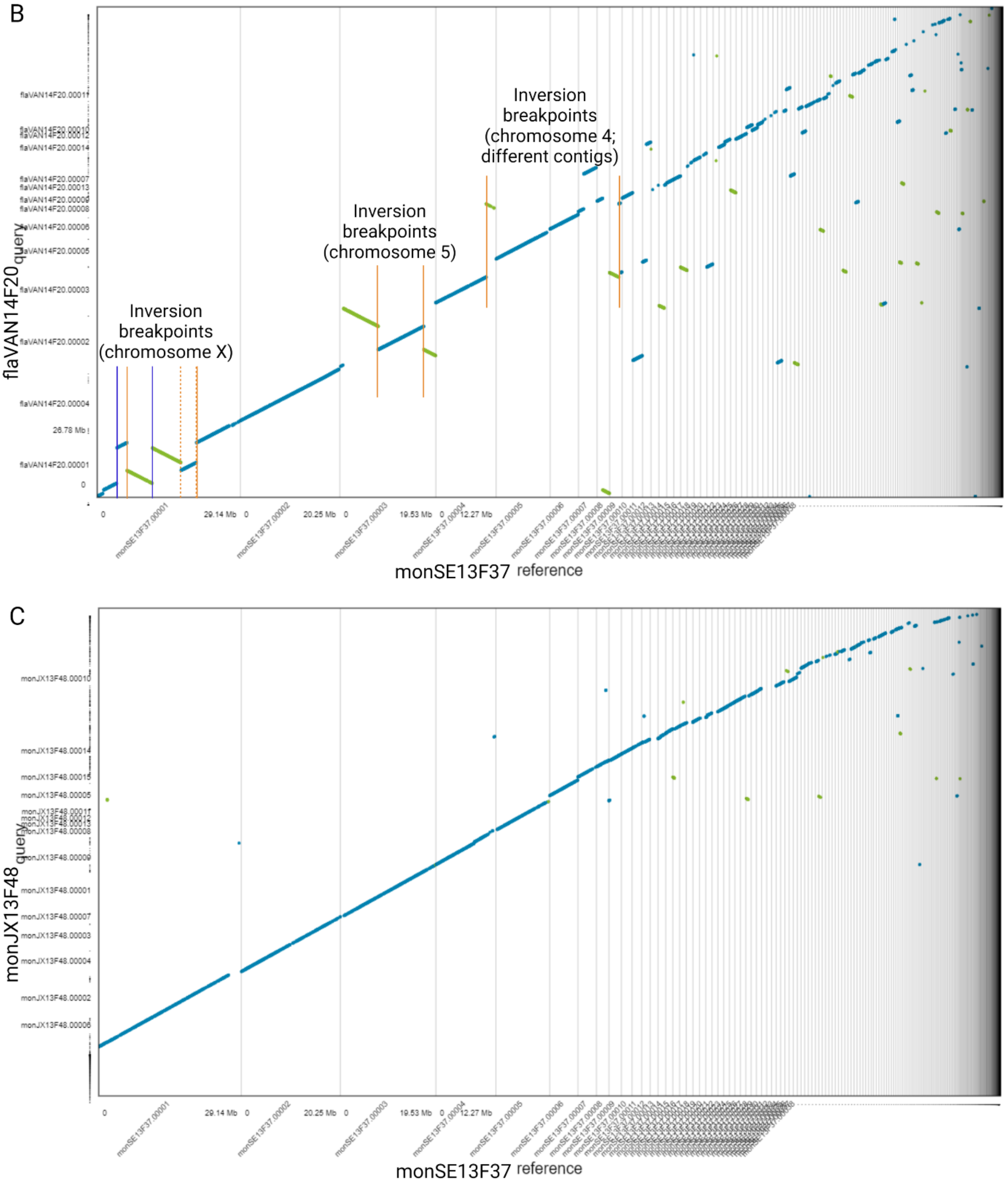

Figure S6. Nucmer alignments of the MUMmer package (Marçais et al. 2018) illustrated with Dot plots (https://dot.sandbox.bio/). *D. montana* monSE13F37 was used as a reference and (A) *D. flavomontana* flaMT13F11, (B) *D. flavomontana* flaVAN14F20, or (C) *D. montana* monJX13F48 as query. A change in color and orientation within a single pair of contig (e.g. monSE13F37.00001 and flaMT13F11.00001) indicate inverted regions. Inversions are evident only when both breakpoints are found within the same contig in both the reference and the query. Alternatively fixed inversions of *D. montana* and *D. flavomontana* are marked with blue solid lines, and orange solid and dashed lines respectively (see Table S9, Supplementary file 1).

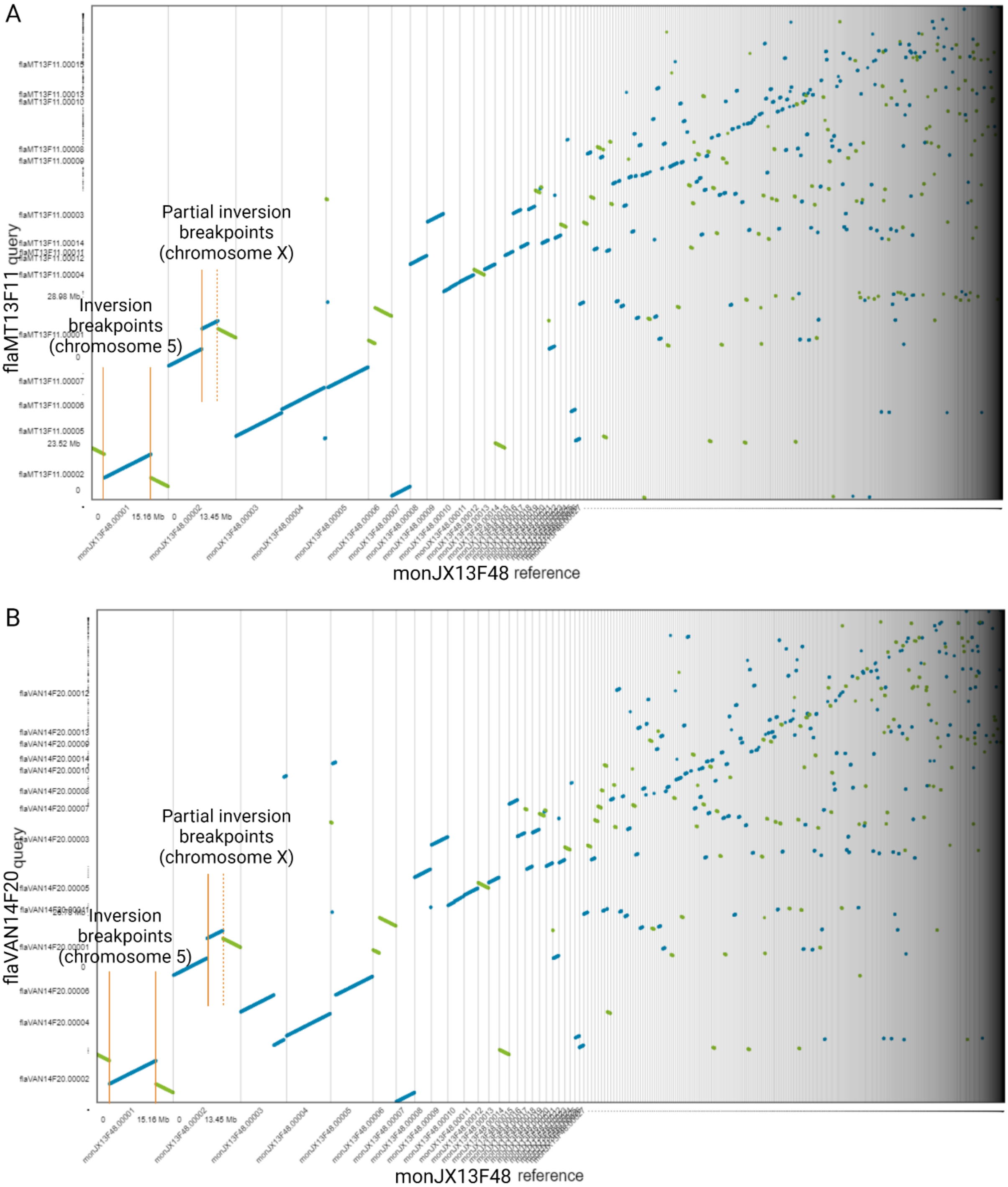
A
Partial inversion breakpoints (chromosome X)
Inversion breakpoints (chromosome 5)
flaMT13F11 query
monJX13F48 reference
B
Partial inversion breakpoints (chromosome X)
Inversion breakpoints (chromosome 5)
flaVAN14F20 query
monJX13F48 reference

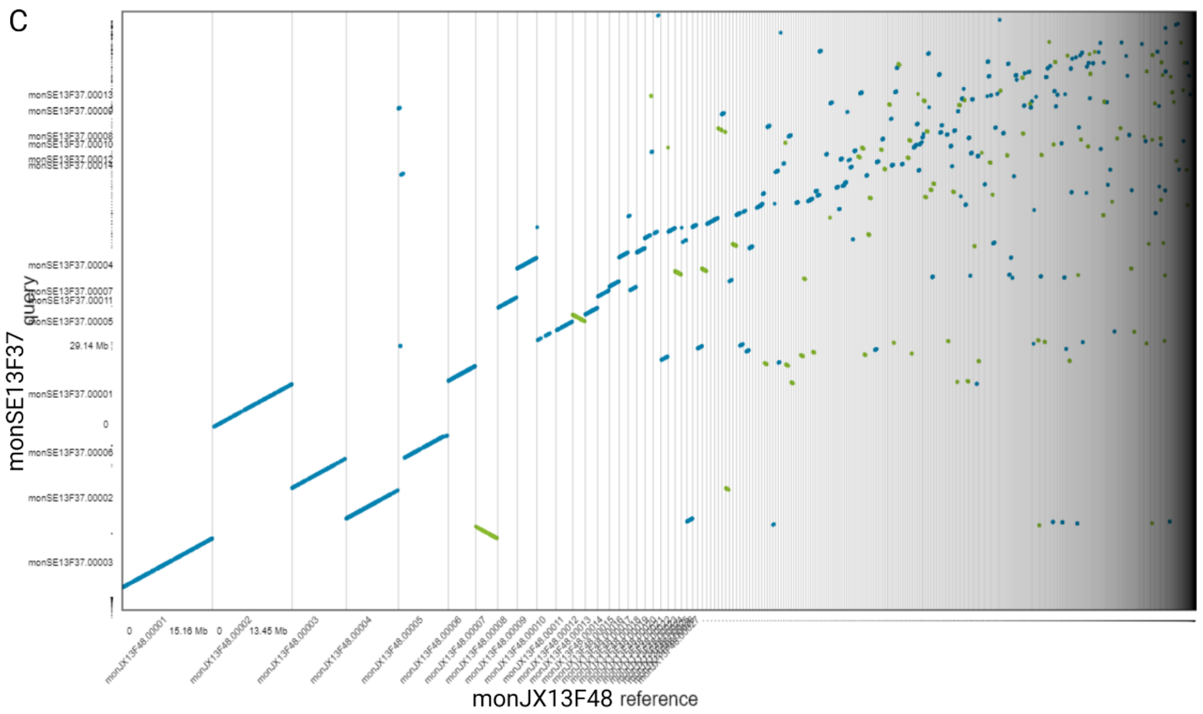

Figure S7. Nucmer alignments of the MUMmer package (Marçais et al. 2018) illustrated with Dot plots (https://dot.sandbox.bio/). *D. montana* monJX13F48 was used as a reference and (A) *D. flavomontana* flaMT13F11, (B) *D. flavomontana* flaVAN14F20, or (C) *D. montana* monSE13F37 as query. For guidance on the plot see Fig. S6.

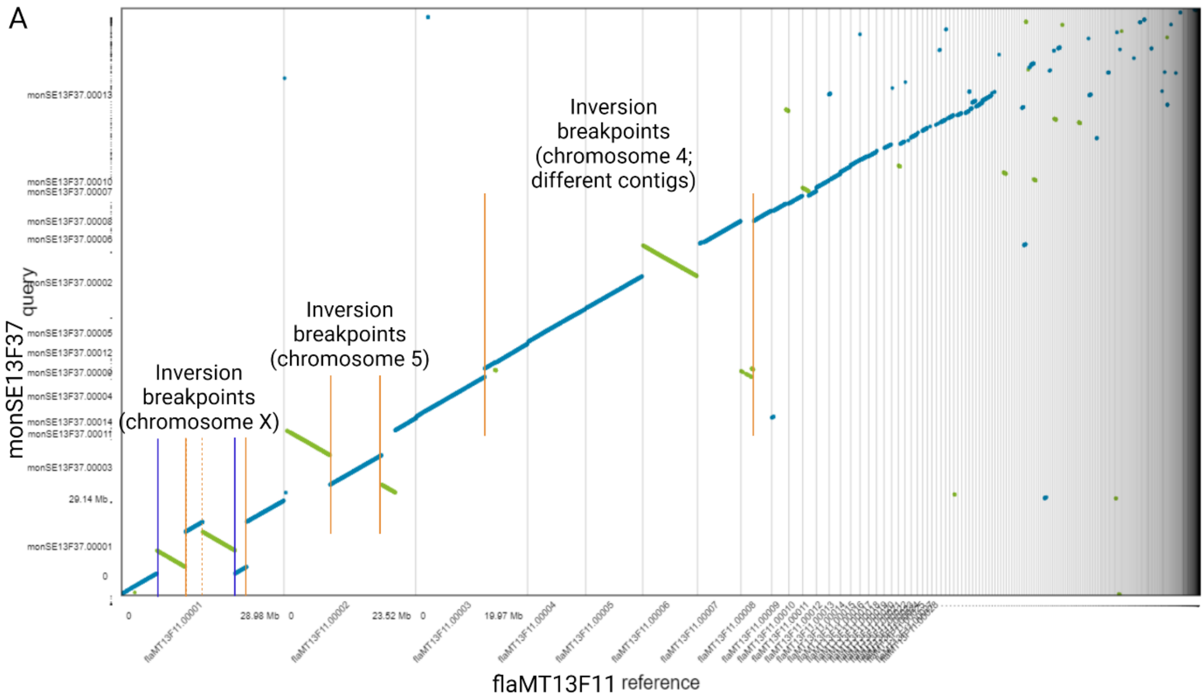

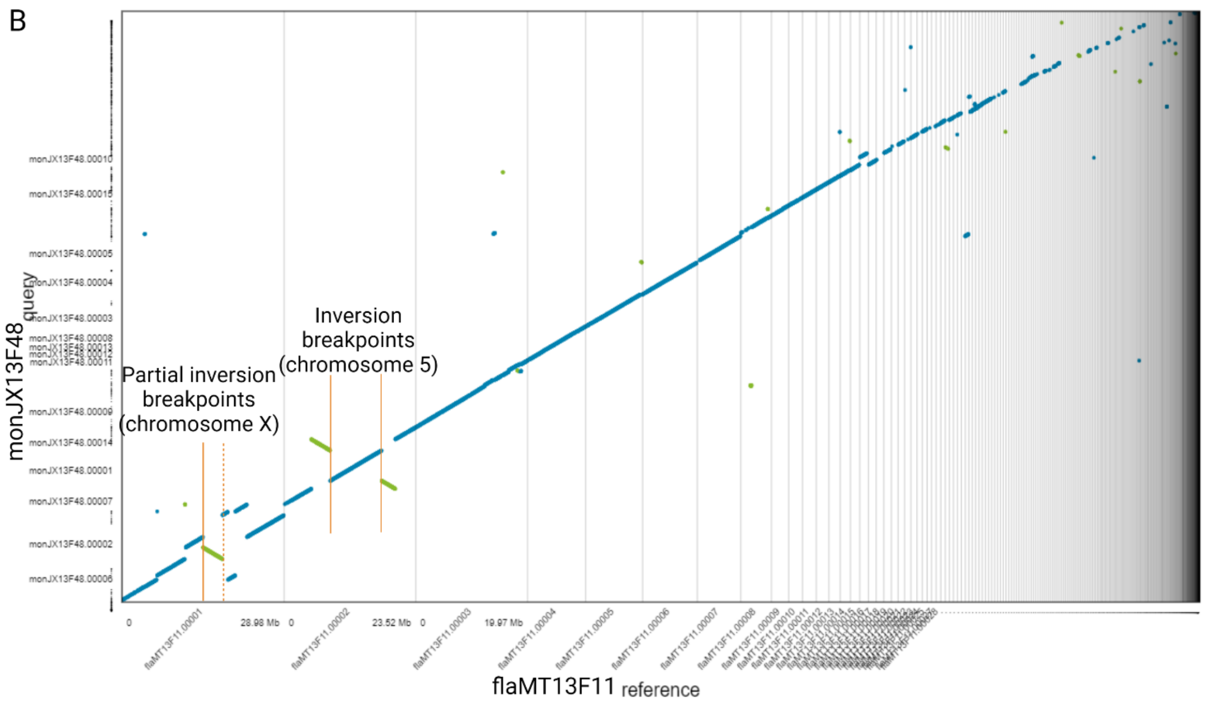

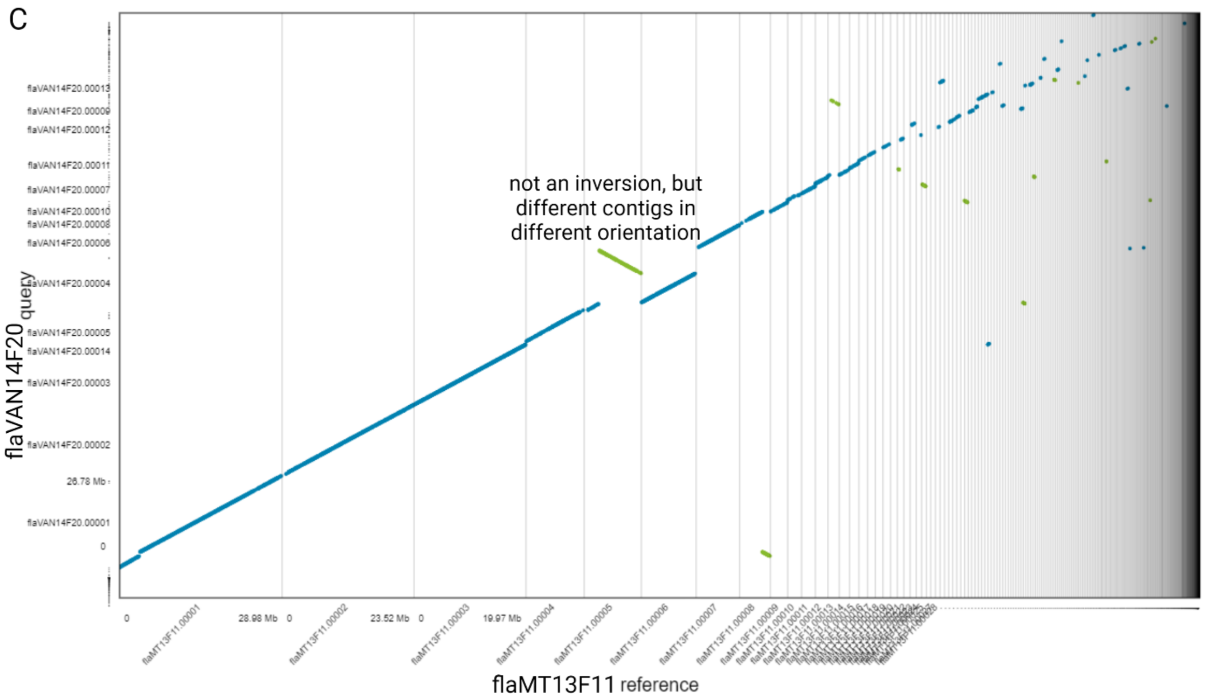

Figure S8. Nucmer alignments of the MUMmer package (Marçais et al. 2018) illustrated with Dot plots (https://dot.sandbox.bio/). *D. flavomontana* flaM13F11 was used as a reference and (A) *D. montana* monSE13F37, (B) *D. montana* monJX13F48, and (C) *D. flavomontana* flaVAN14F20 as query. For guidance on the plot see Fig. S6.

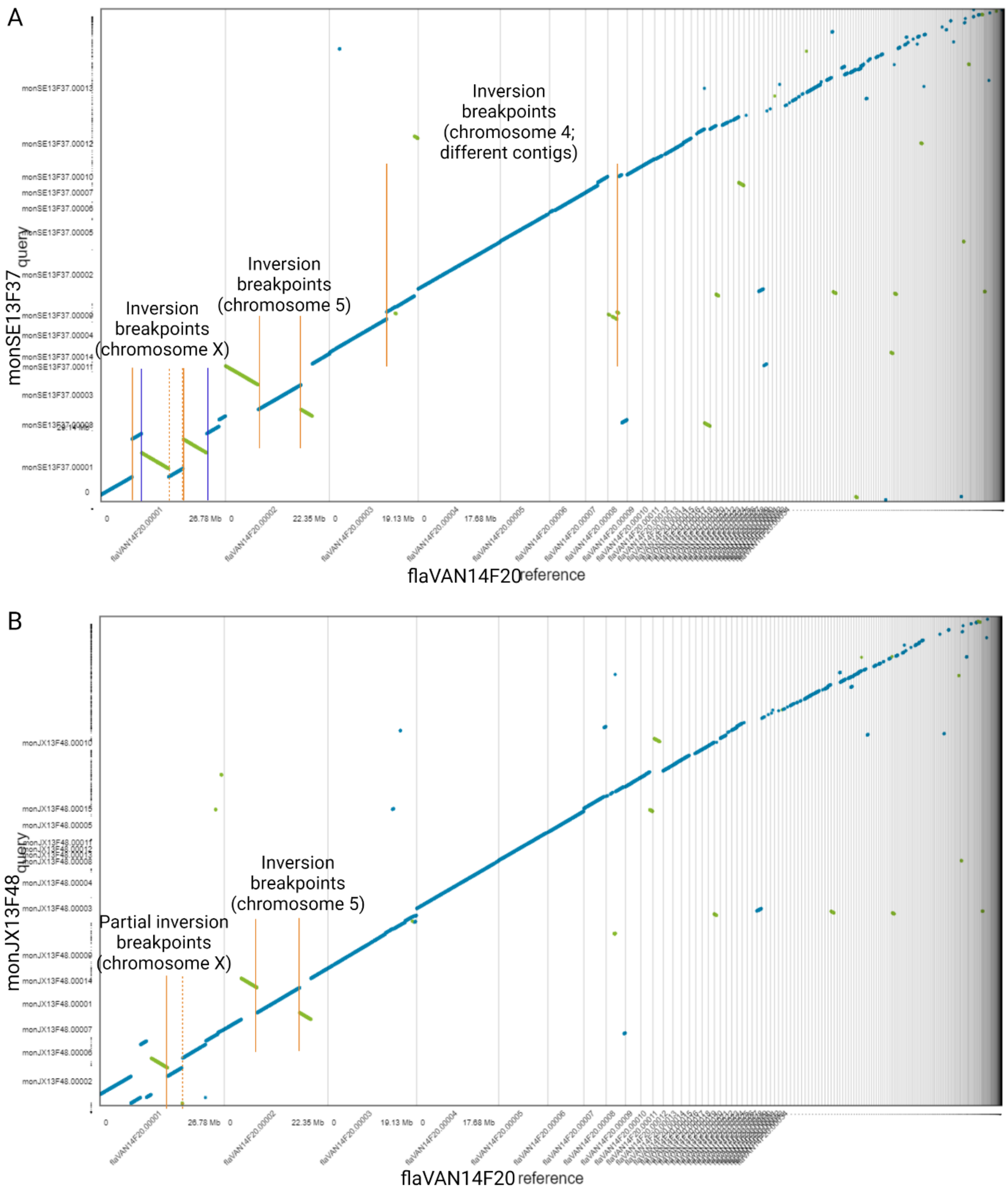
A
Inversion breakpoints (chromosome 4; different contigs)
Inversion breakpoints (chromosome 5)
Inversion breakpoints (chromosome X)
monSE13F37 query
flaVAN14F20 reference
B
Inversion breakpoints (chromosome 5)
Partial inversion breakpoints (chromosome X)
monJX13F48 query
flaVAN14F20 reference

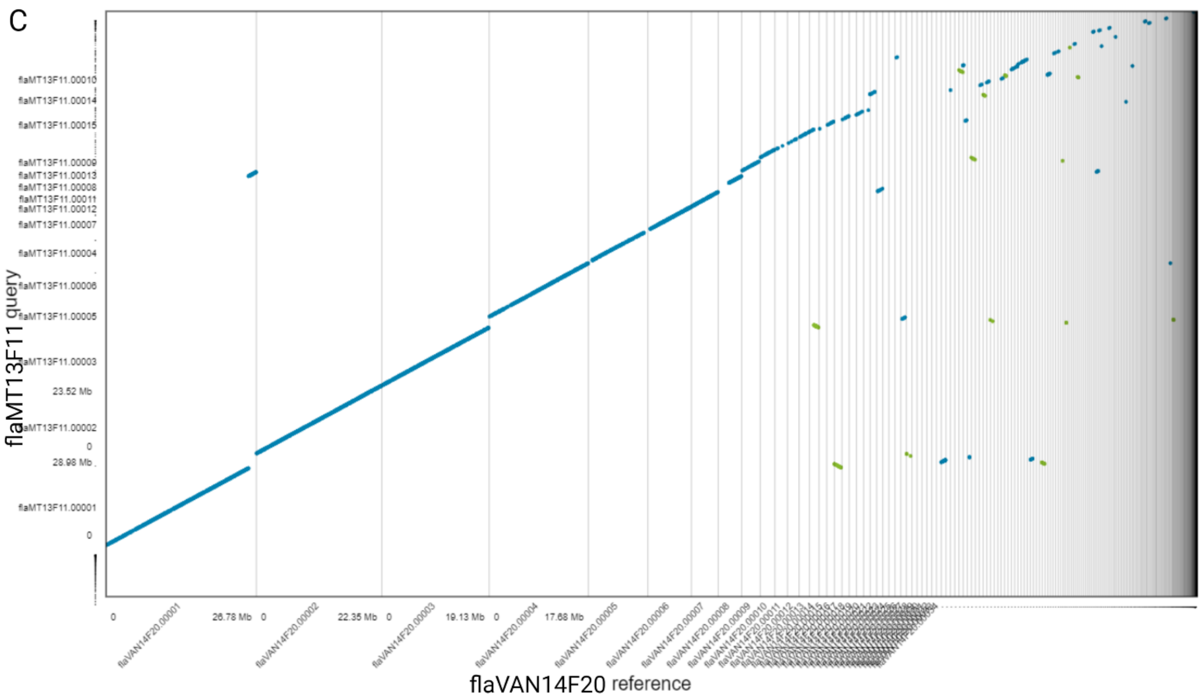

Figure S9. Nucmer alignments of the MUMmer package (Marçais et al. 2018) illustrated with Dot plots (https://dot.sandbox.bio/). *D. flavomontana* flaVAN14F20 was used as a reference and (A) *D. montana* monSE13F37, (B) *D. montana* monJX13F48, or (C) *D. flavomontana* flaMT13F11 as query. For guidance on the plot see Fig. S6.

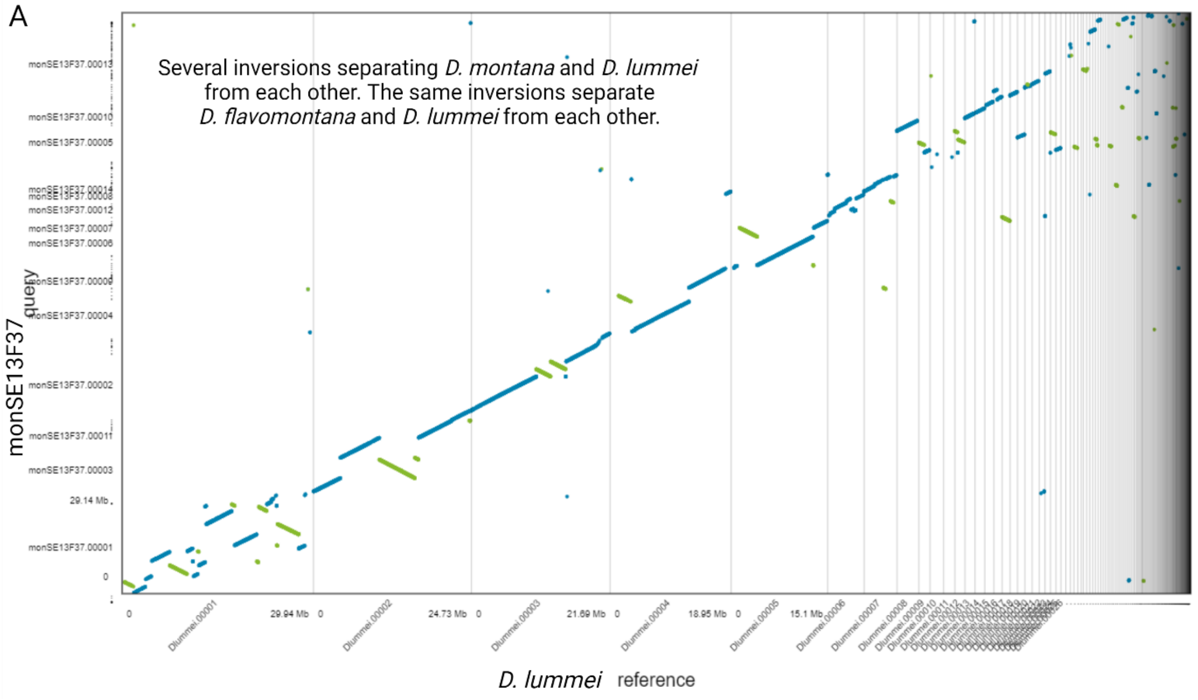

B

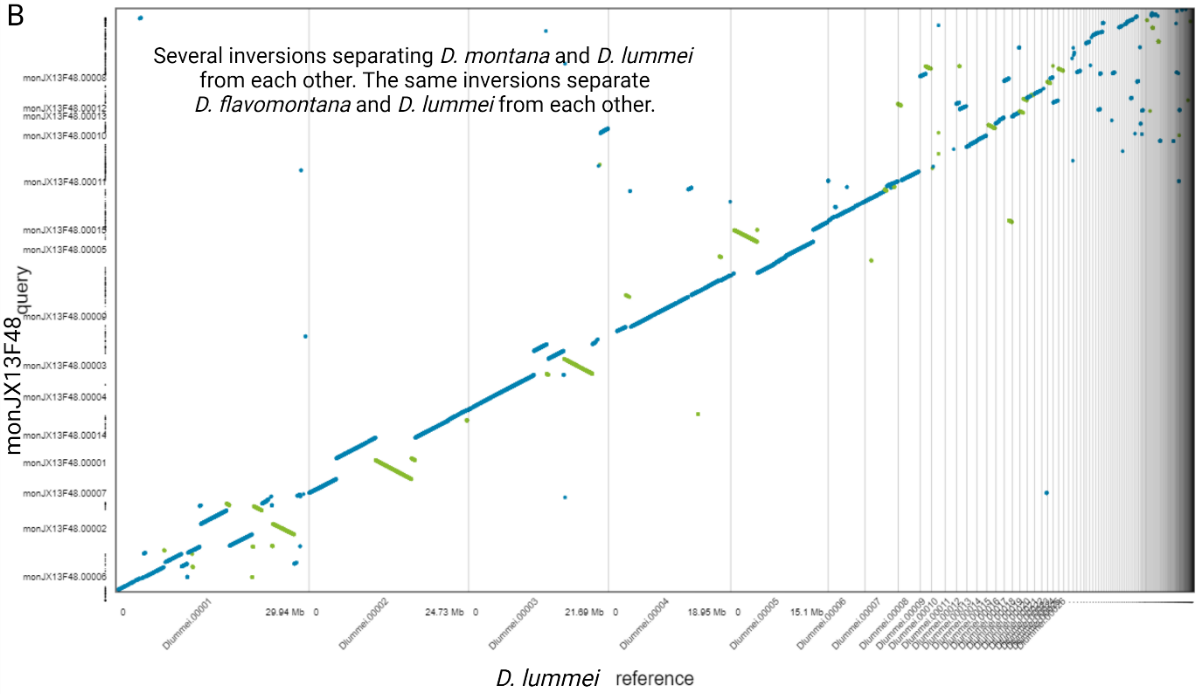
Several inversions separating D. montana and D. lummei
from each other. The same inversions separate
D. flavomontana and D. lummei from each other.
monJX13F48 query
29.94 Mb
24.73 Mb
21.69 Mb
18.95 Mb
15.1 Mb
D. lummei reference

C

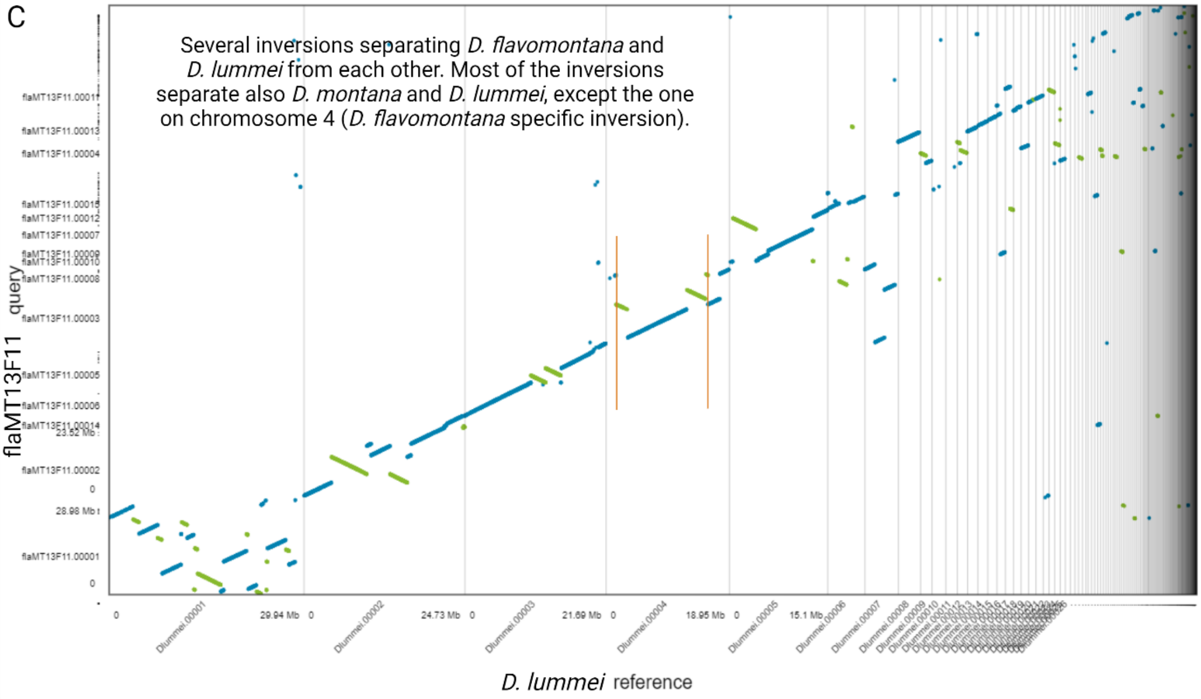
Several inversions separating D. flavomontana and
D. lummei from each other. Most of the inversions
separate also D. montana and D. lummei, except the one
on chromosome 4 (D. flavomontana specific inversion).
flaMT13F11 query
29.94 Mb
24.73 Mb
21.69 Mb
18.95 Mb
15.1 Mb
D. lummei reference

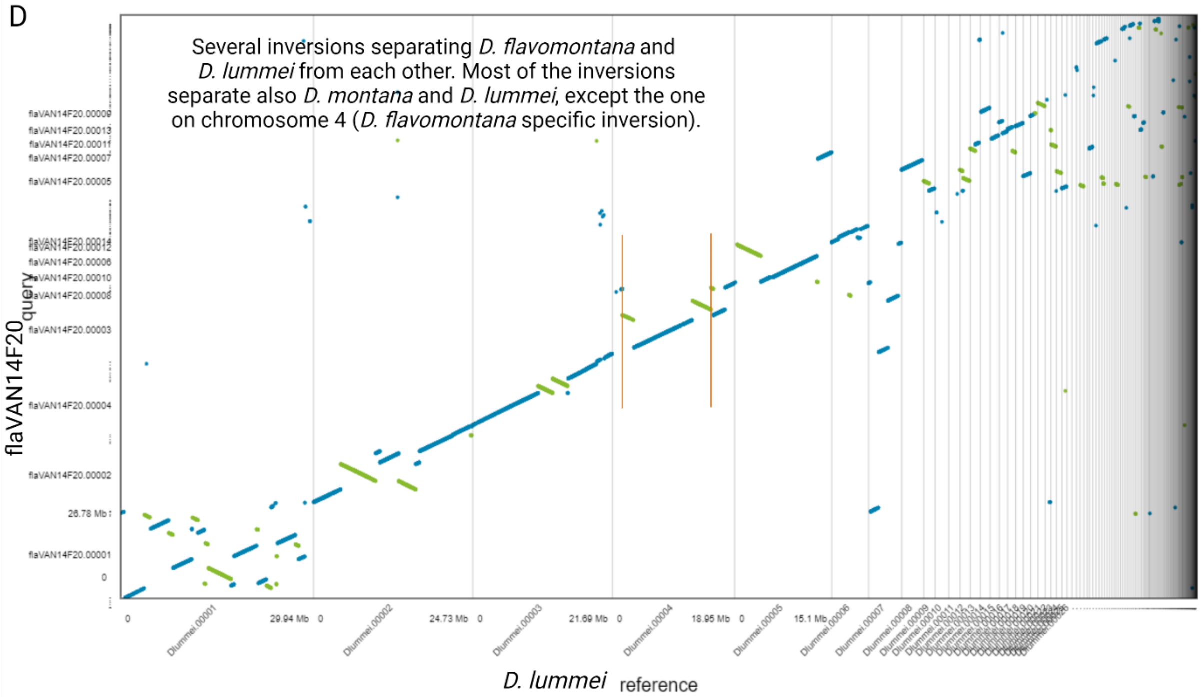

Figure S10. Nucmer alignments of the MUMmer package (Marçais et al. 2018) illustrated with Dot plots (https://dot.sandbox.bio/). *D. lummei* was used as a reference and (A) *D. montana* monSE13F37, (B) *D. montana* monJX13F48, or (C) *D. flavomontana* flaMT13F11, or (D) *D. flavomontana* flaVAN14F20 as query. For guidance on the plot see Fig. S6.

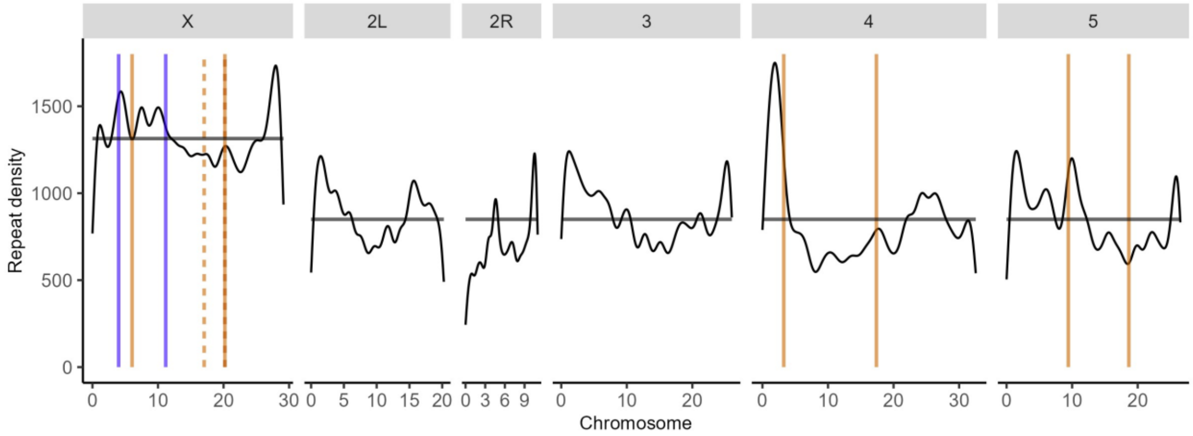

Figure S11. Density of repetitive sequences across the genome. Breakpoints of alternatively fixed inversions of *D. montana* and *D. flavomontana* are marked with solid blue, and solid and dashed orange lines, respectively (see Fig. S5). Black horizontal lines represent the mean repeat density for the X chromosome and autosomes.

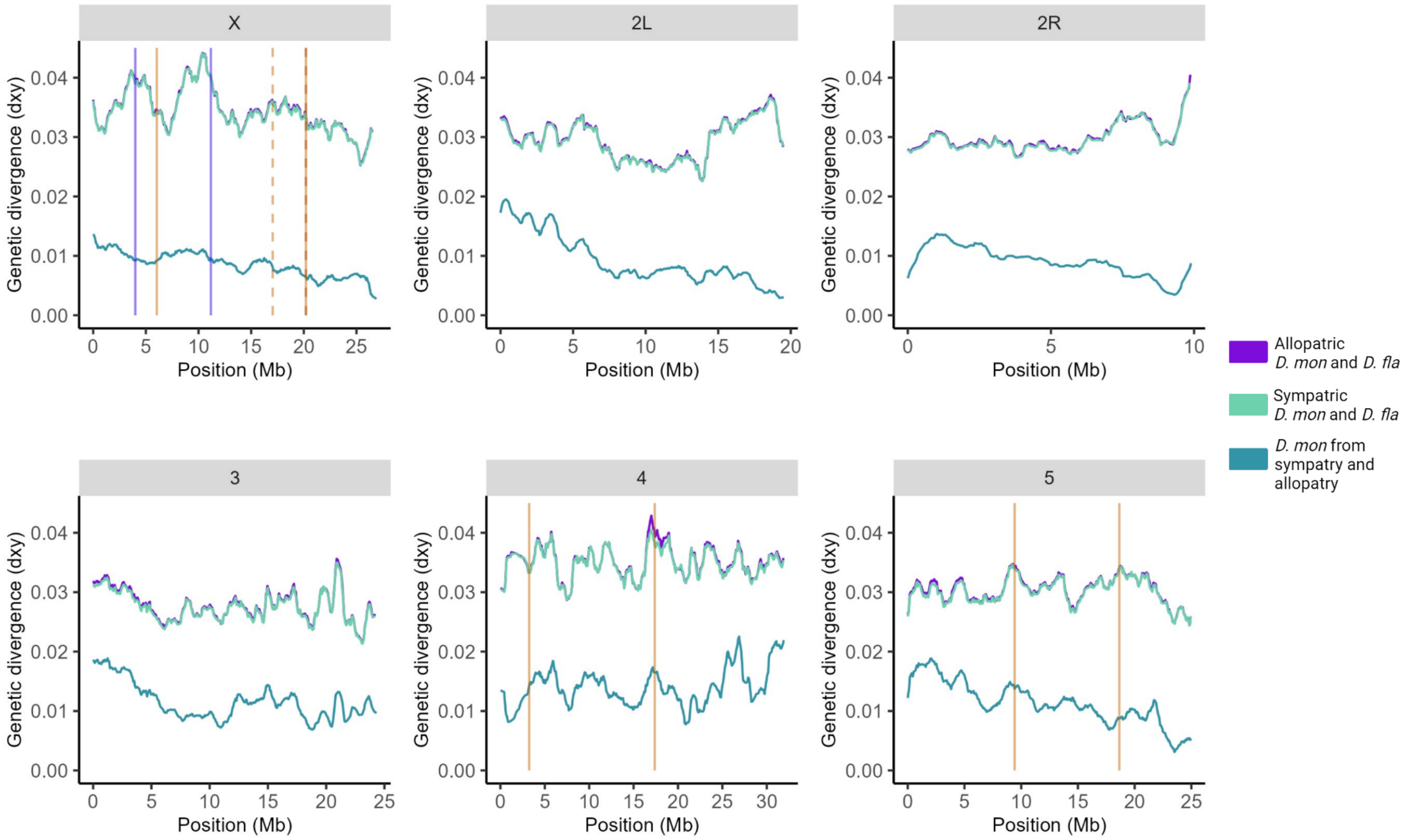

Figure S12. Genetic divergence (measured as $d_{XY}$) for different chromosomes (including intergenic regions) in sliding windows (window size 10 000 blocks, step size 1 000 blocks, block length 64b) for allopatric and sympatric comparisons of *D. montana* and *D. flavomontana* (interspecific), and for *D. montana* originating from allopatry and sympatry (intraspecific). Vertical lines represent breakpoints of different inversions (see Fig. S5): blue solid lines, and orange solid and dashed lines indicate alternatively fixed inversions of *D. montana* and *D. flavomontana*, respectively.

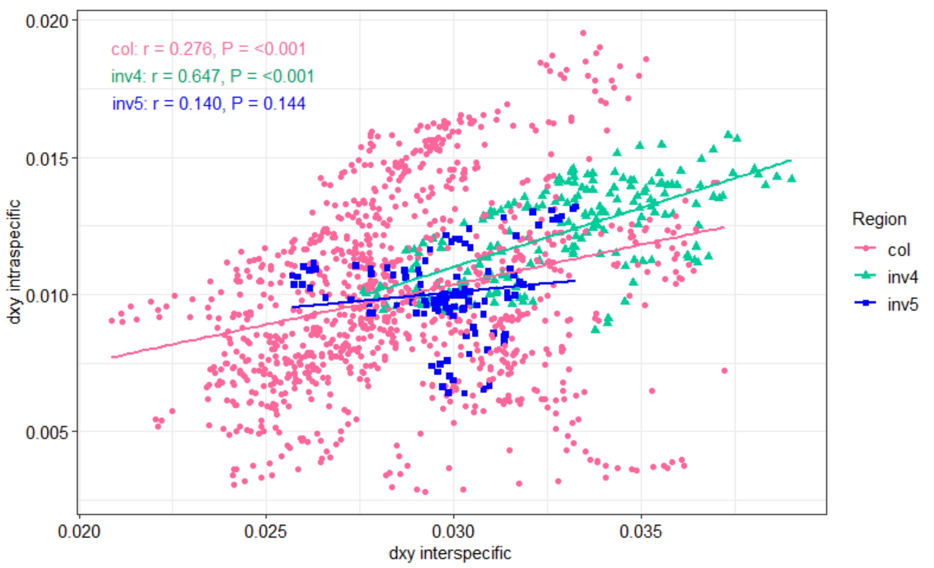

Figure S13. Pearson correlation analysis to test if correlation between interspecific and intraspecific $d_{xy}$ is greater for inverted autosomal regions compared to colinear background. Lines represent linear regressions.

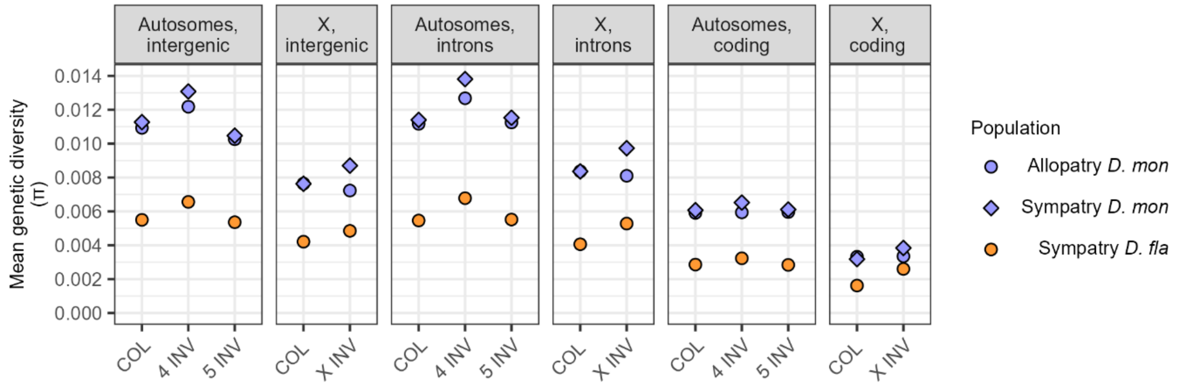

Figure S14. Mean genetic diversity (π) separately for intergenic, intronic and coding sequences, for autosomal and X chromosomal colinear (COL) and inverted (INV) chromosomal regions, as well as for *D. montana* originating from allopatric and sympatric populations and *D. flavomontana* originating from sympatric populations.

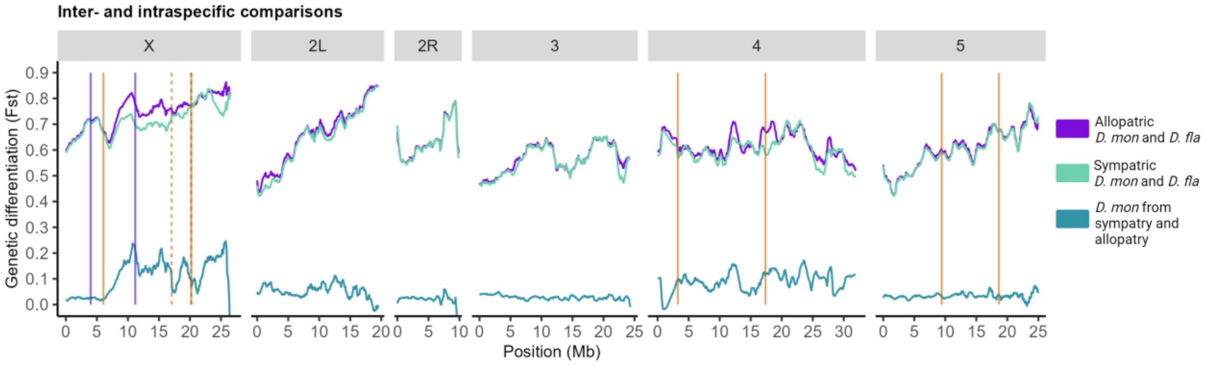

Figure S15. Genetic differentiation (measured as $F_{ST}$) across the genome (including intergenic regions) in sliding windows (window size 5,000 blocks, step size 500 blocks, block length 64b) for allopatric and sympatric comparisons of *D. montana* and *D. flavomontana* (interspecific), and for *D. montana* originating from allopatry and sympatry (intraspecific). Vertical lines represent breakpoints of different inversions (see Fig. S5): blue solid lines, and orange solid and dashed lines indicate alternatively fixed inversions of *D. montana* and *D. flavomontana*, respectively.

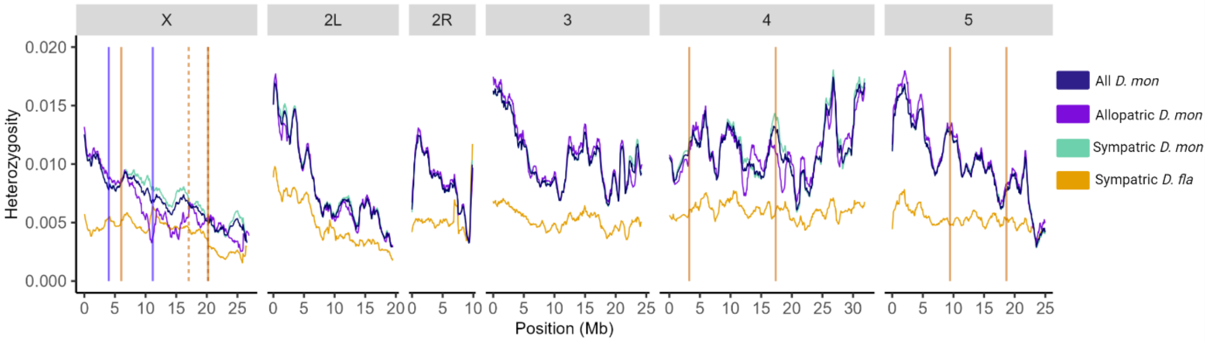

Figure S16. Heterozygosity across the genome (including intergenic regions) in sliding windows (window size 5,000 blocks, step size 500 blocks, block length 64b) separately for *D. montana* and *D. flavomontana* originating from sympatric and allopatric populations. Vertical lines represent breakpoints of different inversions (see Fig. S5): blue solid lines, and orange solid and dashed lines indicate alternatively fixed inversions of *D. montana* and *D. flavomontana*, respectively.

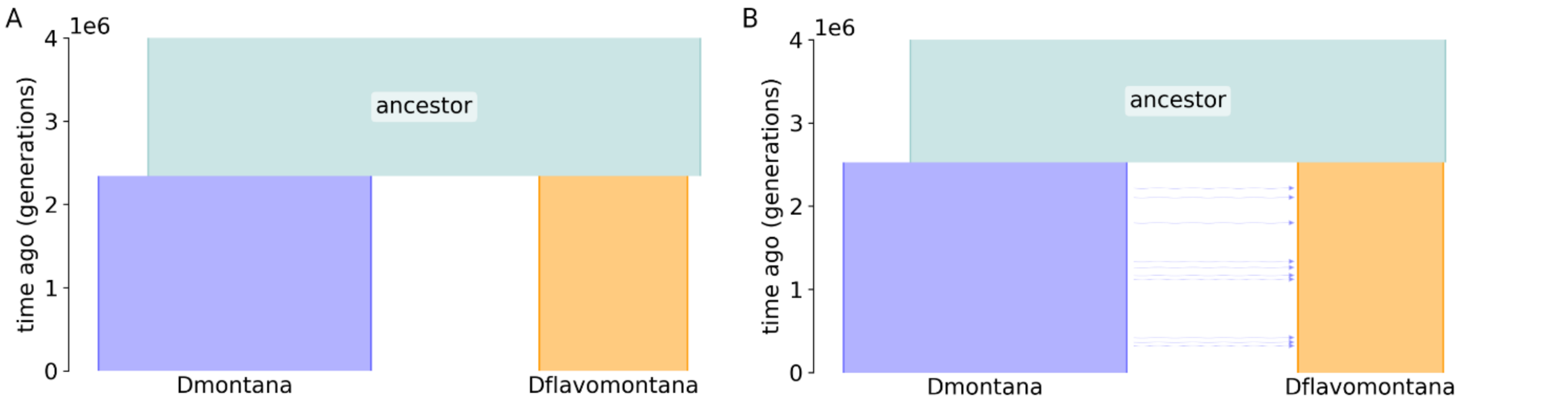

Figure S17. Sketches of the demographic models. The block widths indicate relative effective population sizes ($N_e$) of the ancestral population and its two descendants *D. montana* and *D. flavomontana*. (A) This model assumes no gene flow, i.e. strictly allopatric speciation (DIV). (B) The isolation with migration (IM) model allows unidirectional gene flow at rate *M* migrants per generation in this case from *D. montana* to *D. flavomontana*. Figures were produced with demesdraw (Gower et al. 2022).

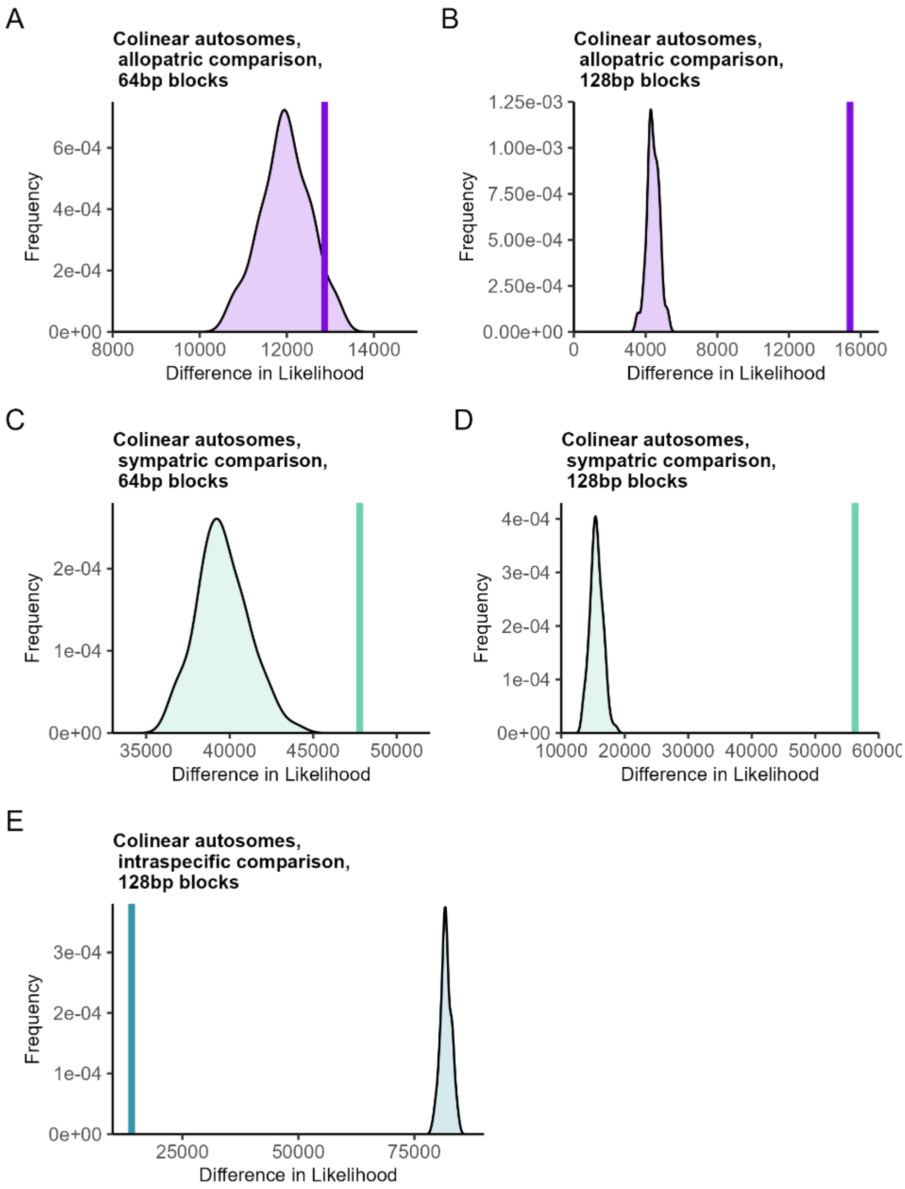

Figure S18. Datasets simulated under the strict divergence model (DIV) were fitted to both the DIV model (true model) and the IM model (wrong model), and the null distribution of ΔlnL was compared to the observed ΔlnL (vertical line). If the observed ΔlnL falls outside that null distribution, the IM model fits significantly better than the DIV model. The data were analysed using the *D. montana* genome as a reference. The analysis was performed for colinear autosomal regions of (A) allopatric comparison with block size 64b, (B) allopatric comparison with block size 128b, (C) sympatric comparison with block size 64b, (D) sympatric comparison with block size 128b, (E) intraspecific comparison with block size 128b.

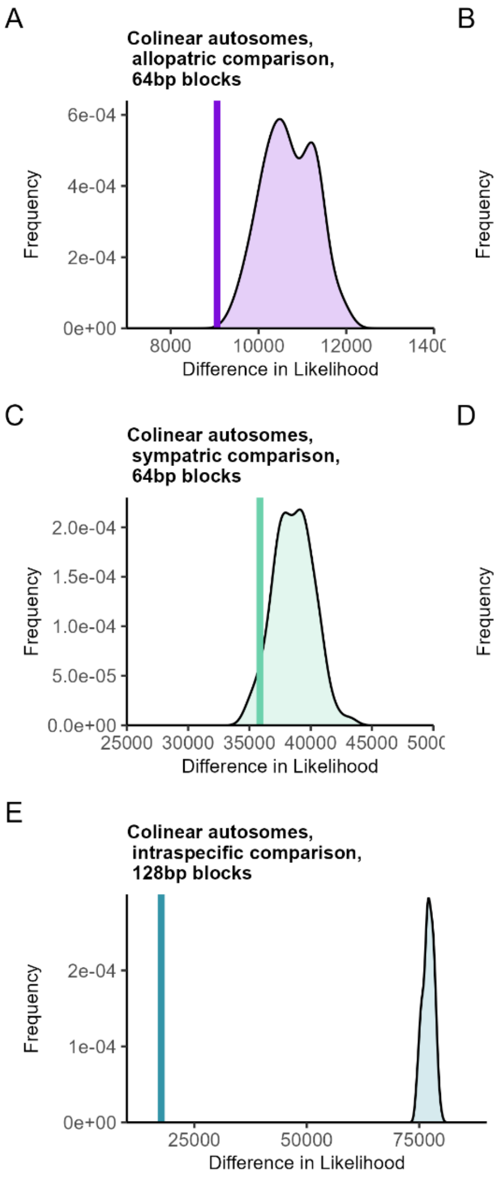

Figure S19. Datasets simulated under the strict divergence model (DIV) were fitted to both the DIV model (true model) and the IM model (wrong model), and the null distribution of ΔlnL was compared to the observed ΔlnL (vertical line). If the observed ΔlnL falls outside that null distribution, the IM model fits significantly better than the DIV model. The data were analysed using the *D. flavomontana* genome as a reference. The analysis was performed for colinear autosomal regions of (A) allopatric comparison with block size 64b, (B) allopatric comparison with block size 128b, (C) sympatric comparison with block size 64b, (D) sympatric comparison with block size 128b, (E) intraspecific comparison with block size 128b.

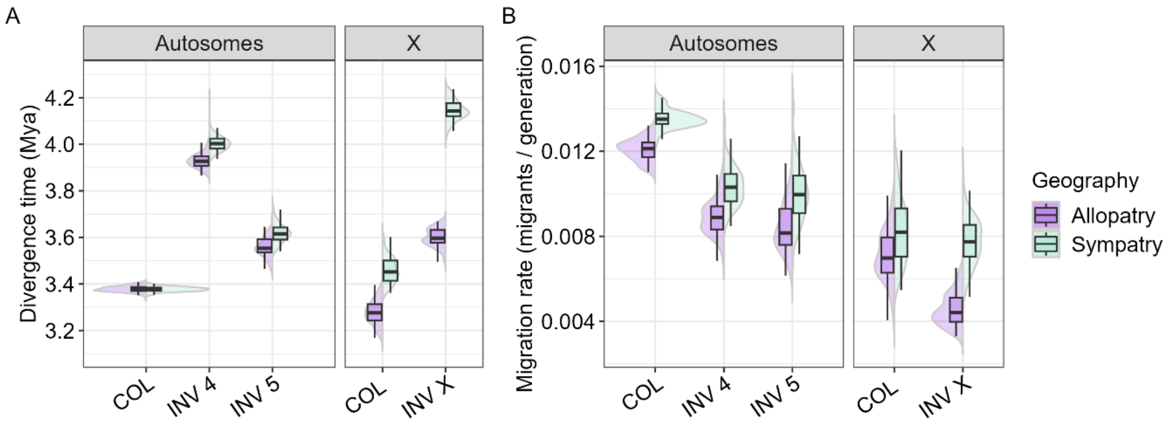

Figure S20. Estimates of (A) split times and (B) migration rates between *D. montana* and *D. flavomontana* for different chromosome partitions and for allopatric (purple) and sympatric (green) comparisons. Confidence intervals were estimated using 100 parametric bootstrap simulations (with recombination) under the best-fit IM model. Boxplot whiskers represent ±1.5×IQR; the distributions of estimates are shown as smoothed histograms.

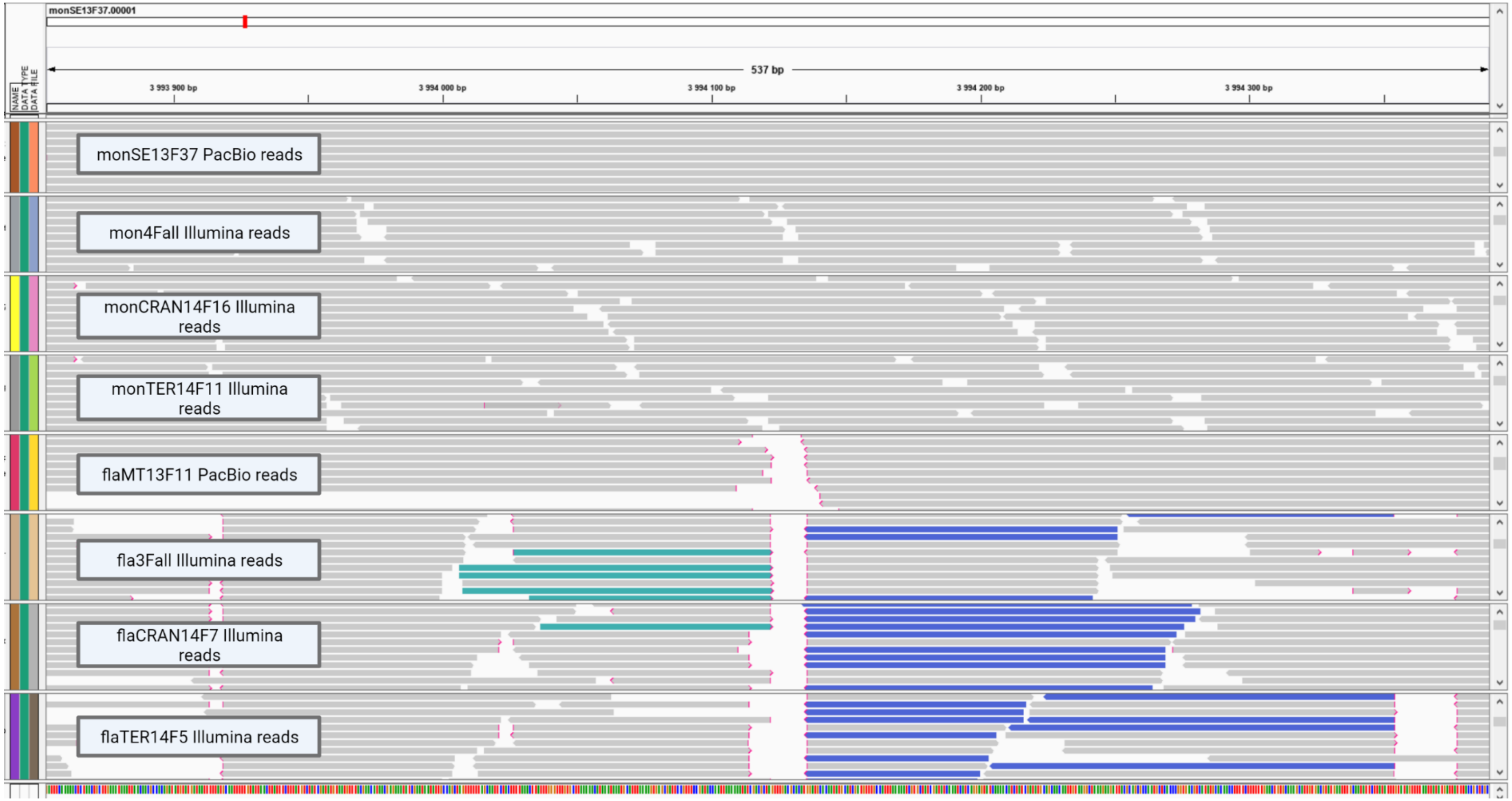

Figure S21. Example plot of an inversion breakpoint illustrated with Integrative Genomics Viewer (IGV) v2.8.0 (Thorvaldsdóttir et al. 2013) using both long read PacBio and short read Illumina data. Both long and short reads of *D. flavomontana* samples, mapped against *D. montana* genome, are split (red tip of the read) or in reversed orientation (blue and green reads) due to alternatively fixed inversion of the species.

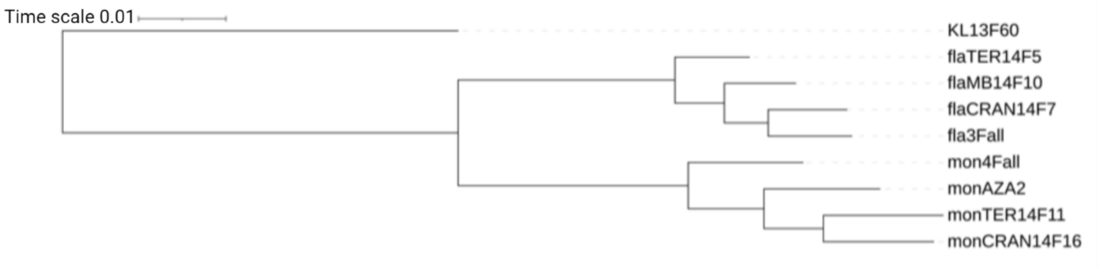

Figure S22. Phylogenetic tree produced by Orthofinder. Prefix “mon” and “fla” before a fly strain ID refer to *D. montana* and *D. flavomontana*, respectively. *D. littoralis* sample (KL13F60) was used as an outgroup in the tree. Details of the samples are given in Table S14.
